# Combined tumor and immune signals from genomes or transcriptomes predict outcomes of checkpoint inhibition in melanoma

**DOI:** 10.1101/2021.07.03.450733

**Authors:** Samuel S. Freeman, Moshe Sade-Feldman, Jaegil Kim, Chip Stewart, Anna L. K. Gonye, Arvind Ravi, Monica B. Arniella, Irena Gushterova, Thomas J. LaSalle, Emily M. Blaum, Keren Yizhak, Dennie T. Frederick, Tatyana Sharova, Ignaty Leshchiner, Liudmila Elagina, Oliver G. Spiro, Dimitri Livitz, Daniel Rosebrock, François Aguet, Jian Carrot-Zhang, Gavin Ha, Ziao Lin, Jonathan H. Chen, Michal Barzily-Rokni, Marc R. Hammond, Hans C. Vitzthum von Eckstaedt, Shauna M. Blackmon, Yunxin J. Jiao, Stacey Gabriel, Donald P. Lawrence, Lyn M. Duncan, Anat O. Stemmer-Rachamimov, Jennifer A. Wargo, Keith T. Flaherty, Ryan J. Sullivan, Genevieve M. Boland, Matthew Meyerson, Gad Getz, Nir Hacohen

**Affiliations:** Broad Institute of MIT and Harvard; Department of Biomedical Informatics, Harvard Medical School; Department of Medicine, Center for Cancer Research, Massachusetts General Hospital; Department of Medical Oncology, Dana-Farber Cancer Institute; Department of Cell Biology and Cancer Science, Rappaport Faculty of Medicine, Technion - Israel Institute of Technology; Division of Public Health Sciences, Fred Hutchinson Cancer Research Center; Department of Pathology, Massachusetts General Hospital; Departmnet of System Biology, Harvard Medical School; Department of Medical Oncology, Massachusetts General Hospital; Department of Surgical Oncology, University of Texas MD Anderson Cancer Center; Department of Surgery, Massachusetts General Hospital; Department of Genetics, Harvard Medical School; Department of Pathology, Harvard Medical School; Department of Medicine, Harvard Medical School

## Abstract

Cancer immunotherapy with checkpoint blockade (CPB) leads to improved outcomes in melanoma and other tumor types, but a majority of patients do not respond. High tumor mutation burden (TMB) and high levels of tumor-infiltrating T cells have been associated with response to immunotherapy, but integrative models to predict clinical benefit using DNA or RNA alone have not been comprehensively explored. We sequenced DNA and RNA from melanoma patients receiving CPB, and aggregated previously published data, yielding whole exome sequencing data for 189 patients and bulk RNA sequencing data for 178 patients. Using these datasets, we derived genomic and transcriptomic factors that predict overall survival (OS) and response to immunotherapy. Using whole-exome DNA data alone, we calculated T cell burden (TCB) and B cell burden (BCB) based on rearranged TCR/Ig DNA sequences and found that patients whose melanomas have high TMB together with either high TCB or high BCB survived longer and had higher response rates as compared to patients with either low TMB or TCB/BCB. Next, using bulk RNA-Seq data, differential expression analysis identified 83 genes associated with high or low OS. By combining pairs of immune-expressed genes with tumor-expressed genes, we identified three gene pairs associated with response and survival (Bonferroni *P*<0.05). All three gene pair models were validated in an independent cohort (n=180) (Bonferroni *P*<0.05). The best performing gene pair model included the lymphocyte-expressed *MAP4K1* (Mitogen-Activated Protein Kinase Kinase Kinase Kinase 1) combined with the transcription factor *TBX3* (T-Box Transcription Factor 3) which is overexpressed in poorly differentiated melanomas. We conclude that RNA-based (*MAP4K1*&*TBX3*) or DNA-based (TCB&TMB) models combining immune and tumor measures improve predictions of outcome after checkpoint blockade in melanoma.

## Introduction

Why only a subset of cancer patients respond to checkpoint blockade therapies, including anti–CTLA-4, anti–PD-1 monotherapies or the combination of both, is still unclear. For example, while patients with microsatellite instability (MSI), which have high indel and mutation burden, have improved response rates compared to non-MSI cases of the same tumor type, the predictive value of tumor mutation burden (TMB) is not always strong^1–4^. Furthermore, although T cells are essential for responses, their presence alone does not determine whether patients will benefit from checkpoint therapy^4–10^. Studies of acquired resistance have discovered rare mutations associated with lack of response^11–15^, but these do not explain the majority of resistant cases.

In recent years many studies have identified different mechanisms that could explain either response or resistance to CPB therapies. Initially, tumor mutational burden (TMB) was identified as a predictor of melanoma response to checkpoint blockade in melanoma ^16, 17^, A subsequent study in larger cohorts demonstrated a significant association between TMB as a continuous variable and overall survival for multiple tumor types^2^, however the Samstein et al. study did not identify a significant relationship between high TMB (defined either as above the 20th or 30th percentile) and overall survival in melanoma^2, 18^. Nonetheless, not all studies have observed a strong association between high tumor mutational burden and immunotherapy response, and in all cases there is an overlap in TMB between patients who respond or progress. Additionally, some studies have identified sun exposure or melanoma subtype as a potential factor confounding the association of melanoma TMB with immunotherapy response^4, 19^, but these results were not validated when analyzing independent cohorts. Additionally, inactivating mutations in *SERPINB3*/*SERPINB4* were found to be associated with immunotherapy outcome in two cohorts^15^, but these findings were not reproduced in a meta-analysis of other cohorts^20^. Another recent meta-analysis by Litchfeld et al. found that many immunotherapy predictors were not significant when analyzing multiple cohorts together, and even fewer predictors were significant when tested in independent cohorts^20^. The authors found that by combining TMB with gene expression signatures^21, 22^ or other immune features (e.g. CXCL9)^20^, they were able to predict immunotherapy outcomes in a meta-analysis cohort and validate their findings in an independent cohort. Thus, meta-analysis of larger cohort studies using independent cohorts for validation is therefore essential for identifying novel robust features underlying response to checkpoint inhibition.

While most studies have been focused on analyzing either the malignant^2–4, 6, 11, 12, 23–27^ or microenvironmental^5, 7, 28–33^ features and their association with clinical outcomes, recent studies using integrative models have improved our ability to distinguish between patients who are more likely to benefit from checkpoint blockade therapies. For example, integrating a T-cell inflamed gene expression profile (GEP) composed of 28 immune related genes with DNA-based TMB improved strasifications of HNSCC and melanoma patients treated with anti-PD1 monotherapy^21, 22^. Additionally, DNA-based sequencing of either tumor or cfDNA combined with immunohistochemical staining for PD-L1 improved prediction of response in NSCLC patients treated with anti-PD-L1^34^ or a combination of anti-CTLA4 and anti-PD1 therapies^35^. One major limitation of these approaches is the usage of multiple assays that require large samples and multiple nucleic acid isolation and preservation techniques (e.g., RNA and DNA), that in many cases are not available at different sites.

To address some of the mentioned limitations, we analyzed tumor exomes and transcriptomes from patients with melanoma receiving checkpoint blockade therapy, using two independent large cohorts, and derived several DNA or RNA-based pre-treatment features that are predictive of survival and response to therapy. First, combining TMB with quantification of T and B cell abundance using only WES data identified a subgroup of patients with high immune infiltration and high TMB that are more likely to benefit from immunotherapy compared to other patients. Next, using transcriptomic data, we identified the combination of the transcription factor *TBX3,* expressed in poorly differentiated melanomas, with *MAP4K1*, expressed in lymphocytes and dendritic cells, to be predictive of overall survival and response in both the primary and secondary meta-analysis cohorts. Overall, the present study serves as a resource for investigating immunotherapy outcome predictors and improves our knowledge of potential mechanisms underlying response or resistance to immunotherapy.

### Bulk DNA and RNA meta-analysis in melanoma patients treated with checkpoint inhibitors

To better understand the factors that predict response and/or survival in the context of immunotherapy, we sequenced DNA and RNA from melanoma samples before and after checkpoint blockade. We performed whole-exome sequencing (WES) of 109 samples from 56 patients (of which 37 patients had matched pre/post-treatment biopsies) and bulk RNA sequencing (RNA-seq) of 88 samples from 48 patients. These data were aggregated with published WES^12, 17, 36^ and bulk RNA-seq^6, 17, 37^ (Supplementary Fig. 1 and Supplementary Table 1- 2). In total, we analyzed 258 DNA WES samples from 189 patients (52 with matched pre/post-treatment samples) and 261 bulk RNA-seq samples from 178 patients (68 with matched pre/post-treatment samples; Fig. 1a). Overall, 59 patients had both DNA WES and RNA-seq data from a pre-treatment time point and 154 patients had RNA-Seq from a pre-treatment time point. Throughout the study, we evaluated the response and overall survival for all patients, except for one patient from the Hugo at el. study for whom survival data was missing. For the MGH cohort, we determined response based on a combination of radiographic measurements routinely performed on all patients with clinical evaluations (range 4-12 weeks after initiation of treatment), and we defined overall survival from initiation of therapy until the date of death or last follow-up. For the published exome and RNA-Seq cohorts, we used their overall survival data and their definitions for classification of response (Methods).

**Figure 1.**
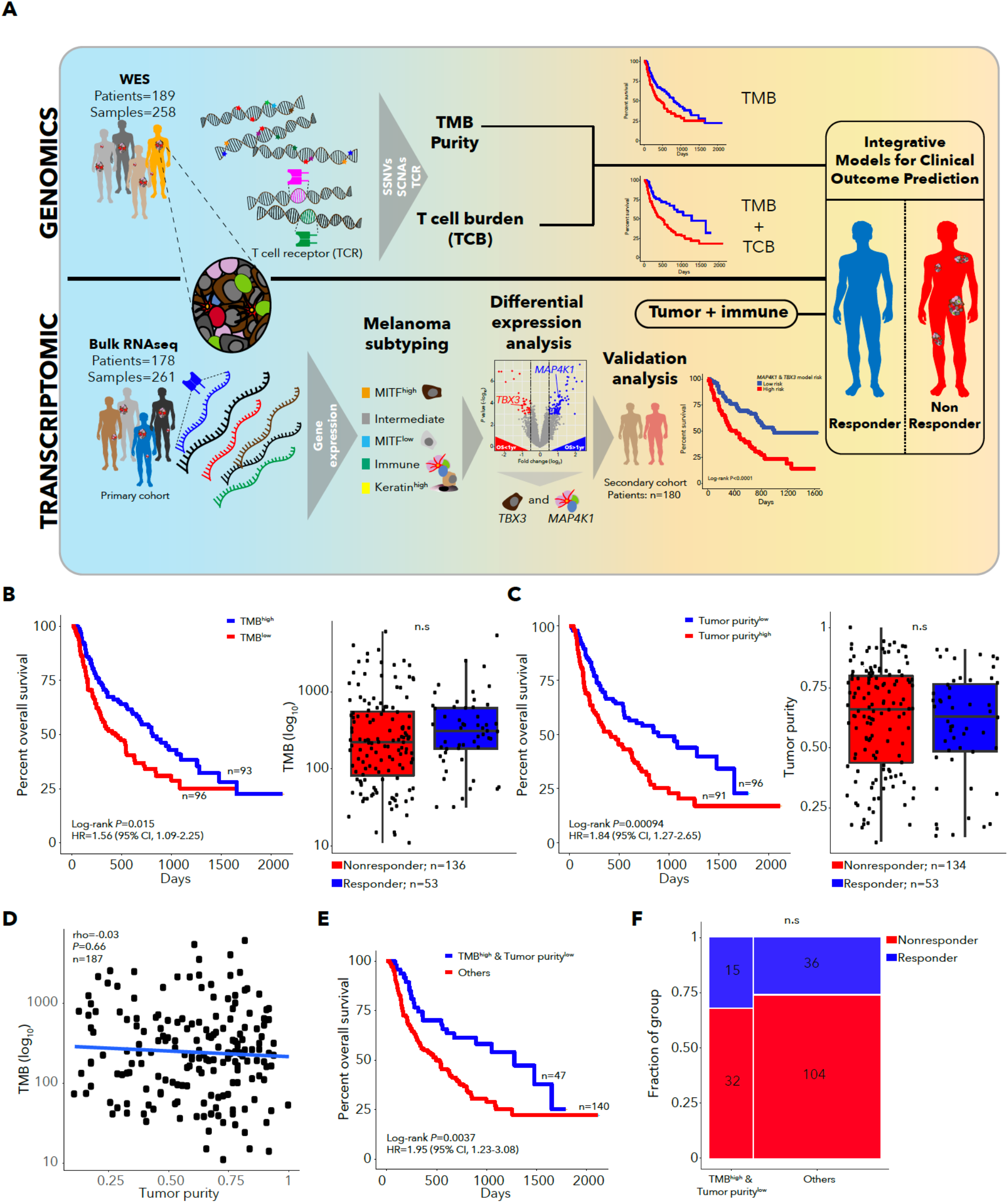
**A.** Study overview. **B.** Kaplan-Meier survival curve for patients with high (above median) and low (below median) TMB (left), and TMB for responders and non-responders (right). **C.** Kaplan-Meier survival for patients with high (above median) and low (below median) tumor purity (left), and tumor purity for responders and non-responders (right). **D.** Correlation between TMB and tumor purity. **E.** Kaplan-Meier survival curve for the TMB high, low tumor purity subgroup vs. other patients. **F.** Response rate for the TMB high, low tumor purity subgroup vs. other patients.

### DNA-based analysis demonstrates that combining TMB with DNA-based measures of immune infiltration improves predictions of immunotherapy outcomes

Our cohort-level analysis of WES data (*n*=189 patients, Supplementary Fig. 2) identified significantly mutated genes, somatic copy number alterations (SCNAs) and mutation signatures (Supplementary Fig. 3-4 and Supplementary Table 3-12), similar to previous studies^38^. As others observed^6, 16, 17^, we found that patients with TMB above median (TMB-high) or TMB above 10 mutations/Mb had increased survival after immunotherapy (TMB-high log-rank *P*=0.015, HR 1.56, Fig. 1b, TMB>10 *P*=0.017, HR 1.56, Supplementary Fig. 5). However, TMB was not significantly higher in responders than non-responders (Wilcoxon *P*=0.13, Fig. 1b). Neoantigen burden and clonal TMB highly correlated with TMB (rho=0.99 and 0.97 respectively) but did not provide additional predictive power over TMB (Supplementary Fig. 5). Though some studies have demonstrated that aneuploidy is associated with poor outcomes after immunotherapy^36, 39^, we found that tumor ploidy was not associated with survival (log-rank *P*=0.35). Additionally, survival models using the mutation status of single genes did not identify associations that passed multiple hypothesis correction (Supplementary Fig. 6), likely due to lack of power^3^. We identified somatic mutations in *B2M* that were present in post-treatment but not pre-treatment tumors in multiple patients^11, 12^, but did not identify any novel genes with statistically significant mutations that occurred only in post-treatment tumors (Supplementary Fig. 6-7). Consistent with prior reports of the association between low tumor purity and melanoma immunotherapy outcomes due to higher immune infiltration^4^, we found that tumor purity below median was associated with survival (log-rank *P*=0.00094), but not response] (Fig. 1c). Since tumor purity and TMB were not correlated (rho=-0.03, Fig. 1d), we combined these factors and found that the subgroup with high TMB and low tumor purity had longer survival (log-rank *P*=0.0037, HR 1.95, Fig. 1e, Supplementary Fig. 5), but was not associated with response (Fig. 1f). While single-gene analyses did not identify important features, the combined analysis of tumor purity and TMB suggests that combining tumor and immune features may improve predictive models.

Since T and B cell infiltration based on bulk RNA-seq analysis has been associated with response to immunotherapy and are inversely correlated with tumor purity^6, 7, 17, 22, 31, 32, 36, 40^, we next considered the predictive value of T and B cells quantified using rearranged T cell receptor (TCR) and immunoglobulin (Ig) sequences, respectively, from RNA. Targeted TCR and Immunoglobulin repertoire sequencing is required for full analysis of TCR and Ig clonotype diversity, but analysis of TCR and Ig rearrangements in bulk RNA-Seq or exome data can quantify T or B cell infiltration levels. High TCR or Ig read counts in pre-treatment RNA-seq samples (n=154) (TCR_RNA_ and Ig_RNA_ respectively) were associated with survival and response (Fig. 2a-b and Supplementary Table 13). Furthermore, TCR_RNA_ and Ig_RNA_ were highly correlated with expression of known T or B cell markers (Fig. 2c-d). Thus, we created an RNA-based metric of T or B cell burden (TCB_RNA_ or BCB_RNA_) using the number of rearranged TCR or Ig reads, respectively (Methods), and found them to be relatively consistent across cohorts and correlated with each other (rho=0.67; Supplementary Fig. 8). The patient subgroup with high TCB_RNA_ and BCB_RNA_ had increased overall survival but no association with response (Fig. 2e, Supplementary Fig. 8).

**Figure 2.**
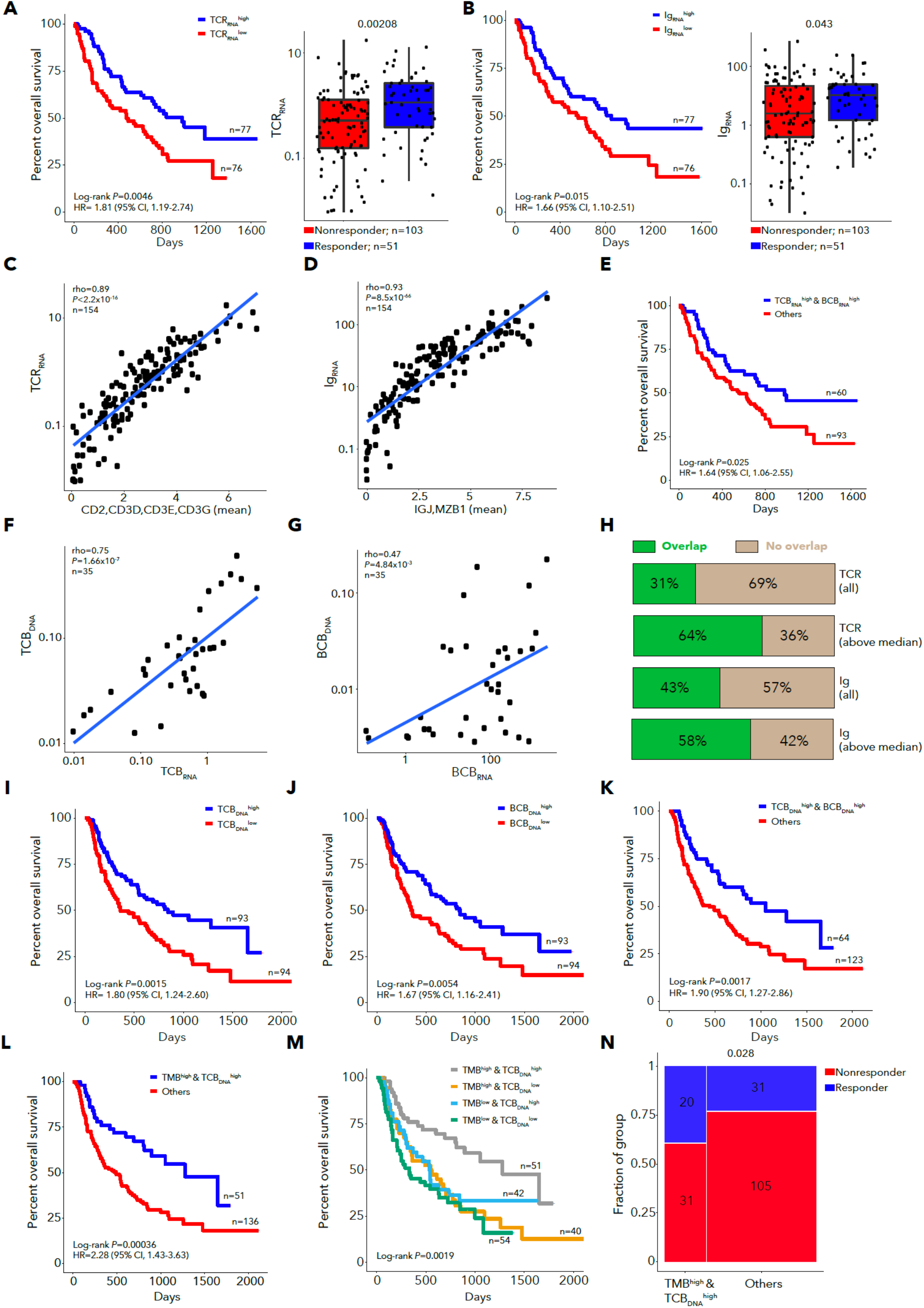
**A**. Kaplan-Meier survival curve for patients with high and low TCR_RNA_ (left panel) and TCR_RNA_ for responders and non-responders (right panel). **B.** Kaplan-Meier survival curve for patients with high and low Ig_RNA_ (left panel), and Ig_RNA_ for responders and non-responders (right panel). **C.** Correlation between TCR_RNA_ and T cell gene expression. **D.** Correlation between Ig_RNA_ and B cell gene expression. **E.** Kaplan-Meier survival curve for the T cell burden (TCB) in RNA (TCB_RNA_) high, B cell burden (BCB) in RNA (BCB_RNA_) high subgroup vs. other patients. **F.** Correlation between TCB_RNA_ and TCB_DNA_, for patients with DNA and RNA extracted from the same location in the tumor. **G.** Correlation between BCB_RNA_ and BCB_DNA_, for patients with DNA and RNA extracted from the same location in the tumor. **H.** Fraction of cases with TCR or Ig CDR3 clonotypes shared between RNA and DNA, for patients with DNA and RNA extracted from the same location in the tumor. **I.** Kaplan-Meier survival curve for patients with high and low TCB_DNA_. **J.** Kaplan-Meier survival curve for patients with high and low BCB_DNA_. **K.** Kaplan-Meier survival curve for the TCB_DNA_ high, BCB_DNA_ high subgroup vs. other patients. **L.** Kaplan-Meier survival curve for the TMB high, TCB_DNA_ high subgroup vs. other patients. **M.** Kaplan-Meier survival curve for all TMB and TCB_DNA_ subgroups. **N.** Response rate for the TMB high, TCB_DNA_ high subgroup vs. other patients.

To extend the association between TCB/BCB and outcome to a larger cohort, we generated equivalent metrics based on rearrangements in DNA, TCB_DNA_ and BCB_DNA_ (Supplementary Table 14). Since we did not perform targeted TCR sequencing but rather used WES data, we first tested whether the RNA and DNA-based metrics were correlated using 35 cases with DNA and RNA extracted from the same area of the tumor (Fig. 2f-h and Supplementary Table 15-16). As the levels of TCB_DNA_ and BCB_DNA_ differed between cohorts, we classified samples as above or below median within each cohort, and found that these TCB_DNA_ and BCB_DNA_ classes associated with survival, as did their combination (Fig. 2i-k, Supplementary Fig. 9). Further validating the consistency of TCB/BCB, we detected shared TCR and Ig CDR3 sequences across DNA and RNA (Fig. 2g), with increased sharing in samples with higher TCB_DNA_ or BCB_DNA_. These results demonstrate that lymphocyte infiltration can be quantified using rearranged TCR or Ig reads from tumor exomes alone and is associated with survival.

Next, we combined tumor and immune features using DNA-based data in order to improve outcome models. Since TMB did not correlate with TCB_DNA_ (rho=0.03, Supplementary Fig. 10), we tested a model combining both and found that patients with high TMB and high TCB_DNA_ survived longer (*P*=3.6×10^-4^, HR=2.28, Fig. 2l,m) and had a higher response rate (*P*=0.028, OR=2.18, Fig. 2n). This combined model was superior to models incorporating TMB or TCB_DNA_ alone (Likelihood ratio test *P*=0.0036 and 0.038, respectively). Similarly, the subgroup with high TMB and high BCB_DNA_ was associated with longer survival and higher response rates (log-rank *P*=1.6×10^-3^, Fisher *P*=0.021, Supplementary Fig. 10c-d). However, a model with all three factors (TMB, TCB_DNA_ and BCB_DNA_) did not provide additional predictive value (Supplementary Fig. 10f-i). We also reanalyzed stage III/IV melanomas from The Cancer Genome Atlas (TCGA, Supplementary Fig. 11 and Supplementary Table 17-18), a subset of which were treated with checkpoint blockade or other immunotherapy, and found that the TMB high, TCB_DNA_ high subgroup had increased survival (log-rank *P*=0.025, Supplementary Fig. 11d-k). Thus, through combining tumor and immune features by quantifying TMB and TCB_DNA_ from DNA sequencing alone, we were able to identify patients with a higher chance of benefiting from checkpoint blockade immunotherapies.

### Tumor RNA-Seq analysis identifies melanoma subtypes associated with immunotherapy survival

As previous studies have demonstrated expansion of T cell clones after immunotherapy^5, 41^, we next analyzed lymphocyte expansion in paired pre-treatment and post-treatment samples using TCB and BCB. We found that TCB measured in RNA and DNA increased from pre-treatment biopsies to post treatment biopsies, but BCB did not increase in either RNA or DNA (Supplementary Figure 12a-d and Supplementary Table 19). Recent work has shown that pre-treatment immune infiltration is associated with response in patients receiving anti-CTLA-4 prior to anti-PD-1 but not in patients receiving anti-PD-1 alone with no prior anti- CTLA-4^4^. We, therefore, tested whether TCB increases were specific to patients with no prior anti-CTLA-4 therapy. Indeed, we found that TCB_RNA_ increased in patients who had not received prior anti-CTLA-4, but not in those who were treated with prior anti-CTLA-4 (Supplementary Fig. 12e-h). However, TCB_DNA_ increased irrespective of prior CTLA-4 therapy, though there were few DNA samples from patients with prior CTLA-4 (Supplementary Fig. 12e-h). BCB did not significantly increase in DNA or RNA irrespective of prior CTLA-4 treatment. Finally, we found that an increase in TCB or BCB from pre-treatment to post-treatment biopsies was not associated with response or survival (Supplementary Fig. 13a-h), however, there is a bias since most post-treatment biopsies were from resistant lesions. Our results are consistent with the association of levels of both T and B cells in pre-treatment samples with outcome but suggest that checkpoint blockade may induce T cell, but not B cell expansion.

**Figure 3.**
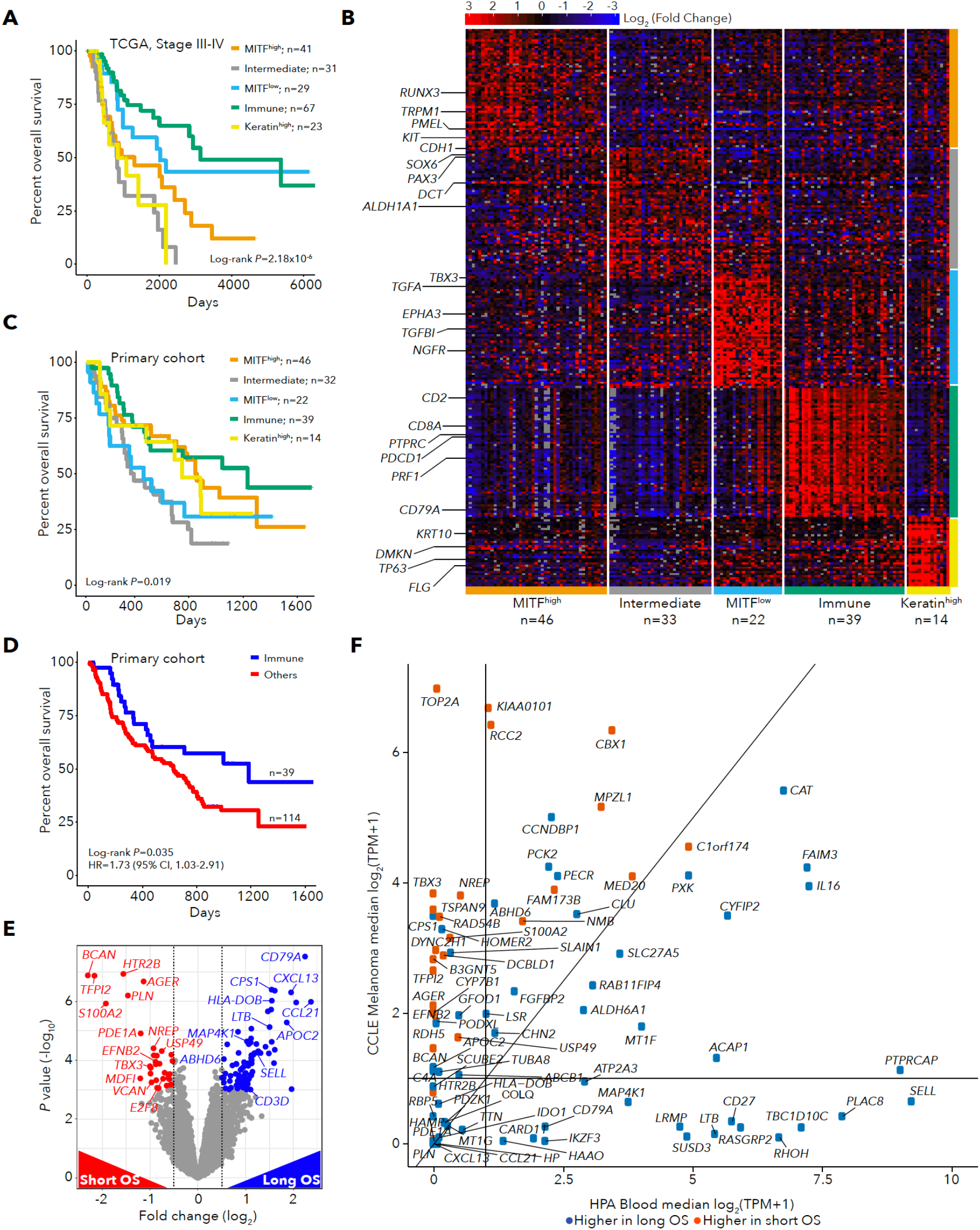
**A.** Kaplan-Meier survival curve by expression subtype for TCGA melanoma stage III/IV patients. **B.** Heatmap of marker gene expression for pre-immunotherapy (primary cohort; n=154) patients grouped by subtype. **C.** Kaplan-Meier survival curve by subtype for pre-immunotherapy (primary cohort; n=153) patients. **D.** Kaplan-Meier survival curve for immune subtype vs. other patients in the primary cohort. **E.** Differential expression between patients with Overall Survival (OS) > 1 year (long OS) and patients with OS < 1 year (short OS) in the primary cohort. **F.** Expression of differentially expressed genes in melanoma CCLE cell lines and Human Protein Atlas blood cell types.

Expression-based cancer subtypes have been linked with patient survival both with or without immunotherapy^10, 38, 42–44^. Using bulk RNA sequencing data from 469 TCGA melanoma specimens (101 primary and 368 metastatic biopsies^38^), we identified 5 robust tumor subtypes with Bayesian non-negative matrix factorization (NMF) clustering^45^ (Supplementary Fig. 14 and Supplementary Table 20-21). As expected from previous studies^6, 17^, one subtype was marked by high levels of immune infiltrate (“Immune”), and a second by high levels of keratin expression (“Keratin-high”), likely due to keratinocytes. The other three subtypes were associated with tumor-intrinsic properties related to the degree of melanocyte differentiation. Two subtypes were classified by expression of the melanocyte-inducing transcription factor (*MITF*) gene (“*MITF*-low” and “*MITF*-high”), and a third by intermediate melanocyte differentiation (“Intermediate”) (Supplementary Fig. 15). The *MITF*-low and Immune subtypes were concordant with two subtypes previously identified in TCGA^38^ (Supplementary Fig 14). The *MITF*-high, Intermediate, and *MITF*-low subtypes were closely related to differentiation states identified in melanoma cell lines^46^ (Supplementary Fig. 14d). The poorly differentiated *MITF*-low subtype resembles neural crest stem cells and is associated with resistance to targeted therapies^47, 48^ and immunotherapies^37^. TCB_RNA_ and BCB_RNA_ were higher in Immune subtype tumors (Supplementary Fig. 15). Using TCGA data, the five subtypes were strongly associated with survival for all (log-rank *P*=2.00 x 10^-10^, Supplementary Fig. 15g) and for stage III/IV melanoma patients (*P*=2.18 x 10^-6^, Fig. 3a). When we analyzed RNA-seq data from pre-immunotherapy patients (n=154), after batch-effects correction between cohorts (Supplementary Fig. 16a-g and Supplementary Table 22-23), we found that tumor subtypes were associated with post-immunotherapy survival (log-rank *P*=0.019, Fig. 3b-c), but not response (Supplementary Fig. 16h), with the Immune subtype associated with the longest survival (Fisher *P*=0.035, HR=1.73, Fig 3d).

### Identification and validation of gene-pair models combining tumor and immune genes to predict immunotherapy survival and response

To further pinpoint gene expression markers of outcome, we determined which genes were differentially expressed between patients with overall survival (OS) >1 year (long OS) and patients with OS < 1 year (short OS), irrespective of subtype (*n*=154 samples; Fig. 3e and Supplementary Table 24; median TPM>1, DESeq q<0.05). We identified 83 genes differentially expressed between long and short OS patients (55 overexpressed in long OS patients, 28 overexpressed in short OS patients). Genes associated with better outcomes included both T and B lymphocyte expressed genes (*CD3E*, *CD3G, LTB*, *SELL*, *SLAMF6*, *CD52, CD79A, CXCL13*, *MAP4K1*), and genes associated with poor outcomes included multiple tumor-expressed genes (*TBX3*, *EFNB2*, *NREP*, *S100A2*, *AGER*). We also identified 101 genes differentially expressed between responders and non-responders (q<0.05), and they significantly overlapped the 83 genes associated with OS (overlap 29/101, *P*=8.16×10^-41^, Supplementary Fig. 17). We next analyzed the 55 genes associated with long OS and found that most were expressed in multiple immune cell types, including lymphocytes and memory CD8 T cells, which are known to be important for anti-tumor immune responses^7^ (Fig. 3f; Supplementary Figs. 17-18). In contrast, the 28 genes overexpressed in patients with short OS were highly expressed in melanoma cell lines, with the highest expression in the *MITF*-low subtype (Fig. 3f; Supplementary Fig. 18d-e). We found similar tumor and immune expression patterns for genes differentially expressed between responders and non-responders (Supplementary Fig 19 and Supplementary Table 25-27).

Since previous studies successfully combined tumor and immune features to improve the prediction of immunotherapy outcomes (^21, 22, 34, 35^) and we found that the combination of TMB with TCB_DNA_ improved predictions of outcome, we next sought to create models using gene pairs by combining an immune-associated gene with a tumor-associated gene. We tested all pairwise combinations of the 83 OS differentially expressed genes as predictors of survival and response (Supplementary Table 28). Additionally, we tested a metagene pair model in which we averaged the normalized expression of the 55 long OS genes as one metagene and the 28 short OS genes as the other metagene. This metagene pair model was highly predictive of response and survival (Supplementary Fig. 20). Considering all gene pair models, we found that models based on pairs of short OS genes were significantly worse than either models based on pairs of long OS genes or pairs with one long OS and one short OS gene (Supplementary Fig. 20c-e and Supplementary table 29-31).

After testing all pairwise models of genes differentially expressed between patients with long or short OS, we identified 3 gene pairs which were significantly associated with survival and with response (Bonferroni-corrected *P*<0.05 for both models, Fig. 4a, Supplementary Fig. 20f-i). The three significant pairs were *MAP4K1*&*TBX3*, *MAP4K1*&*AGER* and the metagene pair model. *MAP4K1* (also known as Hematopoietic Progenitor Kinase 1) is expressed in multiple immune cell types including T and B lymphocytes as well as dendritic cells^49, 50^. In contrast, *TBX3, AGER* and the short OS metagene are most highly expressed in the dedifferentiated MITF-low melanoma subtype (Supplementary Fig. 21). Next, we compared the performance of the three gene pair models to six published models of immunotherapy outcomes: CD274 (PD-L1) expression, GEP, CYT, IMPRES, TIDE and MHC II^4, 22, 51–53^. We also evaluated TCB_RNA_ as a predictor of immunotherapy outcomes. When we computed values for each predictor for each patient and clustered these values, we identified a cluster of patients with high immune infiltration and high values for multiple immune-based models (Fig. 4b and Supplementary Table 32). Additionally, we found clusters of patients with low levels of immune infiltration and high values for tumor-associated predictors, and many of these tumors were classified in the Intermediate and MITF low subtypes (Fig. 4b). We found that the three gene pair models outperformed the previous models in predictions of response and survival (Fig. 4c-d, Supplementary Fig. 21d-f). To assess the robustness of the three gene pair models, we performed a cross-validation analysis, and we found that: 1) increasing the training set size increased the number of gene pair models discovered, 2) increasing the training set size increased the robustness of the long OS metagene but the short OS metagene was still variable, and 3) the top gene pair models were repeatedly discovered in training data subsets, but were very rarely significant in the held out validation sets, supporting the need for larger datasets (Supplementary Fig. 22 and Supplementary Table 33-34).

**Figure 4.**
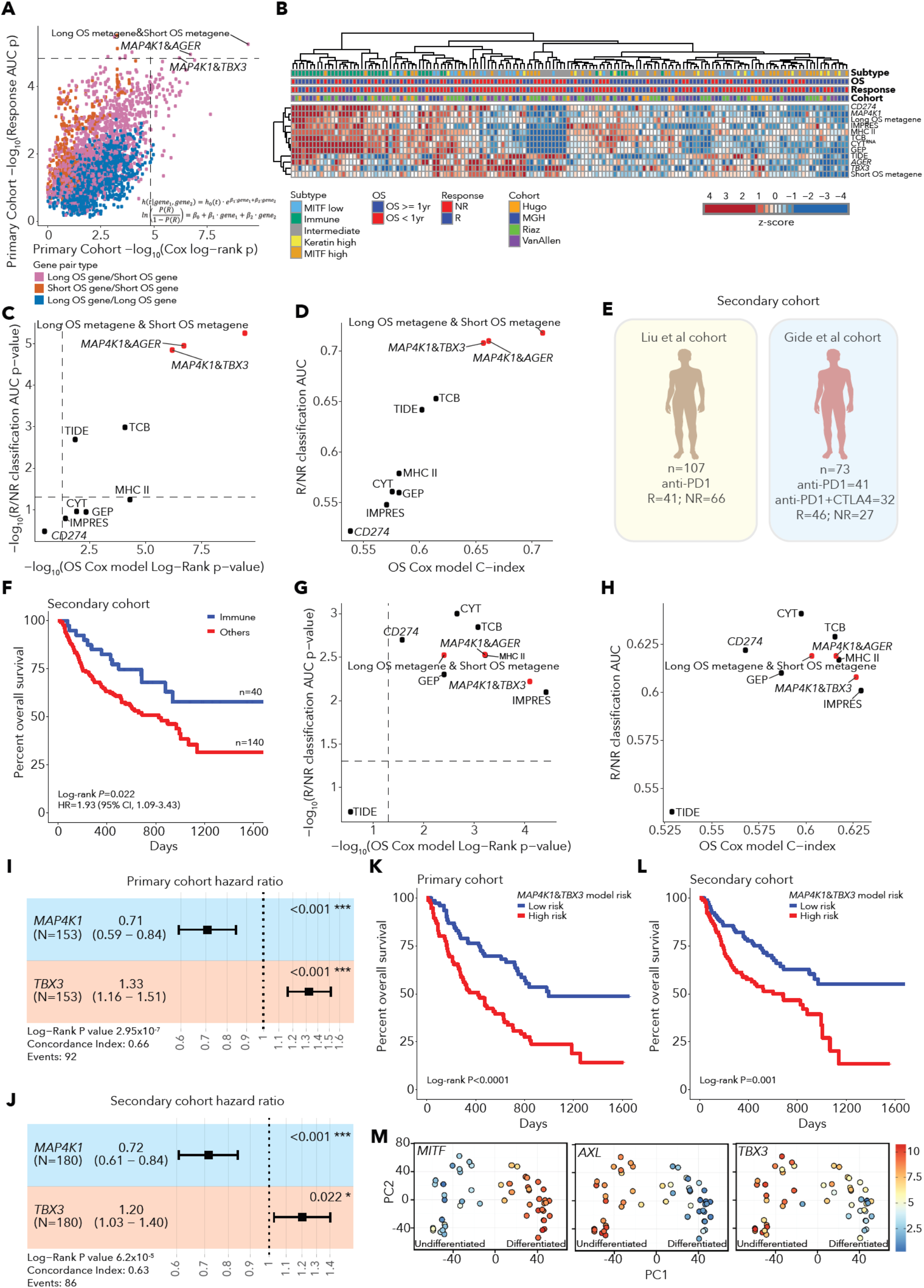
**A.** Performance of gene pair models in predictions of survival (Cox proportional hazards model Log-Rank P value) and response (Logistic regression AUC p value) in the primary cohort identifies three gene pairs models with Bonferroni p<0.05. **B.** Heatmap of z-scored values for immunotherapy predictive models and top gene pairs. **C.** Performance of pairwise gene models in comparison to previous immunotherapy predictive models in significance of predictions of response and survival in the primary cohort. **D.** Performance of pairwise gene models in comparison to previous immunotherapy predictive models in effect size of predictions of response and survival in the primary cohort. **E.** Schematic of independent secondary cohort. **F.** Kaplan- Meier survival curve of Immune subtype patients in the secondary cohort. **G.** Performance of pairwise gene models in comparison to previous immunotherapy predictive models in significance of predictions of response and survival in the secondary cohort **H.** Performance of pairwise gene models in comparison to previous immunotherapy predictive models in effect size of predictions of response and survival in the secondary cohort. **I.** Forest plot of *MAP4K1*&*TBX3* survival model performance in the primary cohort. Error bars represent 95% confidence intervals for Cox model hazard ratio estimates. **J.** Forest plot of *MAP4K1*&*TBX3* survival model performance in the secondary cohort. Error bars represent 95% confidence intervals for Cox model hazard ratio estimates. **K.** Survival of patients stratified by high and low risk using *MAP4K1* and *TBX3* expression in the primary cohort. **L.** Survival of patients stratified by high and low risk using *MAP4K1* and *TBX3* expression in the secondary cohort. **M.** Expression of *MITF*, *AXL* and *TBX3* in melanoma cell lines shows that *TBX3* forms a gradient of expression across melanoma cell line dedifferentiation states.

In order to confirm our findings in an independent data set, we merged and analyzed two published cohorts of pre-treatment samples from melanoma patients receiving PD-1 or combination CTLA-4/PD-1^4, 54^ (n=180), which we refer to as the ‘secondary cohort’ (Fig. 4e). After batch-effects correction and subtype analysis (Supplementary Fig. 23 and Supplementary Table 35-37), we found that patients with Immune subtype tumors in the secondary cohort indeed had longer survival than other patients (log-rank *P*=0.022, HR=1.93, Fig. 4f, Supplementary Fig. 23). When we tested the performance of the three gene pair models, we found that all three models were validated in the secondary cohort with Bonferroni *P*<0.05 for predictions of survival and response. However, for predictions of response in the secondary cohort, their performance was statistically equivalent to that of previous models (DeLong’s test p>0.05), with AUC and C-index values between 0.6 and 0.7 (Fig 4g-h, Supplementary Fig. 24a-d and Supplementary Table 38- 40).

The top performing gene pair model in the secondary cohort was the model combining *MAP4K1* with *TBX3* (Fig. 4g-h). In Cox models incorporating *MAP4K1* and *TBX3*, both genes were significant predictors in both the initial cohort and the secondary cohort (Fig. 4 i-j). As expected, patient stratification into risk groups using *MAP4K1* and *TBX3* expression was also significantly associated with survival in both cohorts (Fig. 4 k-l, Supplementary Fig. 24). We found that treatment (PD-1 vs. combination CTLA-4/PD-1) was significantly associated with survival in the secondary cohort, but in a Cox survival model incorporating *MAP4K1*, *TBX3* and treatment, all three covariates were significant (Supplementary Fig. 24j-k). When we analyzed the performance of models in individual external cohorts, we found that their performance was variable, highlighting the importance of meta-analysis and large cohorts for identifying robust predictors (Supplementary Fig. 25a-j and Supplementary Table 41). The top gene pair models were able to predict outcomes, and the simplicity of these models points to potential biological connections between gene expression and outcome.

### *TBX3* is a marker of poorly differentiated melanomas

To better understand potential roles of *TBX3*, we analyzed multiple melanoma datasets. First, *TBX3* was expressed in most melanoma cell lines^46^ except for well-differentiated ones with high MITF expression (Fig. 4m), in concordance with *TBX3* expression levels increasing from the *MITF*-high (well-differentiated tumors) to the *MITF-*low subtypes in TGCA and pre-treatment tumors (Supplementary Fig. 21). Second, genes negatively correlated with *TBX3* expression in cell lines^55^ were enriched for pigmentation gene sets (Supplementary Fig. 26a and Supplementary Table 42), consistent with the low expression of *TBX3* in *MITF-*high tumors. Third, *TBX3* was expressed in malignant cells but rarely in non-tumor cells based on melanoma scRNA-seq data^48, 56^ and was more highly expressed in cells expressing the neural crest marker *NGFR* (Supplementary Fig. 26b-f), consistent with prior studies showing that NGFR+ malignant cells are resistant to checkpoint therapy^57^. Functional studies have shown that overexpressing *TBX3* in melanoma cells enhances tumor formation and invasion *in vivo*^58^. In summary, *TBX3* is a tumor-specific gene expressed in poorly-differentiated melanomas, and a model combining the lymphocyte-expressed gene *MAP4K1* with *TBX3* is associated with patient outcomes after immunotherapy. Overall, our results suggest that models combining a metric of immune infiltration with a tumor-derived metric associated with poor melanoma differentiation can predict melanoma immunotherapy outcomes.

## Discussion

By extracting several biological features from tumor DNA and RNA sequences, we uncovered factors that predict outcomes in melanoma patients treated with checkpoint blockade: (i) tumor purity and mutation burden, (ii) T/B cell infiltration combined with mutation burden based on DNA alone, (iii) transcriptome-wide subtyping of tumors, and (iv) expression of *MAP4K1* and *TBX3* (as well other gene pairs involving an immune-associated gene and a tumor-associated gene). In addition, although some features are correlated (e.g. tumor purity, immune infiltration, and T/B cell burden), specific combinations of uncorrelated features improved predictions of response and survival.

Our data show that integrative models measuring immune infiltration (*MAP4K1*) along with expression of tumor-associated genes (*TBX3)* result in predictive models of patient outcome. These simple models using gene pairs were able to perform as or more accurately than previous complex models and also provide insight into biological features associated with clinical outcome. This finding is consistent with previous studies which have found that multivariate models that combine multiple data types can often outperform simple models of post-immunotherapy survival (such as TMB, PD-L1 staining or T cell infiltration alone)^4, 22, 24, 34, 35^. We also show that high TMB (likely providing more neoantigens) combined with high TCB (indicating T cell response) can predict treatment outcome. Our results suggest that including TCR and Ig sequences in targeted sequencing panels, along with genes that allow TMB estimation, may be useful for prediction of outcome using a single DNA assay. Patients with multiple positive prognostic factors may be better served by PD-1/PD-L1 monotherapy, whereas those with more negative prognostic factors may benefit from more aggressive combinations of standard, targeted and immune therapies, but these predictions would need to be evaluated in prospective clinical trials.

Previous studies have demonstrated that including clinical variables and single-cell profiling can significantly improve predictive models of immunotherapy outcomes^4, 7^. We were able to identify predictive models and validate them in an independent cohort (n=180). While these models were more predictive than previously published models, their performance is still modest (with AUC and C-index of 0.6-0.7). Future studies will likely require deeper clinical, tumor and immune characterization of larger cohorts to further discover and test the utility of genetic and non-genetic predictors. To achieve this goal, the community will need to generate and share genomic, transcriptomic and outcome data from patients receiving immunotherapy.

## Methods

### Patient specimens and consent

All patients analyzed in this study (referred as the MGH cohort) provided written informed consent for the collection of tissue and matched normal blood samples for research and genomic profiling, as approved by the Dana-Farber/Harvard Cancer Center Institutional Review Board (DF/HCC Protocol 11-181) and UT MD Anderson Cancer Center (IRB LAB00-063 and 2012- 0846). All samples in this study are from patients with metastatic melanoma treated with checkpoint blockade therapy (Supplementary Table 1-2) at Massachusetts General Hospital (Boston, MA) and University of Texas MD Anderson Cancer Center (Houston, TX). Patient response status at initial restaging examination for the MGH cohort was defined based on a combination of radiographic measurements routinely performed on all patients with clinical evaluations (range 4-12 weeks after initiation of treatment), with patients achieving PR or CR for responders and SD or PD for nonresponders. Subsequent responses for biopsies taken during treatment were determined on the date of biopsy acquisition based on a combination of clinical and radiographic measurements. Overall survival (OS) was defined from the date of treatment initiation until the date of death or last follow-up.

### Clinical outcomes from previously published cohorts

For the analysis of outcomes of patients from previously published cohorts, we used the published overall survival and clinical response data. Overall survival data was reported in each study. For clinical response, we used the binary classification of clinical outcome from each individual study. For Hugo et al., patients with PR/CR by irRECIST criteria were classified as responders, and patients with PD by irRECIST criteria were classified as non-responders^37^. For Zaretsky et al., the 4 patients were classified as PR/CR by irRECIST criteria and were classified as responders^12^. For Roh et al., we used the same response definitions as stated in their paper: responders had either complete resolution of tumors or partial reduction of at least 30% in tumor size (based on imaging) or stable disease for over 6 months, and nonresponders had an increase of at least 20% in tumor size (based on imaging) or stable disease for under 6 months^36^. We used these definitions for each sample (anti-CTLA-4 response and anti-PD-1 response) from Roh et al. Table S1B. For Riaz et al., the authors separated patients with RECIST PD, SD and PR/CR, so we classified patients in the same manner as the MGH cohort with RECIST PR/CR as responders and patients with RECIST PD or SD as non-responders^6^. Additionally, for the two patients in the Riaz cohort (Pt76_pre and Pt23_pre) that died prior to disease assessment (after 10 and 52 days respectively), we included these patients as non-responders. For Van Allen et al., patients with durable clinical benefit (DCB, PR/CR or SD with OS>1 year) were classified as responders and patients with no durable clinical benefit (NDB, PD or SD with OS<1 year) were classified as non-responders^17^. For four patients in this cohort with no annotated RECIST response but very short overall survival (MEL-IPI_Pat157, MEL-IPI_Pat166, MEL-IPI_Pat168 and MEL-IPI_Pat175 with deaths after 86, 77, 67 and 90 days respectively), we included these patients as non-responders. Additionally, one patient with no RECIST annotation and an overall survival of 1326 days (MEL-IPI_Pat29) was included as a responder. For Liu et al., patients with RECIST PR, CR or listed as MR (mixed response) were classified as responders, and patients with RECIST PD or SD were classified as non-responders^4^. For Gide et al., patients with RECIST PR/CR or SD with PFS over 6 months were classified as responders, and patients with RECIST PD or SD with PFS under 6 months were classified as non-responders^54^. Overall survival was available for all patients (except for one patient from the Hugo at el. study for which survival data was missing).

### Whole exome sequencing (WES)

Whole exome sequencing was performed at the Broad Institute genomic platform on 109 tumor and matched normal blood samples from 56 patients using a protocol previously described^11^. Briefly, 150-500ng of gDNA was extracted using Qiagen AllPrep DNA/RNA Mini Kit (cat# 80204). DNA was fragmented using acoustic shearing followed by size selection to achieve library insert size distribution in the range of 300-650bp. Libraries for all samples were prepared using the Kapa HyperPrep kit, according to manufacturer’s specifications, followed by quantification and normalization using PicoGreen to ensure equal concentration. Adaptor ligation was performed using the TrueSeq DNA exome kit from Illumina according to the manufacturer’s instructions. Libraries were sequenced using the HiSeq2500 with paired end 76bp reads, followed by analysis with RTA v.1.12.4.2. The minimum depth of coverage was 150X and 80X for tumor and normal samples respectively.

### Whole transcriptome sequencing of bulk tumor samples

Whole transcriptome sequencing was performed at the Broad Institute genomic platform on 88 bulk tumor samples from 48 metastatic melanoma patients treated with CPB therapy (53% anti- CTLA4; 26% anti-PD-1; 11% anti-PD-L1 and 9% anti-CTLA-4+PD-1), using the Transcriptome Capture method (FFPE compatible) as previously described^11^. 250-500ng of purified total RNA with DV200 scores >30% was considered acceptable for library preparation and sequencing. First a stranded cDNA library from isolated RNA was constructed followed by hybridization of the library to a set of DNA oligonucleotide probes to enrich the library for mRNA transcript fragments (capturing 21,415 genes). The normalized, pooled libraries were loaded onto HiSeq2500 for a target of 50 million 2x76bp paired reads per sample.

### Whole-exome mutation calling

Using the Broad Picard pipeline, we aligned bams from all blood normal samples and tumor samples to hg19 using bwa 0.5.9^59^. For somatic mutation calling, we used the CGA WES Characterization pipeline within the Firecloud framework (https://app.terra.bio/#workspaces/broad-fc-getzlab-workflows/CGA_WES_Characterization_OpenAccess, ^60^). For mutation calling, we assessed the Agilent exome target regions for the Van Allen samples and the Illumina Capture Exome (ICE) target regions for samples from all other cohorts. We first assessed cross-sample contamination levels using ContEst^61^ and checked for sample swaps using Picard CrossCheckFingerprints (https://software.broadinstitute.org/gatk/documentation/tooldocs/4.0.1.0/picard_fingerprint_CrosscheckFingerprints.php). We then called somatic single nucleotide variants using MuTect^62^ with the cross-sample contamination estimates as lower bounds for allele fraction and called somatic indels using MuTect2 and Strelka (https://github.com/broadinstitute/gatk/tree/master/scripts/mutect2_wdl63). Next, we applied DeTiN to detect cases with evidence of tumor in the exome normal sample and recover somatic mutations that had been incorrectly filtered out due to alternate reads present in the normal sample^64^ and we annotated somatic mutations consequences using Oncotator^65^. We merged adjacent SNV calls and annotated them as di-nucleotide variants as CC>TT mutations are frequently detected in sun-exposed melanomas^38^. To filter the somatic mutation calls, we applied OxoG and FFPE Orientation Bias filters^66^ and we filtered out mutations present in a panel of TCGA normal samples or a panel of Illumina Capture Exome (ICE) normal samples^67^. We then used a BLAT Realignment filter^68^ to filter out mutations that were only supported by reads that had mapped ambiguously and finally filtered out mutations using a final panel of normal samples^67^ that included FFPE normal samples.

To exclude exome samples that were low quality, we excluded samples with tumor or normal mean bait coverage below 30 or cross-sample contamination values above 5%. WES samples from the Hugo cohort had high cross-sample contamination levels, so we did not include the WES DNA data from this cohort^3^. We retained one sample (Case4-Relapse) from the Zaretsky cohort with a contamination fraction of 5.4% as this represented an important clinical scenario (acquired resistance to PD-1 therapy after a response). Additionally, we excluded samples with tumor in normal estimates above 15%, as these samples would have reduced sensitivity for somatic mutation calling. Additionally, we excluded samples with ABSOLUTE estimated tumor purity below 10% (described in the ABSOLUTE section below^69^. After applying these filters, we retained a total of 258 WES samples from 189 patients as passing QC. Additionally, for analysis of outcomes, in order to use a single WES sample per patient for patients with multiple samples, if the patient had multiple WES samples we used the earliest biopsy available unless the patient was a non-responder for the first line of immunotherapy but a responder for a subsequent line of immunotherapy, in which case we used the later sample which was directly preceding or in some cases after the response. Additionally, for patient 33 in the Roh cohort, we used the post-treatment sample (33C) rather than the pre-treatment sample (33A) as the pre-treatment sample had tumor purity at our QC threshold level (10% tumor purity) which severely limited the analysis of clonality.

### Mutation Significance Analysis

We used MutSig2CV^70, 71^ to identify significantly mutated driver genes in the 189 melanoma WES samples. This meta-analysis cohort was smaller than previously published melanoma analyses which included patients not treated with immunotherapy and melanoma has a high mutation burden, so we were underpowered to discover novel melanoma drivers. We used a MutSig2CV q < 0.1 threshold to classify genes as significantly mutated (Supplementary Figure 3a). Additionally, we excluded significantly mutated genes with a median log_2_(TPM+1) below 1 in CCLE melanoma cell lines (analysis of CCLE melanoma RNA-Seq data described below).

### Mutation signature analysis

We used the SignatureAnalyzer Bayesian NMF approach to identify mutation signatures present in the somatic coding SNVs in the 189 melanoma WES samples^72–74^. As melanoma has a high mutation burden, we used the SignatureAnalyzer with the hypermutation option, and we excluded one sample (from patient 16 in the Roh cohort) which had evidence of an MSI signature (COSMIC Signature 26^75, 76^). We ran the Bayesian NMF with 50 random initializations, and converged to k=3 signatures, 44 converged to k=4 and 4 converged to k=5, so we chose the solution with k=4 with the maximum posterior probability (Supplementary Figure 4c). We compared the four signatures to COSMIC mutational signatures using cosine similarity (Supplementary Figure 4b) and we assigned mutations to signatures based on the association probability and computed the counts and fractions of mutations assigned to signatures by sample (Supplementary Figure 4a). As expected, we identified a UV signature and a CpG signature in many samples as well as a temozolomide (TMZ) signature (as a subset of patients had prior TMZ treatment in the Van Allen cohort)^17^. Additionally, we identified a fourth signature which primarily consisted of T to C mutations and had some similarity to COSMIC signature 26, but this signature was only present as a small proportion of mutations in a subset of samples (Supplementary Figure 4a).

### Neoantigen and TMB analysis

To identify neoantigens, we used normal WES samples to call germline MHC Class I alleles using POLYSOLVER^77^. We considered somatic single nucleotide and di-nucleotide variants as potential neoantigens and predicted the binding affinity of all possible 9mers and 10mer peptide sequences that overlapped the mutated residue using NetMHCPan 4.0^78–80^. We predicted binding affinities to all six germline MHC Class I alleles. We counted predicted binders as neoantigens if they had a NetMHCPan 4.0 percentile rank of 2 or lower (note that certain mutations may lead to multiple neoantigens). To compute tumor mutation burden (TMB), we counted the number of non-silent somatic SNVs, DNVs and Indels per sample, and we used log_10_(TMB) in continuous Cox models with overall survival. For the analysis of TMB above or below 10 mutations/Mb, we obtained the number of mutations per megabase by dividing the number of non-silent somatic SNVs, DNVs and Indels by the ICE target territory and multiplying by 10^6^, and we split samples using a threshold of 10 mutations/Mb.

### Association of SNVs with outcome

To investigate the relationship between somatic mutations and outcomes, we associated mutation status with survival using Cox models which included mutation status and log_10_(TMB) and we associated mutation status with response using logistic regression models which incorporated mutation status and log_10_(TMB). For the associations with survival and response, we tested loss of function mutation status (loss of function mutation or wild type) for all genes with 3 or more loss of function mutations and non-synonymous mutation status for all genes with 3 or more non-synonymous mutations. For the survival models, we used the *P* value of the mutation status in the combined Cox model with log_10_(TMB) and for the logistic regression response models, we used the *P* value of mutation status in the combined logistic regression model with log_10_(TMB).

### Somatic Copy Number Alteration (SCNA) Analysis

To assess somatic copy number alterations (SCNAs) in autosomes from WES data, we used GATK4 CNV (with GATK version 4.0.8.0, https://github.com/gatk-workflows/gatk4-somatic-cnvs^81^). We created a panel of normals to normalize the copy number data using GATK4 CNV CNV Somatic Panel Workflow using a set of 928 normal WES samples including TCGA melanoma, the matched normals from the MGH, Roh, Zaretsky and Hugo cohorts (the Hugo samples were included to improve the normalization of the Zaretsky samples sequenced by the same group, even though the Hugo samples failed cross-sample contamination QC) and a set of FFPE normal samples. We set the GATK CNV bin_length to 0 to skip binning (as recommended for WES data). Additionally, in order to be able to compare copy number results across cohorts, we used the hg19 Illumina Capture Exome (ICE) target set with 250bp of padding as the target list for all samples for collecting read counts for copy number calling. As this created a more heterogeneous distribution of coverage per target, we set --maximum-zeros-in-interval- percentage=1 (the threshold of the minimum percentage of samples in the panel of normals with zero-coverage in a target, and intervals failing this filter are removed).

We then used the GATK4 CNV Somatic Pair Workflow incorporating GATK ACNV to estimate somatic copy number alterations using the described panel of normals. To allow for heterozygous SNPs to be used in estimating allelic coverage, we set minimum-total-allele-count=10 (the minimum coverage required in the tumor to collect allelic counts at heterozygous SNP sites). To estimate allelic copy number in GATK4 CNV, we used the GATK set of frequently polymorphic SNP sites (gs://gatk-test-data/cnv/somatic/common_snps.interval_list). Additionally, we filtered potential germline or artifactual segments that appeared in both the germline blood normal and the tumor sample and segments at recurrent copy number segment breakpoints. After removing these events, gaps were imputed if the two neighboring segments had the same copy ratio and were 2.5Mb apart. Additionally, we filtered out allelic copy number segments supported by zero heterozygous SNPs.

### GISTIC analysis of significant SCNAs

We used GISTIC 2.0 to identify recurrently amplified and deleted regions in the 189 melanoma WES samples^82^. Similar to the mutation significance analysis, we were underpowered to discover novel melanoma drivers in comparison to previous studies. We ran GISTIC 2.0 on the GATK4 CNV segment mean log_2_ copy ratio (after the post-processing to remove potential germline and artifactual events as described).

### Analysis of tumor purity and ploidy using ABSOLUTE

In order to estimate tumor purity and ploidy as well as integer copy number, we used ABSOLUTE^69, 83^. Using the called somatic mutations and somatic copy number alterations as input, we then use ABSOLUTE to generate candidate purity/ploidy solutions. We then manually reviewed these potential ABSOLUTE solutions and selected the solution most concordant with the copy number and mutation data. Finally, after obtaining the integer somatic copy number segments, we imputed gaps by extending the neighboring segments to meet in the middle of the gap and we annotated the integer copy number and LOH status of GENCODE v19 genes. For analyses of purity and outcomes in the primary cohort, we excluded two samples from the Zaretsky cohort (Case2-Baseline and Case3-Baseline) as they were derived from cell lines.

### Analysis of longitudinal samples using PhylogicNDT

To investigate clonal dynamics in longitudinal WES samples, we used ABSOLUTE and PhylogicNDT (https://github.com/broadinstitute/PhylogicNDT)^84^. After calling somatic mutations in all 123 samples from the 54 patients with WES data at multiple time points, we then took the union of mutations called in each patient and then calculated the number of reference and alternate reads present at the union of the mutated sites in each sample from the patient. For this analysis of the union of mutated sites, we used non-duplicate reads with mapping quality of at least 5 and with a recalibrated base quality scores of at least 20 at the mutated site. We then called ABSOLUTE solutions for all samples and clustered the mutation clonality across time points using the multidimensional Dirichlet process model to estimate the mutation cancer cell fractions (CCFs) and somatic copy number alteration clonality across samples. We plotted clonality of mutations and gene level copy number in paired biopsies, with 108 biopsies in total from 54 patients (Supplementary Table 1, Supplementary Figure 7). For patients in the MGH cohort with three or more biopsies, we used the earliest two WES samples available, except for patients 208T and 272T as these patients both had responses but then developed resistance at the time of the third sample. For patient 208T, we used samples 208A and 208C and for patient 272T we used samples 272A and 272C^11^. In addition to the analysis of longitudinal samples, we also separately applied the PhylogicNDT mutation clustering to single samples in order to estimate the mutation CCFs, and we counted mutations as clonal if they had estimated CCF≥0.85.

### RNA-Seq gene expression analysis

In order to quantify gene expression in RNA-Seq data, we used the GTEx RNA-Seq pipeline (https://github.com/broadinstitute/gtex-pipeline/)^85^, which uses STAR^86^ 2 pass alignment followed by quantification of TPMs using RSEM^87^. We quantified gene expression in TPM for all transcripts using the GENCODE v19 reference transcriptome. We derived quality control metrics from the aligned bams using RNA-SeqQC (https://github.com/getzlab/rnaseqc), and we excluded samples with below 15000 genes detected (where genes with 5 or more unambiguous reads were considered detected) or Exon CV MAD > 1. The Exon CV MAD is the median absolute deviation of the coefficient of variation of the exonic coverage (which excludes the first and last 500bp of a gene). Additionally, we performed PCA on the log_2_(TPM + 1) values for all protein coding genes using the MGH pre-treatment samples (including only pre-treatment A RNA-Seq samples and not post-treatment B or C samples), and we excluded two additional samples that were PCA outliers (samples 346AR and 9AR). We also excluded the one post-treatment sample from the Hugo cohort from analysis (Pt16-OnTx). Additionally, we re-processed the TCGA melanoma (SKCM) RNA-Seq data using the GTEx pipeline to obtain log_2_(TPM+1) values.

When we performed PCA on the log_2_(TPM+1) values of protein coding genes (excluding genes with zero expression in all samples) for the 154 pre-treatment samples from the primary cohort, we noted significant batch effects between cohorts (Supplementary Figure 16a). The cohorts clustered in batches based on the RNA-Seq method (polyA selection for Hugo and Riaz vs. transcriptome capture for MGH and Van Allen), so we applied ComBat^88^ in order to remove this batch effect. We also set all negative values following ComBat batch correction to zero, as the input log_2_(TPM+1) values were non-negative. After this batch effects correction, the separate cohorts were overlapping in a PCA (Supplementary Figure 16a).

For the secondary cohort, we obtained log_2_(TPM+1) values in the same way using the GTEx pipeline. Upon inspecting the RNASeqQC metrics for the Liu cohort, we found that 14 of the initial 121 samples failed the 15000 genes detected threshold or the Exon CV MAD > 1 threshold, so we excluded these samples, leaving 107 samples. For the Gide cohort, no samples failed the QC thresholds. For the secondary cohort analysis of RNA-Seq samples, we combined the 107 Liu samples with the 73 Gide pre-treatment samples (excluding the 16 Gide early during treatment samples), leaving 180 samples in total. We observed similar batch effects in the PCA of log_2_(TPM+1) values for the Gide and Liu cohorts (Supplementary Figure 23a), so we applied ComBat to the full secondary cohort which reduced the batch-specific clustering in PCA.

### Quantification of TCR and Ig levels in WES and RNA-Seq

To quantify T and B cell infiltration, we used MixCR v3.0.3^89, 90^. We used the MixCR analyze shotgun pipeline with DNA or RNA as the starting material. We excluded TCR or Ig clonotypes with rearrangements that had a stop codon or resulted in an out-of-frame sequence. We quantified the number of TCR or Ig reads by summing the clone counts of reads assigned to clonotypes. For TCRs, we included TCR alpha, beta, delta and gamma sequences. By counting these reads, we potentially included rearrangements from CD8 T cells, CD4 T cells (some of which may be regulatory T cells) and γδ T cells. For the analysis of RNA-Seq data, in rare cases clonotypes were aligned to both a TCR and an Ig (with MixCR V region alignment allVHitsWithScore containing both a TCR and an Ig), and we included these reads in both the TCR and the Ig counts.

As different samples had different depths of sequencing, we normalized the TCR and Ig read counts by sample coverage. For RNA-Seq, we used the number of mapped reads from RNASeqQC, and for WES we used the number of reads aligned from samtools idxstats. To create the T cell burden (TCB) and B cell burden (BCB) metrics, we computed TCB=(1+TCR read count)/(aligned reads/10^6) and BCB=(1+Ig read count)/(aligned reads/10^6)], with appropriate TCR/Ig read counts and aligned read counts separately for DNA and RNA. Additionally, we plotted TCB and BCB on log_10_ scales and we used log_10_(TCB) and log_10_(BCB) for outcome analysis using Cox or logistic regression models.

TCB_RNA_ values were slightly higher in samples from the Hugo cohort (Supplementary Figure 8a) than in samples from other cohorts, but we saw significant batch effects for both TCB_DNA_ and BCB_DNA_ (Supplementary Figure 9a). As a result, we decided to dichotomize TCB_DNA_ and BCB_DNA_ within cohorts, so we calculated median(TCB_DNA_) separately for each cohort and we labeled samples as TCB ^high^ if they had TCB_DNA_ > median(TCB_DNA_) using the cohort-specific median (with similar analysis for BCB_DNA_). For analyses of TCB_DNA_/BCB_DNA_ and outcomes in the primary cohort, we again excluded the two cell line samples from the Zaretsky cohort (Case2- Baseline and Case3-Baseline).

When we compared TCB_DNA_ and TCB_RNA_, there were a subset of samples in the MGH cohort for which DNA and RNA was extracted from different locations of a tumor biopsy. These samples could have different levels of TCB in RNA and DNA due to sampling rather than technical factors, so when we looked at the correlation between TCB_DNA_ and TCB_RNA_ (as well as the correlation between BCB_DNA_ and BCB_DNA_), we only included Van Allen samples and MGH samples with DNA and RNA extracted from the same location (n=35 total). For analysis of TCB_DNA_ and BCB_DNA_ from longitudinal WES samples, we analyzed samples from the same 54 patients with longitudinal WES samples, and for patients in the MGH cohort with three or more biopsies, we used the earliest two WES samples available from each patient. Additionally, for analysis of TCB_RNA_ and BCB_RNA_ from longitudinal RNA-Seq, we analyzed 172 paired biopsies from 86 patients across the Riaz, Gide and MGH cohorts, and again for patients in the MGH cohort with three or more biopsies, we used the earliest two RNA-Seq samples available from each patient.

### TCR/Ig overlap analysis

In order to further establish that TCRs and Igs were shared between WES and RNA-Seq data, we assessed the degree of overlap between TCR and Ig sequences in primary cohort samples with DNA and RNA extracted from the same location (n=35 total). Using each DNA CDR3 region from the MixCR output, we performed pairwise global alignments using the Needleman-Wunsch algorithm (implemented in the R biostrings package in function pairwiseAlignment with gapOpening=10 and gapExtension=4) between the DNA CDR3 region and all RNA CDR3 regions in the MixCR output from the paired RNA sample. If the top alignment had a Needleman-Wunsch alignment score greater than 25 and the top alignment had S_i_=number of mismatches+total insertion length+total deletion length≤2, then we counted the TCR or lg pair as an overlap. We then calculated the number patients with DNA/RNA overlaps for TCRs and for lgs.

### RNA-Seq Tumor Subtyping

In order to identify melanoma subtypes using bulk RNA-Seq data, we applied a Bayesian NMF based clustering method to 469 TGCA melanoma RNA-Seq samples^38, 44, 45^. We preprocessed the data by removing non-protein-coding genes and by retaining only the 25% of genes with the highest standard deviation of expression across samples, and we removed all genes that were expressed at 0 TPM in at least 10% of samples. After these steps, 2684 genes remained. Next, we transformed the matrix of TPMs into fold changes by subtracting the median of each gene from the log_2_(TPM+1) values for that gene. To cluster samples, we created a distance matrix by calculating the Spearman correlation between each pair of samples and then performed repeated hierarchical clustering with K (the number of clusters) between 2 and 10. We performed hierarchical clustering using 80% sampling and average linkage, and we repeated this clustering K*500 times for each K. We created a consensus matrix M_K_ by calculating the number of times that each pair of samples clustered together in the repeated hierarchical clusterings for a given K. Then, we summed all M_K_ and normalized the final matrix M* by the total number of iterations. We then used Bayesian non-negative matrix factorization with a half-normal prior on the M* matrix to determine the optimal number of clusters K*. In this setup where we are approximating M* ∼ H^T^H, the H matrix represents the association of samples to clusters. The most frequent solution was K*=5, so we selected this clustering solution. We also obtained the normalized matrix H* by normalizing each column so that the values of each sample’s h_ij_ sample- to-cluster association sum to 1.

To identify marker genes for clusters, we took the full TCGA SKCM log_2_(TPM+1) matrix X with 19820 protein-coding genes and performed non-negative matrix factorization by approximating X ∼ WH*. In this case, the values in W represent the association of a gene to each cluster. We obtained the normalized matrix W^*^ by again normalizing each column to sum to 1.

We used a previously developed approach in order to select marker genes for each subtype and project the subtypes identified in TCGA samples to new data using the expression of the reduced set of marker genes^44^. In order to optimize the subtype classifier, we varied parameters for selecting cluster marker genes for the subtype classification. We did not consider genes with 0 TPM in 10% or more samples as candidate marker genes. When selecting marker genes, we used only marker genes which were overexpressed in a cluster relative to the other clusters. We selected marker genes which had normalized association to a cluster greater than a threshold parameter W_cut_ and we limited the number of marker genes for each cluster based on the threshold parameter gene_cut_. For the marker genes in each cluster, we considered only genes with a mean difference in log_2_(fold change) across clusters of 0.5 or better. In order to identify the optimal classifier, we performed a parameter sweep with W_cut_ from 0.1 to 0.96 in increments of 0.01 and gene_cut_ from 30 to 250 in increments of 10. Using these parameters, we selected the reduced set of marker genes g, and then we performed the matrix factorization X_g_∼ W*_g_H, and we assigned samples to clusters using this H matrix. We measured the performance of the classifier using the adjusted rand index when comparing the projected cluster labels to the original cluster labels. We identified W_cut_=0.71 and gene_cut_=65 as the optimal parameter setting, which resulted in an adjusted rand index of 0.653 for the TCGA SKCM samples.

We then used the marker genes and the subtype membership for TCGA samples to compare our melanoma subtypes to previous melanoma subtype classification schemes. The TCGA melanoma RNA-Seq subtyping identified 3 subtypes: an immune subtype, an MITF high subtype and a keratin subtype. The TCGA MITF low and Immune subtypes had strong overlap with two of our subtypes (Supplementary Figure 14b), and the marker genes were consistent with many T cell genes identified as markers for one (*CD2* and *CD8A*) and neural crest marker genes (such as *AXL* and *NGFR*) highly expressed in the other (Supplementary Figure 15a-b). Based on these results, we denoted these subtypes Immune and MITF low. Another subtype had many keratin marker genes such as (*KRT10*, *KRT5* and *KRT19*), suggesting a high degree of keratinocyte infiltration, so we denoted this subtype Keratin high. Next, when we compared our subtype assignments to a melanoma differentiation-based subtype classification scheme, we saw a high degree of overlap between some subtypes (Supplementary Figure 14d).^46^ The analysis in the Tsoi paper attempted to remove non-tumor intrinsic signals including immune infiltration and keratin expression, whereas our goal was to incorporate both tumor-intrinsic markers and potential immune markers. The Tsoi melanocytic subtype strongly overlapped with one of our subtypes, and this subtype had the highest expression of melanocyte markers such as *PMEL*, *MITF* and *MLANA*, so we denoted this subtype MITF high. The MITF high and MITF low subtypes match the previously recognized melanoma differentiation axis with well differentiated melanocyte-like tumors expressing *MITF* and poorly differentiated neural-crest-like tumors expressing *AXL*^46–48^. The final subtype had the highest degree of overlap with the Tsoi transitory subtype (Supplementary Figure 14d) which had an intermediate differentiation state, so we denoted this the Intermediate subtype (though the *MITF* expression level in this subtype is similar to that of the MITF high subtype). Additionally, we looked at the relationships between TMB (by reprocessing TCGA melanoma WES data using the same somatic mutation calling pipeline), TCB_RNA_, BCB_RNA_ and tumor purity (from the previous TCGA melanoma analysis)^38^.

Finally, we sought to classify the tumors in the melanoma immunotherapy meta-analysis cohort by their subtype. We used the ComBat batch corrected log_2_(TPM+1) values for the 154 primary cohort RNA-Seq samples and preprocessed the data by removing genes with zero TPM values in 10% or more samples and transforming the TPM values to log_2_(fold change) values using median centering. This preprocessing removed a subset of the marker genes which were selected in the subtype classifier. Then, we used the weights inferred for the matrix W*_g_ and the log_2_(fold change) values for the selected genes in the immunotherapy expression data to perform the approximation X_g CPB_∼ W*_g_H_CPB_ from which we could identify the subtypes of the immunotherapy samples using the normalized matrix H*_CPB_. We separately applied this same procedure to the ComBat batch corrected log_2_(TPM+1) values for the 180 secondary cohort samples in order to determine their subtype memberships.

### Differential Expression Analysis

We used DESeq2^91^ in order to identify differentially expressed genes from the RNA-Seq data. Based on the previous batch effects that we identified between cohorts, we used the cohort as an additional covariate in all DESeq2-based differential expression analyses. We compared patients with long OS (overall survival > 1 year) vs. short OS (overall survival < 1 year) and responders vs. non-responders in 154 samples in the primary cohort (153 for overall survival as one patient did not have OS data), while including batch as a covariate. We analyzed genes with median log_2_(TPM+1)>1 and we performed Benjamini-Hochberg multiple hypothesis correction using the DESeq2 *P* values. We considered genes with q<0.05 as differentially expressed.

### Analysis of CCLE and Human Protein Atlas (HPA) RNA-Seq data

To assess whether the differentially expressed genes were expressed in melanoma cells, immune cells or both cell types, we assessed gene expression in CCLE cell lines and bulk RNA-Seq data of HPA blood cell types^55, 92^. We downloaded CCLE RNA-Seq TPM data from https://data.broadinstitute.org/ccle/CCLE_RNAseq_rsem_genes_tpm_20180929.txt.gz and used data from 49 melanoma cell lines. When we performed PCA on the log_2_(TPM+1) values, we noted 2 outliers in PCA (CJM_SKIN and LOXIMVI_SKIN), and we removed these two cell lines from the data set, leaving 47 melanoma cell lines. Then, we computed the median log_2_(TPM+1) values for the differentially expressed genes of interest. To quantify expression of genes in immune cells, we downloaded the HPA blood expression data from https://www.proteinatlas.org/download/rna_blood_cell_sample_tpm_m.tsv.zip and took the median log_2_(TPM+1). We used thresholds of log_2_(TPM+1) of 1 for both cohorts to determine whether genes were expressed in immune cells, expressed in tumor cells, expressed in both or had low expression in both.

Additionally, we wanted to determine which immune cell types and which melanoma subtypes had high expression of these differentially expressed genes of interest. First, we assessed which of the differentially expressed genes were coexpressed by calculating the pairwise spearman correlations of all genes in the 154 primary cohort samples (Supplementary Figure 18a, 19a). Then, for the differentially expressed genes that were higher expressed in responders or patients with long OS (which were usually higher expressed in immune cells than in melanoma cell lines), we transformed the HPA log_2_(TPM+1) values to z-scores and took the mean of each gene’s z-scores for each cell type (Supplementary Figure 18b, 19b). Additionally, for each of these genes, we ranked the 19 HPA cell types by log_2_(TPM+1) of the gene, with 1 being the highest expression and 19 being the lowest expression, allowing for ties. We then ordered each cell type by their median rank across the genes of interest (Supplementary Figure 18c, 19c). Additionally, using the primary cohort batch corrected log_2_(TPM+1) data and melanoma subtypes, we repeated the same analysis of mean z-scored expression by subtype and subtype ranking (Supplementary Figure 18d-e, 19d-e). These analyses suggested that lymphocytes had the highest expression of the genes overexpressed in responders and patients with long OS, and that the genes overexpressed in non-responder and patients with short OS were highest expressed in the MITF low melanoma subtype.

### Construction of metagene models

To create gene signatures using the differentially expressed genes, we constructed metagenes from the differentially expressed genes in the long OS vs. short OS and the responder vs. non-responder analyses. For each analysis, we split genes based on their effect size, consisting of 75 genes overexpressed in responders, 55 genes overexpressed in patients with long OS, 26 genes overexpressed in non-responders and 28 genes overexpressed in patients with short OS. We created a metagene using each of these four gene lists. In order to score individual patients, we transformed log_2_(TPM+1) values to z-scores for each gene across patients (so that genes with low expression would not be penalized) and calculated the mean of the gene z-scores for each individual patient.

### Predicting survival and response using gene pair models

To create simple models to predict outcome using gene expression that could potentially incorporate both tumor and immune components, we considered the genes we identified as differentially expressed and tested all genes pairs as predictors of response and overall survival. For this analysis, we used the batch-effects corrected log_2_(TPM+1) values, whereas in the DESeq2 differential expression analysis we used the raw count data and included batch as a DESeq2 model covariate. For analysis of response, we used all 154 patients, but for analysis of survival, we analyzed 153 samples because one patient from the Hugo cohort did not have overall survival data. We performed the gene pair model analysis using 1) all 3403 unique pairs of the 83 genes differentially expressed between patients with long OS and short OS 2) all 5050 unique pairs of the 101 genes differentially expressed between responders and non-responders. For each analysis, we also considered the corresponding pair of metagenes as an additional model (for 1) the long OS and short OS metagenes and for 2) the R and NR metagenes). For each gene (or metagene) pair model, we tested the association with survival using Cox proportional hazards models incorporating two genes, and we calculated the significance with the log-rank *P* value and the performance with the C-index. Similarly, we tested the association between gene pairs and response using a logistic regression model incorporating two genes, and we calculated the significance with a P value testing whether the null hypothesis of an AUC=0.5 can be rejected (implemented in the R verification package^93^) and the model performance with the model AUC (using the logistic regression ŷ values to rank samples).

Next, we performed multiple hypothesis corrections on the model significance values separately for logistic regression model *P* values and for Cox model *P* values. Since the tested gene pairs were genes associated with outcomes in the DESeq2 analysis, many of the gene pair models had low *P* values, and the distribution of gene pair model *P* values strongly deviated from a uniform distribution. Additionally, we wanted to very stringently select significant models, so we used a Bonferroni multiple hypothesis correction rather than a Benjamini-Hochberg correction. We performed Bonferroni correction on the *P* values for the response logistic regression models and the survival Cox models.

To identify gene pair models which predicted both response and survival, we selected gene models with Bonferroni corrected P<0.05 for both response and survival predictions. For the analysis using the 5050 gene pairs derived from the responder vs. non-responder differential expression analysis and the response metagene pair model, 39 gene pair models had Bonferroni corrected P<0.05 for prediction of response (one of which was the response metagene pair model), but 0 gene pair models had Bonferroni corrected P<0.05 for predictions of survival (Supplementary Figure 20f). Thus, the gene pairs derived from the responder vs. non-responder differential expression analysis were predictors of response but not survival. Additionally, when we looked at the performance of the gene pair models by the original differential expression effect size (R genes had higher expression in responders, and NR genes had higher expression in non-responders), gene pairs models with a R gene and an NR gene had higher C-index values than other model types but gene pair models with two NR genes had higher AUC values than other model types (Supplementary Figure 20h-i). For the analysis using the 3403 gene pairs derived from the long OS vs. short OS differential expression analysis and the long/short OS metagene pair model, 67 gene pair models had Bonferroni corrected P<0.05 for prediction of survival and 10 models had Bonferroni corrected P<0.05 for predictions of response. There were 3 gene pair models that had Bonferroni corrected P<0.05 for both predictions of response and survival: *MAP4K1*&*TBX3*, *MAP4K1*&*AGER* and the long OS/short OS metagene pair model (Figure 4a). These results suggest that some gene pairs derived from the long OS vs. short OS differential expression analysis were able to predict both survival and response. Furthermore, gene pairs models with a long OS gene (higher expressed in patients with long OS) and a short OS gene (higher expressed in patients with short OS) had higher C-index values than other model types, but the gene pairs models with two short OS genes had very slightly higher AUC values than gene pair models with a long OS gene and a short OS gene, though the mean AUC values were very similar (mean AUC for models with a long OS gene and a short OS gene was 0.612 and mean AUC for models with two short OS genes was 0.619, Wilcox *P*=0.026, Supplementary Figure 20d- e). Additionally, the 3 gene pairs which passed Bonferroni-corrected *P* value thresholds for survival and response predictions were all pairs which incorporated a long OS gene and a short OS gene. These results suggest that gene model pairs which incorporate a long OS gene or an R gene (often immune-expressed genes) as well as a short OS gene or an NR gene (often tumor-expressed genes) have improved predictions of survival than other gene pair model types, but gene pair models with two poor outcome-associated genes can sometimes perform best in response prediction. Based on the performance of these models, we decided to compare the performance of these three gene pair models to that of other immunotherapy models and assess their performance in an independent data set.

### Gene pair model cross-validation analysis

To assess the robustness of the gene pair model discovery, we performed a cross-validation analysis using the primary cohort data. We used different splits for training and validation sets of 80%/20%, 75%/25%, 70%/30%, 66%/33% and 50%/50%. Using training sets composed of subsets of the cohort, we repeated the differential expression analysis of responders vs. non-responders and long OS vs. short OS (DESeq2 q<0.05). We tested the association between all differentially expressed gene pairs (within gene pair type, response or OS) and response and survival using logistic regression and Cox models as described. In addition to the gene pair models, we also constructed responder/non-responder and long OS/short OS metagene pair models using the differentially expressed genes discovered in each training set cross-validation differential expression analysis. Due to the reduced power in the training sets, there were sometimes few differentially expressed genes detected in the training sets, so we also constructed metagene pair models by ranking differentially expressed gene by *P* values and creating a metagene using the top 25 genes overexpressed in responders or patients with long OS and a separate metagene using the top 25 genes overexpressed in non-responders or patients with short OS (even if these genes had DESeq2 q>0.05). Then, for each training set we performed the discovery analysis of gene pair models using each set of differentially expressed genes by identifying gene pairs with Bonferroni-corrected log-rank *P*<0.05 and/or Bonferroni-corrected response AUC P<0.05. Finally, we attempted to validate the discovered models in a validation set composed of all of the remaining samples in the cohort, checking for validation set Bonferroni-corrected log-rank *P*<0.05 and Bonferroni-corrected response AUC P<0.05.

Based on the results from this cross validation, we found that increasing the training set size increased the number of models discovered with Bonferroni-corrected *P*<0.05 (Supplementary Figure 22a-b). Even with 80% of the dataset as a training set, discovery was still increasing, which suggests that the full cohort analysis is likely underpowered. For the metagene models, increasing the training set size increased the robustness of the long OS metagene signatures, whereas the short OS metagene signatures were more variable across patients (Supplementary Figure 22c-f). Finally, while the top gene pair models were frequently discovered training data subsets (Supplementary Figure 22g-h), they were very rarely statistically significant in validation sets (Supplementary Figure 22i-j), supporting the need for larger clinically annotated immunotherapy datasets.

### Gene pair model validation analysis

We attempted to validate the three gene pair models (*MAP4K1*&*TBX3*, *MAP4K1*&*AGER* and the long OS/short OS metagene pair model) developed using the primary cohort samples in the independent secondary cohort. To be validated, we required these models to achieve significance below the Bonferroni corrected threshold of *P*=0.05/3=0.0167 with both the OS log-rank *P* value<0.0167 and the response AUC *P* value<0.0167 in the secondary cohort. Each of the three gene pair models passed both of these thresholds in the secondary cohort (Figure 4g), so we considered them to be validated in an independent cohort.

In order to visualize the survival of high and low risk groups within the primary and secondary cohorts for the *MAP4K1*&*TBX3* model, we divided patients into groups depending on whether their ŷ values in overall survival Cox models with *MAP4K1*&*TBX3* were above or below median (Figure 4k).

### Performance of external gene-expression models

In order to compare the performance of the top gene pair models to that of other predictors of immunotherapy outcomes, we evaluated a set of external models. We also included TCB_RNA_ as an additional predictor. Additionally, we included *CD274* expression as a simple model. We tested the CYT model^53^ by taking the geometric mean of *PRF1* and *GZMB* expression. We tested the GEP model^22^ by taking the mean expression of the GEP signature genes. We tested the IMPRES model^51^ by counting the number of IMPRES gene pairs that had the expected relationship (each IMPRES high expression gene having greater expression than the paired IMPRES low expression gene). We tested the MHC II model^4^ by performing ssGSEA using the MHC II gene list, and we z-scored the ssGESEA score so that hazard ratios would be more interpretable. Finally, we tested the TIDE model^52^ by calculating TIDE scores using the TIDE web portal (http://tide.dfci.harvard.edu/). The TIDE portal includes separate models for melanoma samples from patients that are immunotherapy naive or had prior immunotherapy. Additionally, mean centering is recommended as a preprocessing step. Thus, we separately mean centered the batch-effects corrected log_2_(TPM+1) data for patients with prior CTLA4 treatment and patients with no prior CTLA4 treatment in the primary cohort, uploaded these matrices to the TIDE web portal and obtained TIDE scores. Separately, for the secondary cohort, we repeated the same process, and for TIDE we separately processed the patients with prior CTLA4 in the Liu cohort (using the melanoma model for patients with prior immunotherapy) and then the combination of the CTLA4 naive patients in the Liu cohort with the Gide cohort (as there was no information regard prior immunotherapy treatment for the Gide cohort). For testing the associations between these models and outcome, we used the predictor values as described, but for visualizing the predictor values across patients, we z-scored the predictor values (Figure 4b, Supplementary Figure 24a). After validating the three gene pair models, we compared the response classification performance of all pairs of models in the secondary cohort using DeLong’s test. Finally, we tested models including the top gene pairs and treatment (PD1 vs. combined CTLA4+PD1) in the secondary cohort (Supplementary Figure 24j-k) and we evaluated the performance of all gene pair models and external models within each cohort separately while including treatment (PD1 vs. CTLA4+PD1) as a covariate in the Liu cohort model (Supplementary Figure 25a-j).

### *TBX3* melanoma cell line GSEA

To identify pathways associated with *TBX3* expression in melanoma, we again analyzed RNA-Seq from the 47 CCLE melanoma cell lines and ranked genes by their spearman correlation with *TBX3*. Then, we performed ranked gene set enrichment analysis using the fgsea R package^94^ with all GO terms (Supplementary Figure 26a). Negatively correlated genes were enriched for pigmentation pathways, which is consistent with *TBX3* being expressed in poorly differentiated neural crest-like melanomas but not well differentiated MITF high melanomas.

### Analysis of Jerby-Arnon melanoma scRNA data

To assess whether genes of interest were expressed in melanoma cells or melanoma tumor-infiltrating immune cells, we downloaded scRNA data from the Jerby-Arnon cohort^56^ from https://singlecell.broadinstitute.org/single_cell/study/SCP109/melanoma-immunotherapy-resistance. We plotted the expression of genes of interest using the TPM values and the pre-computed normal and malignant cell tSNE coordinates. To test whether *TBX3* was higher expressed in NGFR-expressing melanoma cells (which would be consistent with *TBX3* being higher expressed in poorly differentiated neural-crest-like melanoma cells), we performed Wilcoxon tests of *TBX3* expression in melanoma cells vs. immune cells and *TBX3* expression in melanoma cells with an NGFR TPM of 0 vs. *TBX3* expression in melanoma cells with an NGFR TPM>0.

### Analysis of Tsoi cell line data

To assess whether *TBX3* was expressed in poorly differentiated melanoma cell lines^46^, we used the Tsoi web portal (https://systems.crump.ucla.edu/dediff/index.php) and checked the expression of *MITF* and *AXL*, which are known to be expressed in well and poorly differentiated melanoma cells respectively, as well as *TBX3*. *TBX3* was expressed in poorly and intermediately differentiated melanomas but not in well differentiated melanomas (Figure 4m).

### Data availability

Raw sequencing data (WES and RNAseq) from this study (e.g. MGH cohort) will be deposited in dbGAP database upon publication. For the primary cohort, publicly available data was used. The Van Allen^17^ WES and bulk RNA dataset used in this study is available in dbGAP database under accession number phs000452.v2.p1. The Roh^36^ WES dataset is available under BioProject accession number PRJNA369259. The Zaretsky^12^ WES dataset is available in the National Center for Biotechnology Information Sequence Read Archive under accession number SRP076315. The Riaz^6^ bulk RNA dataset used in this study is available under BioProject accession number PRJNA356761 or SRA SRP094781. The Hugo^37^ bulk RNA dataset used in this study is available in GEO (Gene Expression Omnibus) under accession number GSE78220. For the secondary cohort, publicly available data from two recent publications was used. The Gide^54^ bulk RNA dataset is available in the European Nucleotide Archive (ENA) under accession number PRJEB23709. The Liu^4^ bulk RNA dataset is available in dbGAP under accession number phs000452.v3.p1.

## Acknowledgements

First and foremost, we would like to express our deep gratitude to the patients and their families for making this study possible. We thank the Genomics Platform of the Broad Institute of Harvard and MIT for the whole exome and bulk RNA sequencing performed in this study. This study was supported by grants from Broad Next-10 (N.H), Cancer Research Institute (Clinic and Laboratory Integration Program, N.H), Adelson Medical Research Foundation (N.H), the David P. Ryan, MD, Chair funded by a gift from Arthur, Sandra and Sarah Irving (N.H.), the Paul C. Zamecnick, MD, Chair in Oncology at MGH (G.G), NIH/NHGRI 5U54HG003067 (S.G), and NIH/NCI R01CA208756 (N.H). We also thank the following grants for their support: Tosteson & Fund for Medical Discovery fellowships (M.S.F) and NIH 1T32CA207021-01 (J.H.C)

## Author contributions

Conceptualization, S.S.F., M.S.F., M.M., G.G., and N.H.

Methodology, S.S.F., M.S.F., M.M., G.G., and N.H.

Software, S.S.F., J.K., C.S., A.R., M.B.A., K.Y., I.L., L.E., O.G.S., D.L., D.R., F.A., J.C.Z., G.H., and Z.L.

Validation, S.S.F., M.S.F., A.L.K.G., I.G., T.J.L., and E.M.B.

Formal Analysis, S.S.F., M.S.F., J.K., C.S., A.R., M.B.A., K.Y., I.L., L.E., O.G.S., D.L., D.R., F.A., J.C.Z., G.H., and Z.L.

Investigation, S.S.F., M.S.F., A.L.K.G., I.G., T.J.L., and E.M.B.

Resources, S.S.F., M.S.F., A.L.K.G., I.G., T.J.L., E.M.B., D.T.F., T.S., J.H.C., M.B.R., M.R.H.,

H.C.V., S.M.B., Y.J.J., S.G., D.P.L., L.M.D., A.O.S.R., J.A.W., K.T.F., R.J.S., and G.M.B.

Data Curation, S.S.F., M.S.F., and J.K.

Writing- Original Draft. S.S.F., M.S.F., M.M., G.G., and N.H.

Writing- Review & Editing, S.S.F., M.S.F., G.M.B., M.M., G.G., and N.H. Visualization, S.S.F., and M.S.F.

Supervision, M.M., G.G., and N.H. Funding Acquisition, S.G., G.G. and N.H.

## Supplementary Material

### Supplementary Figures

#### List of supplementary figures

1. Supplementary Figure 1: Primary cohort composition in meta-analysis

2. Supplementary Figure 2: Analysis workflow

3. Supplementary Figure 3: Primary cohort WES analysis

4. Supplementary Figure 4: Mutational signatures

5. Supplementary Figure 5: TMB, neoantigens and tumor purity

6. Supplementary Figure 6: Simple mutation models

7. Supplementary Figure 7: Comparison between paired pre-treatment and post-treatment biopsies

8. Supplementary Figure 8: Performance of TCBRNA and BCBRNA models

9. Supplementary Figure 9: Performance of TCBDNA and BCBDNA models

10. Supplementary Figure 10: Performance of integrative DNA-based models for survival and response prediction

11. Supplementary Figure 11: Performance of integrative DNA-based models for survival and response prediction for TCGA melanoma stage III-IV cases

12. Supplementary Figure 12: Dynamics of DNA and RNA-based TCB and BCB abundance between paired pre-treatment and post-treatment biopsies

13. Supplementary Figure 13: Performance of DNA and RNA-based TCB and BCB dynamics

14. Supplementary Figure 14: Subtypes identified using NMF clustering of TCGA melanoma RNA-seq

15. Supplementary Figure 15: TCGA RNA-seq subtypes and their tumor related features

16. Supplementary Figure 16: Subtype classification for pre-immunotherapy RNA-seq samples

17. Supplementary Figure 17: Genes associated with response in the primary cohort

18. Supplementary Figure 18: Expression patterns for long and short OS differentially expressed genes in the primary cohort

19. Supplementary Figure 19: Expression patterns for responder and non-responder differentially expressed genes in the primary cohort

20. Supplementary Figure 20: Performance of gene-pair models in the primary cohort

21. Supplementary Figure 21: Expression of genes from top gene pair models and gene pair model performance

22. Supplementary Figure 22: Cross-validation of gene pair model discovery and validation in the primary cohort

23. Supplementary Figure 23: Batch effects correction and melanoma subtyping for the secondary cohort

24. Supplementary Figure 24: Performance of top gene pair models in the secondary cohort

25. Supplementary Figure 25: Performance of all models within each cohort separately

26. Supplementary Figure 26: Analysis of melanoma *TBX3* expression

### Supplementary Tables

#### List of supplementary tables

1. Supplementary Table 1: Clinical data for DNA (WES) cohorts

2. Supplementary Table 2: Clinical data for RNA (bulk RNA sequencing) cohorts

3. Supplementary Table 3: WES Mutation table

4. Supplementary Table 4: WES pre/post mutation table with clonality

5. Supplementary Table 5: WES copy number table for pre (n=189) samples

6. Supplementary Table 6: WES pre/post DNA copy number table

7. Supplementary Table 7: WES sample metrics for all DNA samples

8. Supplementary Table 8: WES DNA counts - TMB, indel counts, neoantigens, purity, clonal TMB, clonal neoantigens, TCB, BCB

9. Supplementary Table 9: WES Mutsig2CV table

10. Supplementary Table 10: WES GISTIC table

11. Supplementary Table 11: WES mutation signatures table

12. Supplementary Table 12: Single sample DNA mutation association analysis

13. Supplementary Table 13: TCR/Ig counts and TCB/BCB for RNA samples

14. Supplementary Table 14: TCR/Ig counts and TCB/BCB for DNA samples

15. Supplementary Table 15: TCB and BCB for matched DNA/RNA samples

16. Supplementary Table 16: TCR overlap count table

17. Supplementary Table 17: RNA counts - TCB, BCB, GEP

18. Supplementary Table 18: TCGA DNA/RNA - TMB, TCB, BCB, GEP, stage, external purity, TCGA clinical variables (stage, survival)

19. Supplementary Table 19: Pre/post TCB/BCB counts and outcomes

20. Supplementary Table 20: log_2_(TPM+1) values for TCGA samples

21. Supplementary Table 21: NMF table for TCGA samples - NMF H values, NMF subtypes, Tsoi subtypes, TCGA subtypes

22. Supplementary Table 22: Combat batch-corrected log_2_(TPM+1) values for primary cohort samples

23. Supplementary Table 23: NMF table for primary cohort samples - NMF H values, NMF subtypes

24. Supplementary Table 24: DESeq2 results (R vs. NR, Short vs. Long OS)

25. Supplementary Table 25: DE gene CCLE/HPA expression and immune/tumor classification

26. Supplementary Table 26: DE gene co-expression matrix

27. Supplementary Table 27: DE gene HPA gene ranks, HPA gene z-scores, NMF subtype ranks, NMF subtype z-scores

28. Supplementary Table 28: Long/Short OS gene pair model performance for OS and R/NR

29. Supplementary Table 29: R/NR gene pair model performance for OS and R/NR

30. Supplementary Table 30: Metagene values for all primary cohort patients (Long/Short OS and R/NR metagenes)

31. Supplementary Table 31: Primary cohort all predictors values - raw values and z-score values

32. Supplementary Table 32: Primary cohort OS and R/NR external predictors performance (CD274, CYT etc.)

33. Supplementary Table 33: *TBX3*/*MAP4K1*/*AGER*/metagene expression in patient subsets/other sources

34. Supplementary Table 34: Cross-validation model performance

35. Supplementary Table 35: Secondary cohort log_2_(TPM+1) matrix

36. Supplementary Table 36: Secondary cohort TCB/BCB values

37. Supplementary Table 37: NMF table for secondary cohort samples - NMF H values, NMF subtypes

38. Supplementary Table 38: Secondary cohort all predictors values - raw values and z-score values

39. Supplementary Table 39: Secondary cohort gene pair model performance (OS and R/NR), HRs, models with treatment, likelihood ratio tests, binary groups

40. Supplementary Table 40: Secondary cohort external models performance analysis

41. Supplementary Table 41: Within-cohort model performance (gene pair models and external models)

42. Supplementary Table 42: *TBX3* GSEA data

---

**Supplementary Figure 1.**
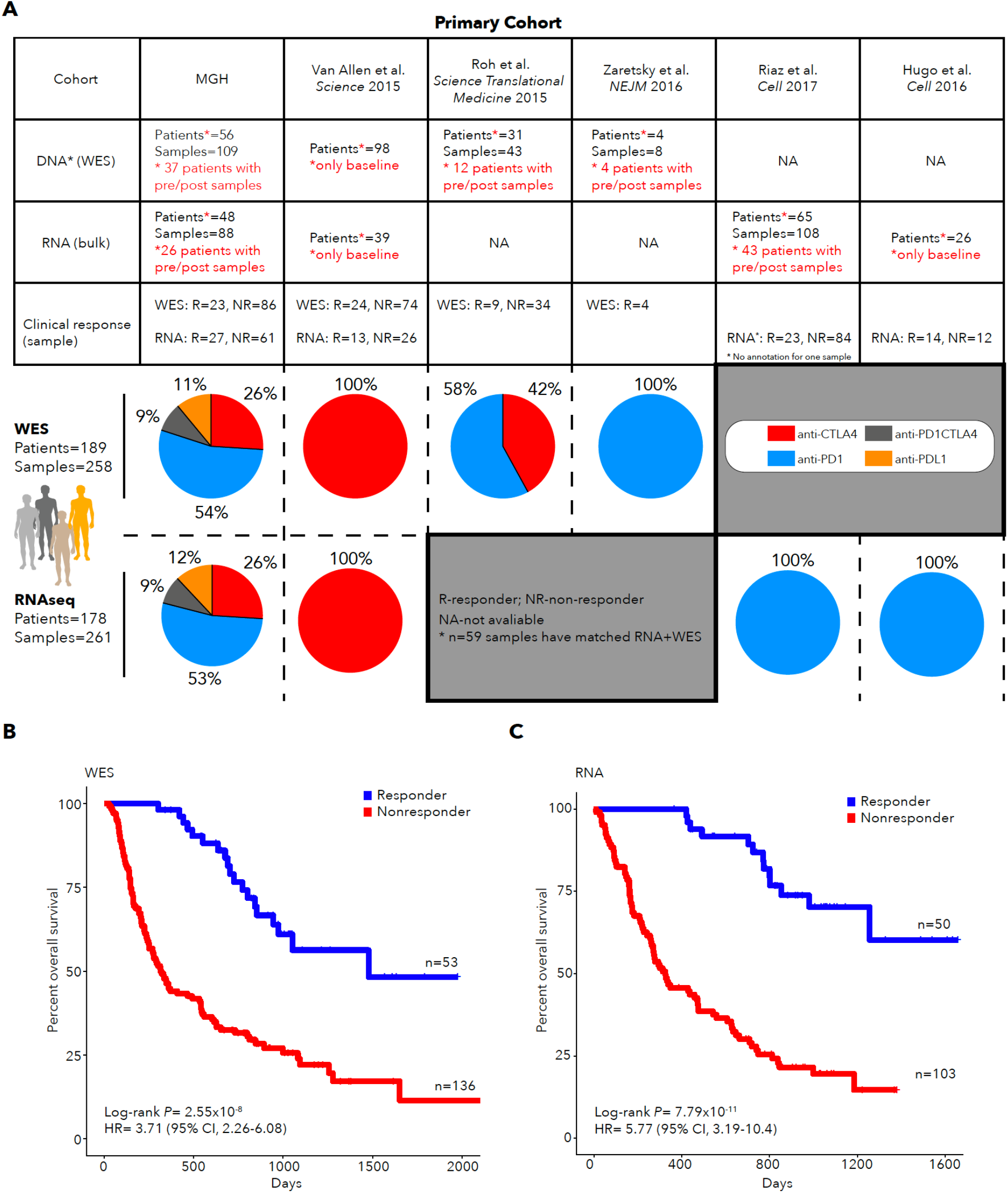
Primary cohort composition in meta-analysis. **A.** Six cohorts were included in the primary cohort meta-analysis. Table indicated cohorts for which DNA and/or RNA was available. The clinical response and numbers of pre-treatment and post-treatment samples are indicated for each cohort. Treatment for patients in each cohort is indicated in the pie charts. **B-C**. Kaplan-Meier survival curve for responders and nonresponders in the primary cohort for DNA (WES) **(B)** and RNA **(C)** samples is shown. One patient in the Hugo cohort had response data but no survival data.

**Supplementary Figure 2.**
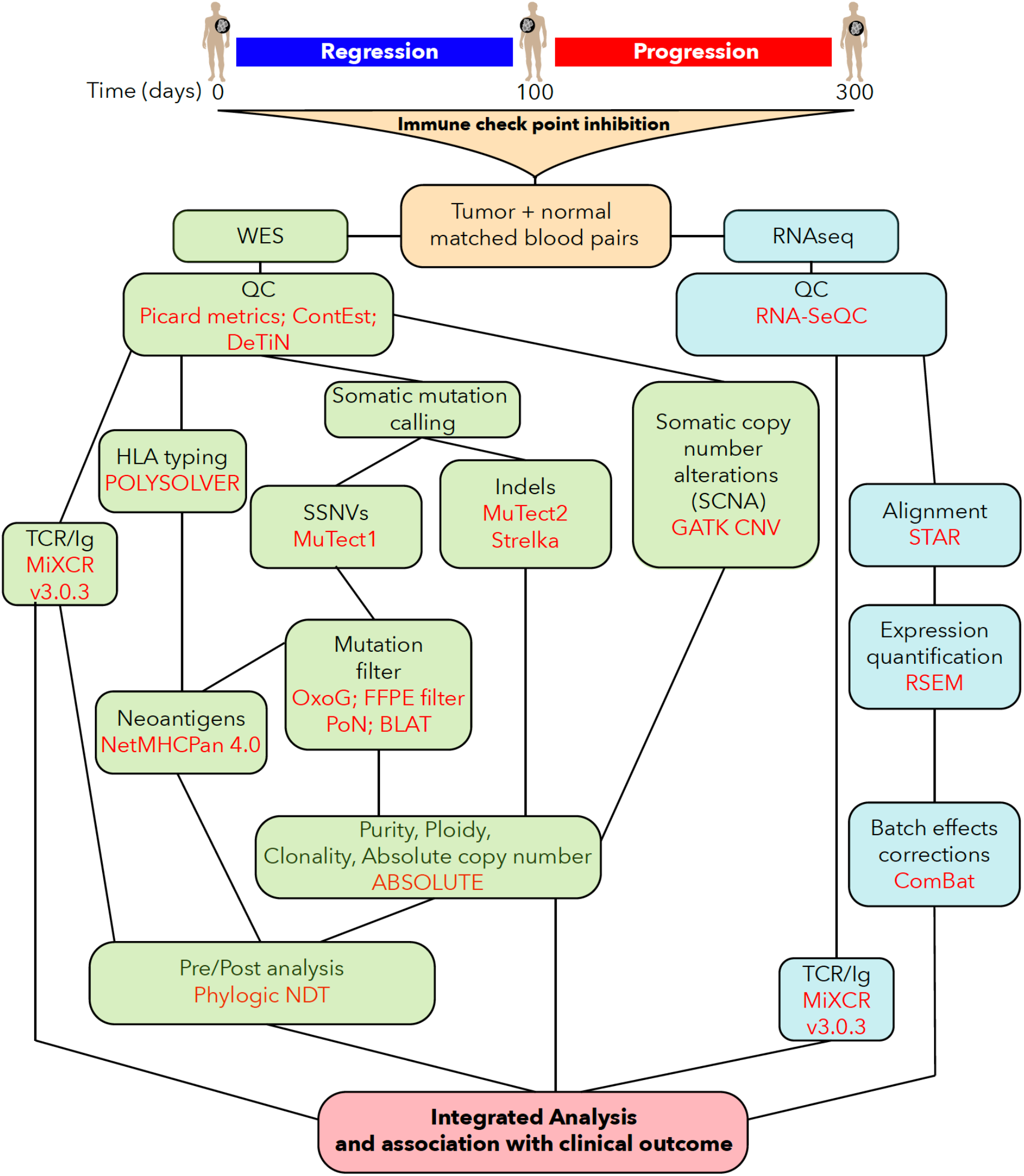
Analysis workflow. Flow chart of analysis pipeline used to process DNA (WES) and RNA-seq data in this study.

**Supplementary Figure 3.**
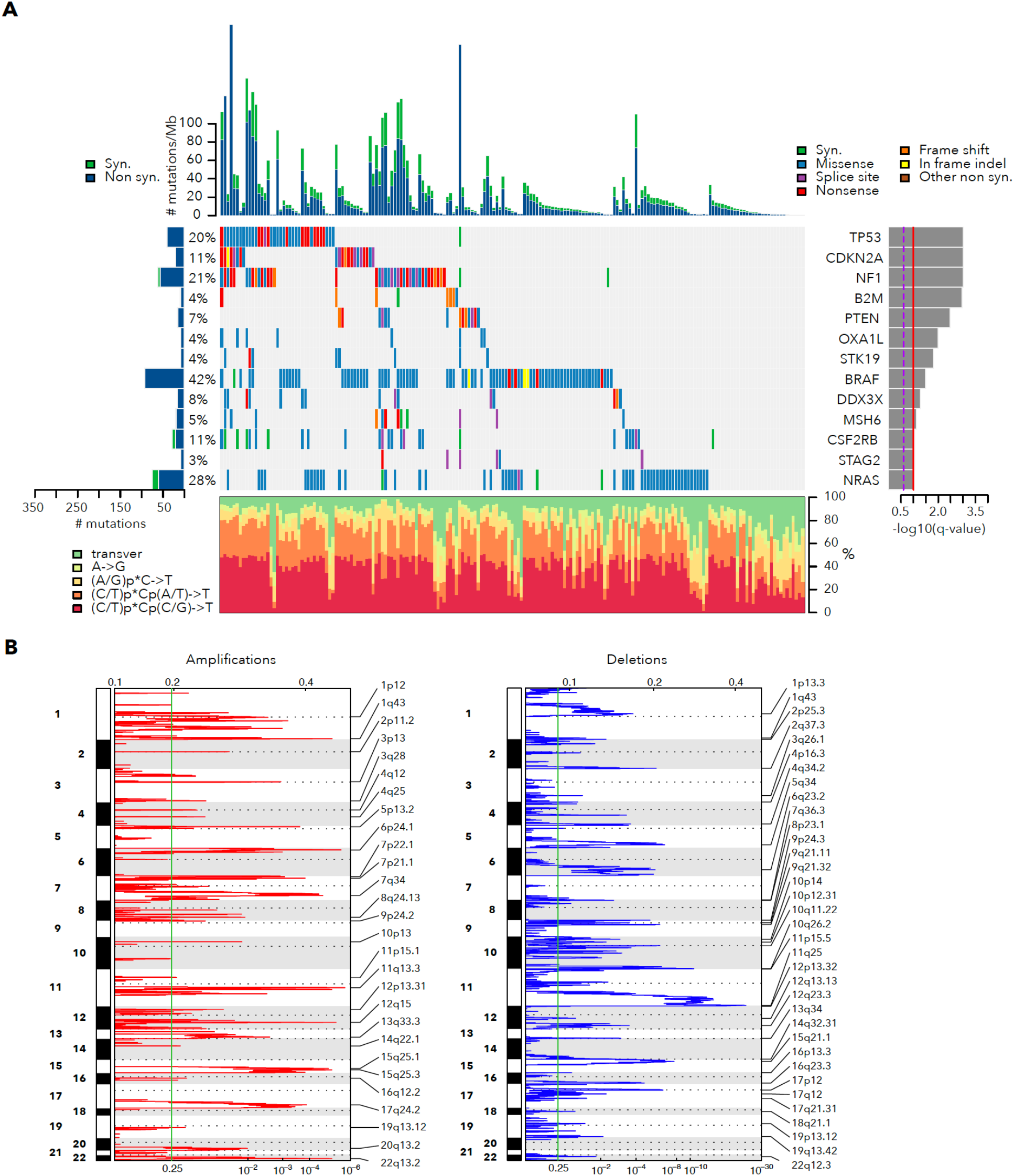
Primary cohort WES analysis. **A.** Significantly mutated genes identified with MutSig2CV in the primary meta-analysis cohort (n=189). Genes with median TPM<1 were filtered out. B. Significant amplifications (red) and deletions (blue) identified with GISTIC2.

**Supplementary Figure 4.**
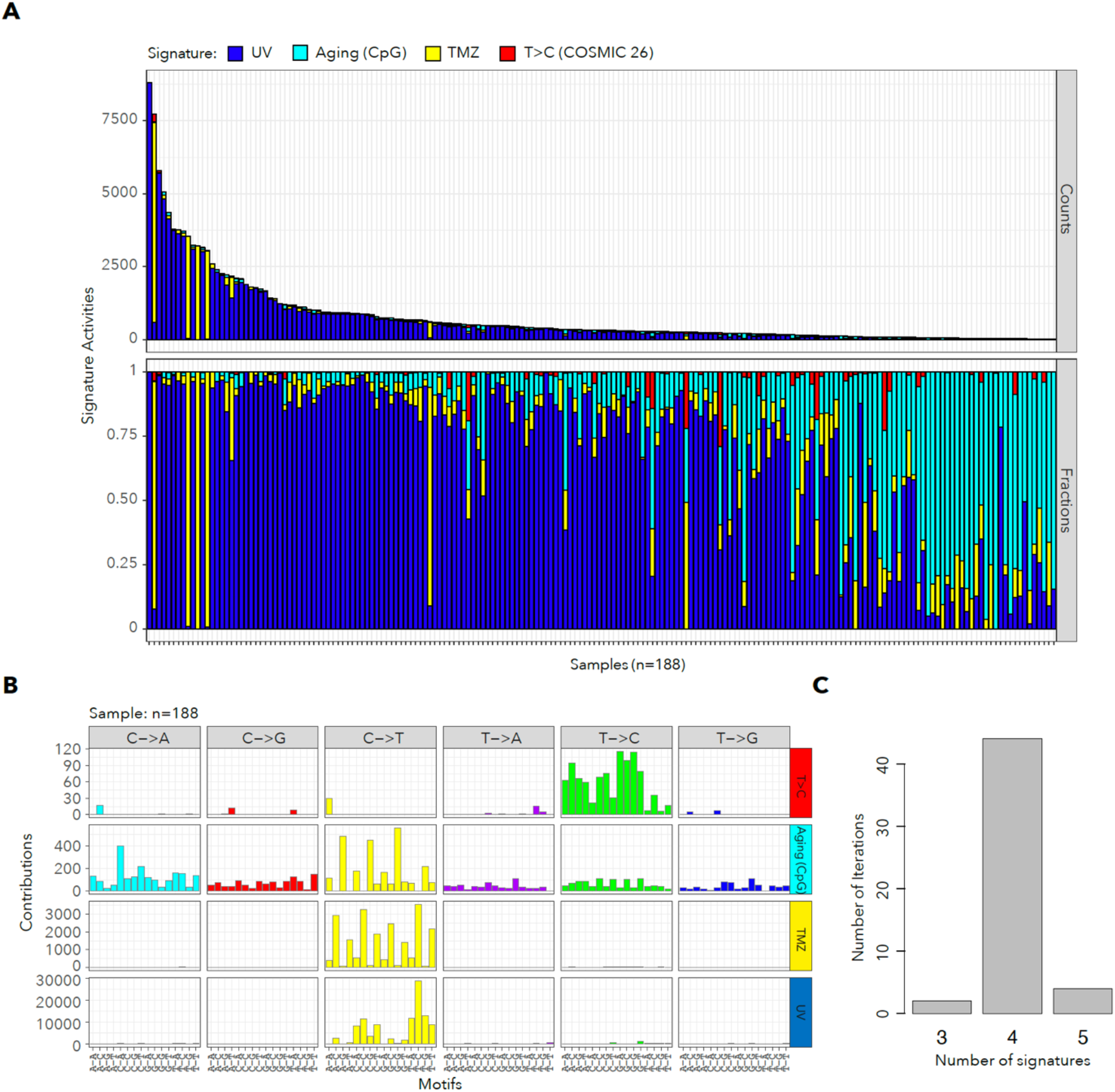
Mutational signatures. **A.** Counts and fraction of mutations from each of four mutational signatures identified using SignatureAnalyzer Bayesian NMF. One sample with MSI was removed. **B.** Sequence contexts for the four mutational signatures with the number of mutations in each context. **C.** Number of Bayesian NMF runs with K chosen signatures out of a total of 50 runs. The solution with K=4 signatures was most frequent.

**Supplementary Figure 5.**
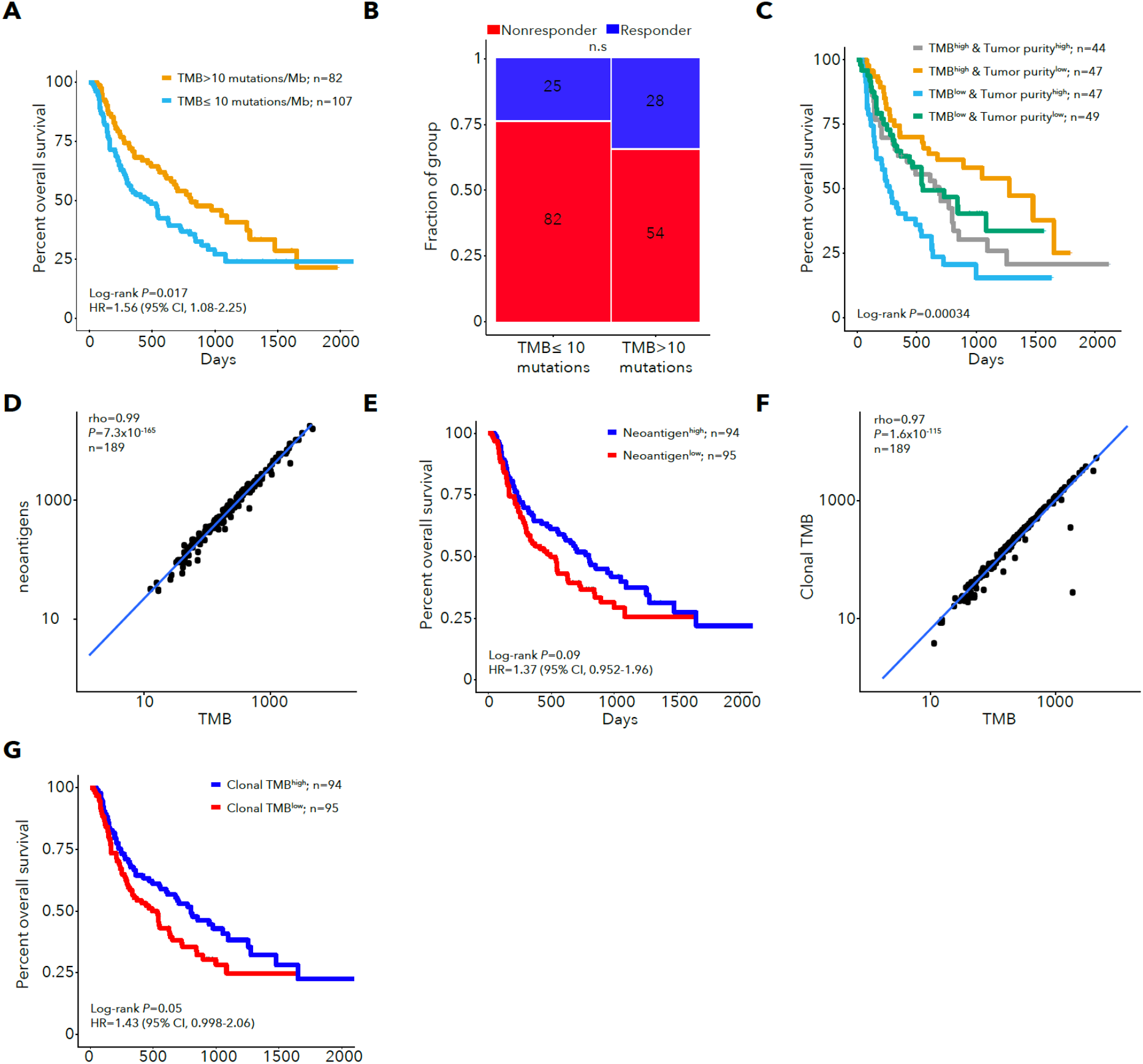
TMB, neoantigens and tumor purity. **A.** Kaplan-Meier survival curve for TMB high and low subgroups using TMB=10 mutations/Mb as a threshold (rather than the median). **B.** Response for patients with TMB over or under 10 mutations/Mb. **C.** Kaplan-Meier survival curve for all four subgroups of TMB (using the median threshold) and tumor purity. D. Spearman correlation of TMB with the number of neoantigens. **E.** Kaplan-Meier survival curve for patients with neoantigen burden above or below median. **F.** Correlation between clonal non-silent mutation burden (Clonal TMB) and TMB. **G.** Kaplan-Meier survival curve for patients with clonal TMB above or below median.

**Supplementary Figure 6.**
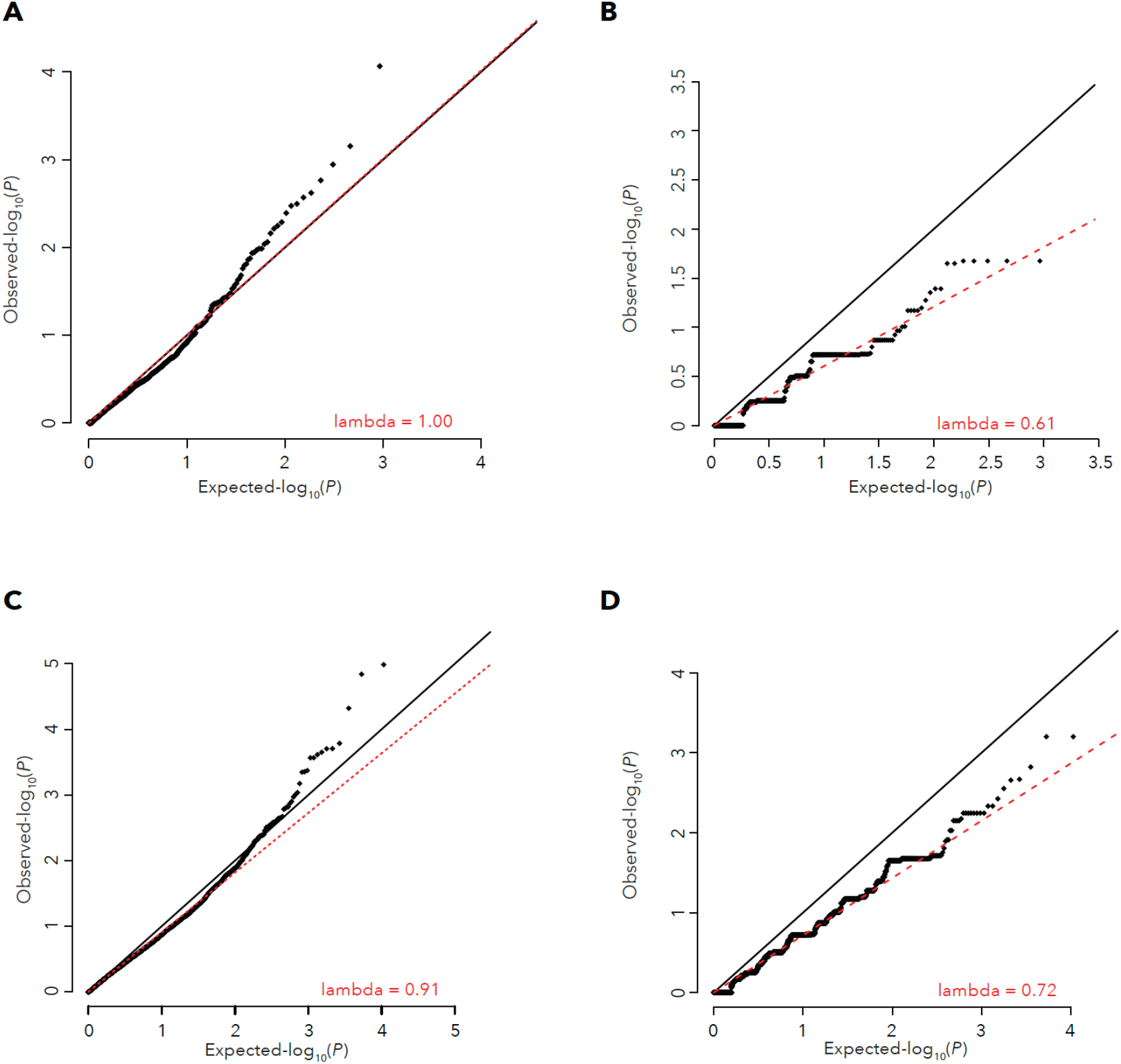
Single gene mutation models. **A.** qq plot for p values of the addition of each gene’s mutation status to TMB in a Cox model for survival using loss-of-function mutation status for genes with 3 or more loss-of-function mutations. **B.** qq plot for p values of the addition of each gene’s mutation status to TMB in a logistic regression model for response status using loss-of-function mutation status for genes with 3 or more loss-of-function mutations. **C.** qq plot for p values of the addition of each gene’s mutation status to TMB in a Cox model for survival using non-synonymous mutation status for genes with 3 or more non-synonymous mutations. **D.** qq plot for p values of the addition of each gene’s mutation status to TMB in a logistic regression model for response status using non-synonymous mutation status for genes with 3 or more non-synonymous mutations.

**Supplementary Figure 7.**
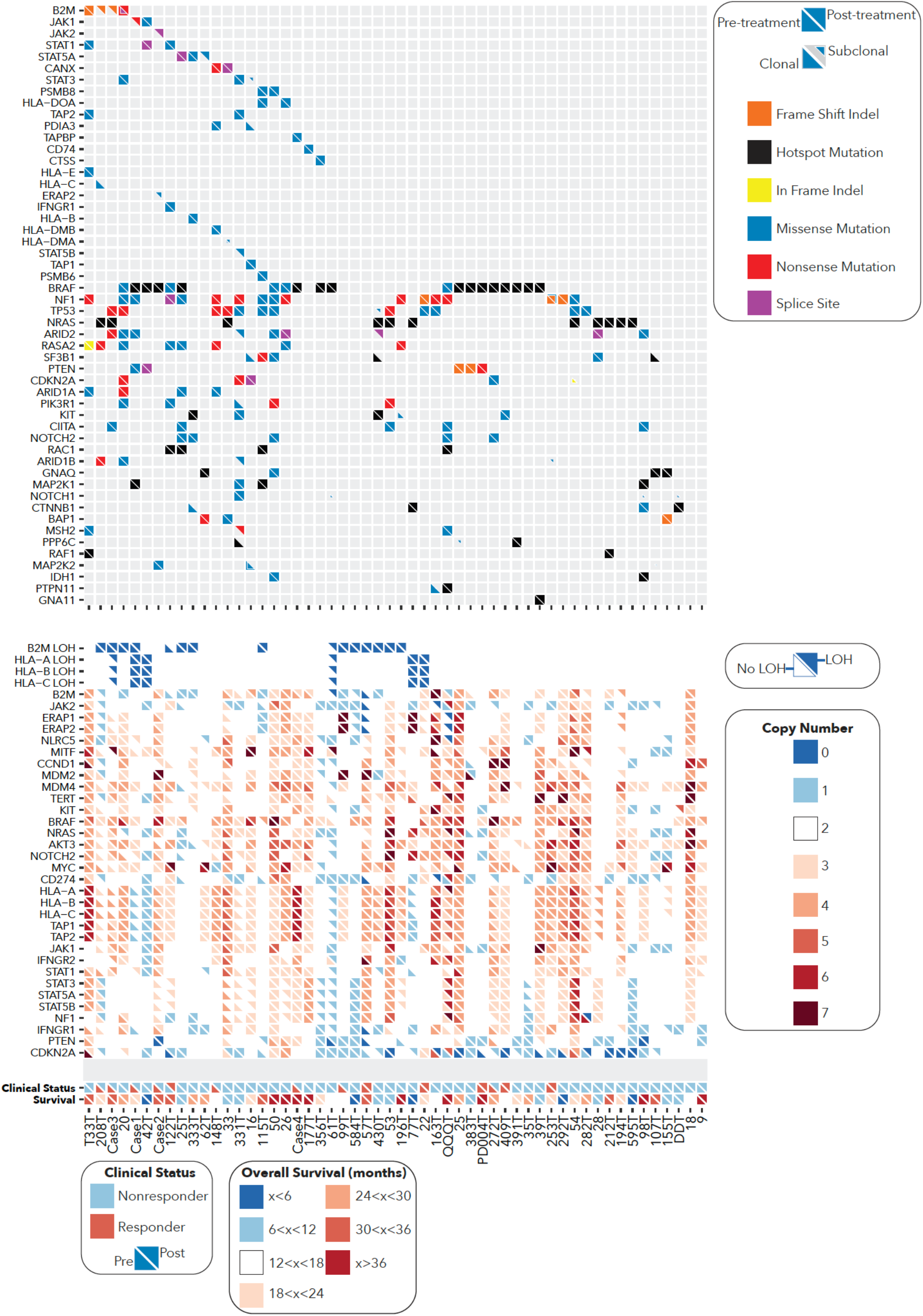
Comparison between paired pre-treatment and post-treatment biopsies. Upper panel shows a plot of mutations for selected genes in matched pre-treatments and post-treatment biopsies. The lower panel shows the integer copy number and LOH status for selected genes. Mutation clonality is represented by the area of each triangle in the upper panel. Lower triangles indicate mutations or copy number pre-treatment and upper triangles indicate mutations or copy number post-treatment. In the lower panel, white indicates absence of LOH in the first four rows and a copy number of 2 in the remaining rows. Patient response characteristics and survival are shown in the bottom two rows.

**Supplementary Figure 8.**
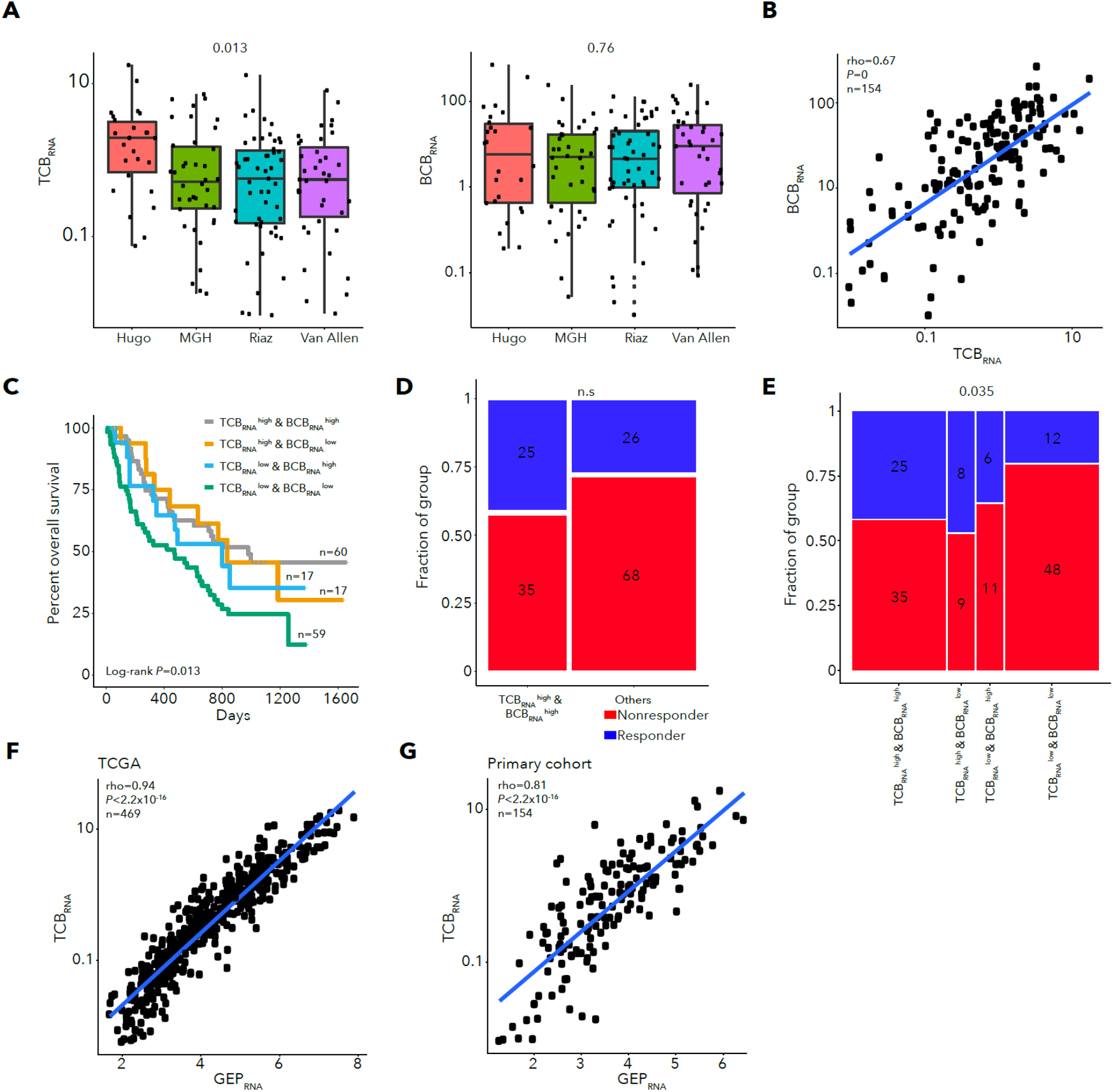
Performance of TCBRNA and BCBRNA models. **A.** TCBRNA (left) and BCBRNA (right) for each cohort with Kruskal-Wallis test p values. **B.** Correlation between TCBRNA and BCBRNA for primary cohort samples. **C.** Kaplan-Meier survival curve for all TCBRNA and BCBRNA high and low subgroups. **D.** Response for TCBRNA high, BCBRNA high subgroup vs. others. **E.** Response for all TCBRNA and BCBRNA high and low subgroups. **F.** Correlation between TCBRNA and GEP in TCGA melanoma samples. **G.** Correlation between TCBRNA and GEP in primary cohort samples.

**Supplementary Figure 9.**
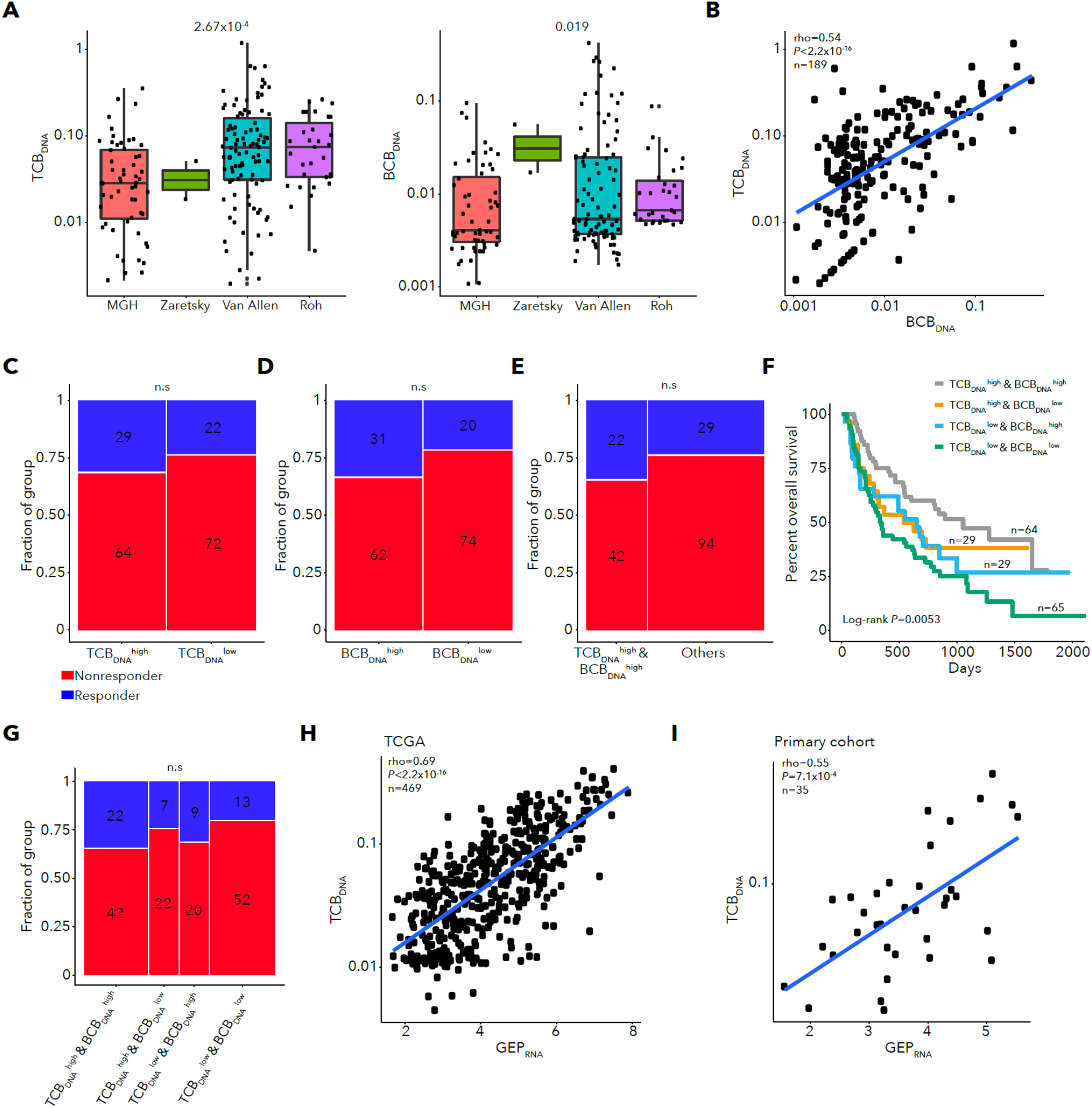
Performance of TCBDNA and BCBDNA models. **A.** TCBDNA (left) and BCBDNA (right) for each cohort with Kruskal-Wallis test p values. Due to cohort differences, we classified samples as above or below median TCBDNA or BCBDNA within each cohort. **B.** Correlation between TCBDNA and BCBDNA for primary cohortsamples. **C.** Response for TCBDNA high and low subgroups. D. Response for BCBDNA high and low subgroups. **E.** Response for TCBDNA high, BCBDNA high subgroup vs. others. **F.** Kaplan-Meier survival curve for all TCBDNA and BCBDNA high and low subgroups. **G.** Response for all TCBDNA and BCBDNA high and low subgroups. **H.** Correlation between TCBDNA and GEPRNA in TCGA melanoma samples. **I.** Correlation between TCBDNA and GEPRNA in primary cohort samples, for samples with DNA and RNA extracted from the same location in the tumor.

**Supplementary Figure 10.**
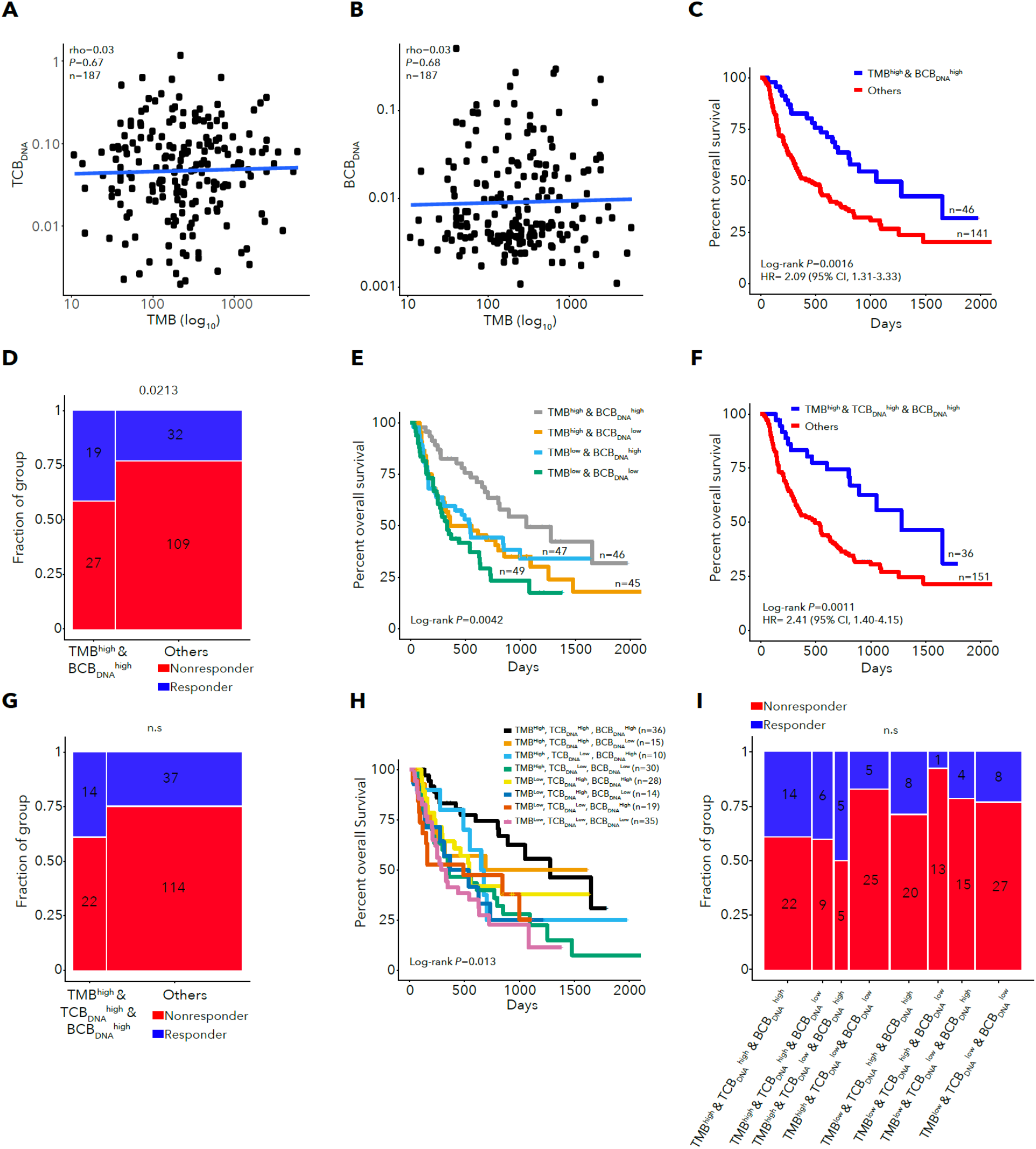
Performance of integrative DNA-based models for survival and response prediction. **A.** Correlation between TMB and TCBDNA. **B.** Correlation between TMB and BCBDNA. C. Kaplan-Meier survival curve for TMB high, BCBDNA high subgroup vs. other patients. **D.** Response for TMB high, BCBDNA high subgroup vs. other patients. **E.** Kaplan-Meier survival curve for all combined TMB and BCBDNA subgroups. **F.** Kaplan-Meier survival curve for TMB high, TCBDNA high, BCBDNA high subgroup vs. other patients. **G.** Response for TMB high, TCBDNA high, BCBDNA high subgroup vs. other patients. **H.** Kaplan-Meier survival curve for all combined TMB, TCBDNA, and BCBDNA subgroups. **I.** Response for all combined TMB, TCBDNA, and BCBDNA subgroups.

**Supplementary Figure 11.**
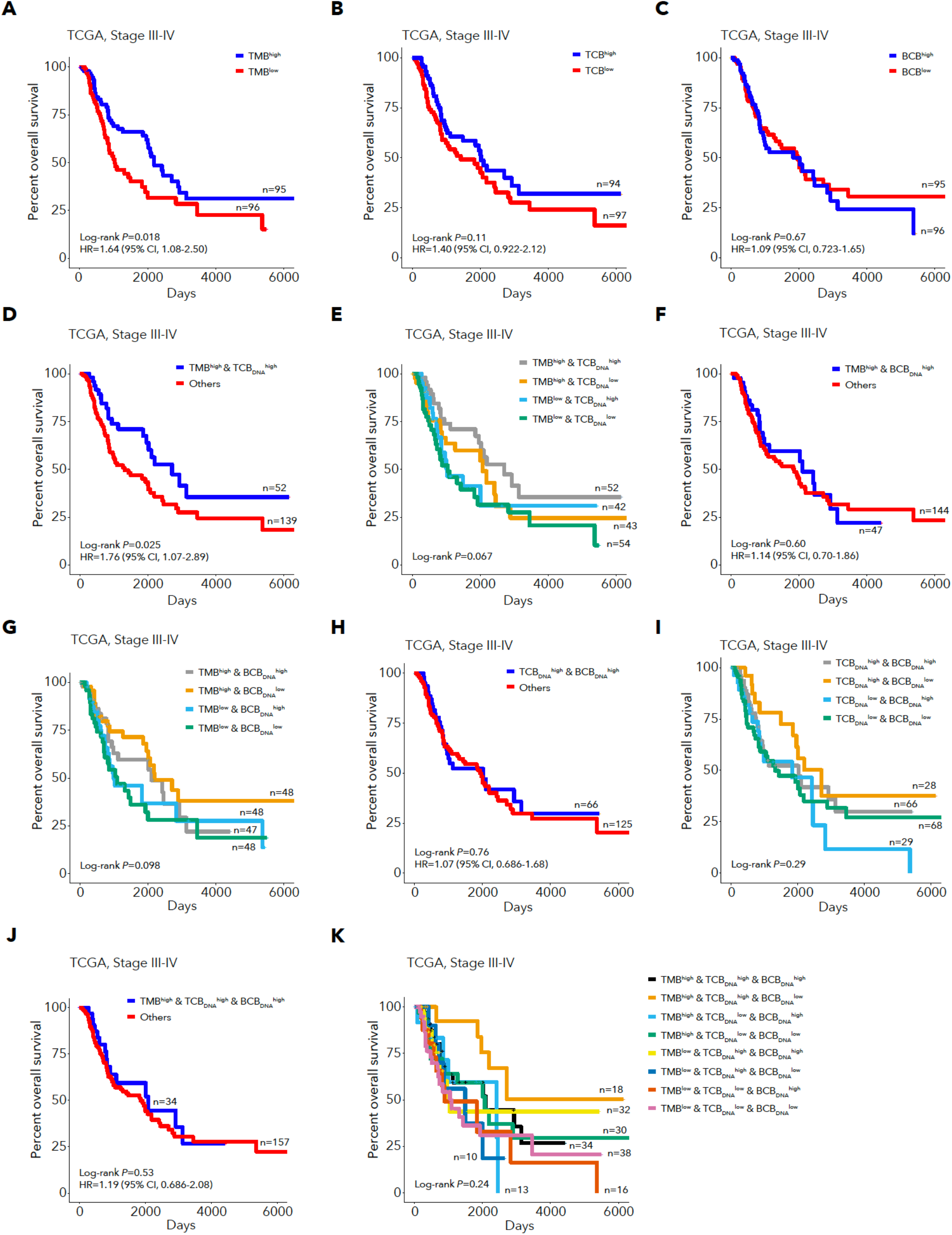
Performance of integrative DNA-based models for survival and response prediction for TCGA melanoma stage III-IV cases. **A.** Kaplan-Meier survival curve for TMB high and low subgroups. **B.** Kaplan-Meier survival curve for TCBDNA high vs. other patients. **C.** Kaplan-Meier survival curve for BCBDNA high vs. other patients. **D.** Kaplan-Meier survival curve for TMB high, TCBDNA high subgroup vs. other patients. **E.** Kaplan-Meier survival curve for all combined TMB and TCBDNA subgroups. **F.** Kaplan-Meier survival curve for TMB high, BCBDNA high subgroup vs. other patients. **G.** Kaplan-Meier survival curve for all combined TMB and BCBDNA subgroups. **H.** Kaplan-Meier survival curve for TCBDNA high, BCBDNA high subgroup vs. other patients. **I.** Kaplan-Meier survival curve for all combined TCBDNA and BCBDNA subgroups. **J.** Kaplan-Meier survival curve for TMB high, TCBDNA high, BCBDNA high subgroup vs. other patients. **K.** Kaplan-Meier survival curve for all combined TMB, TCBDNA and BCBDNA subgroups.

**Supplementary Figure 12.**
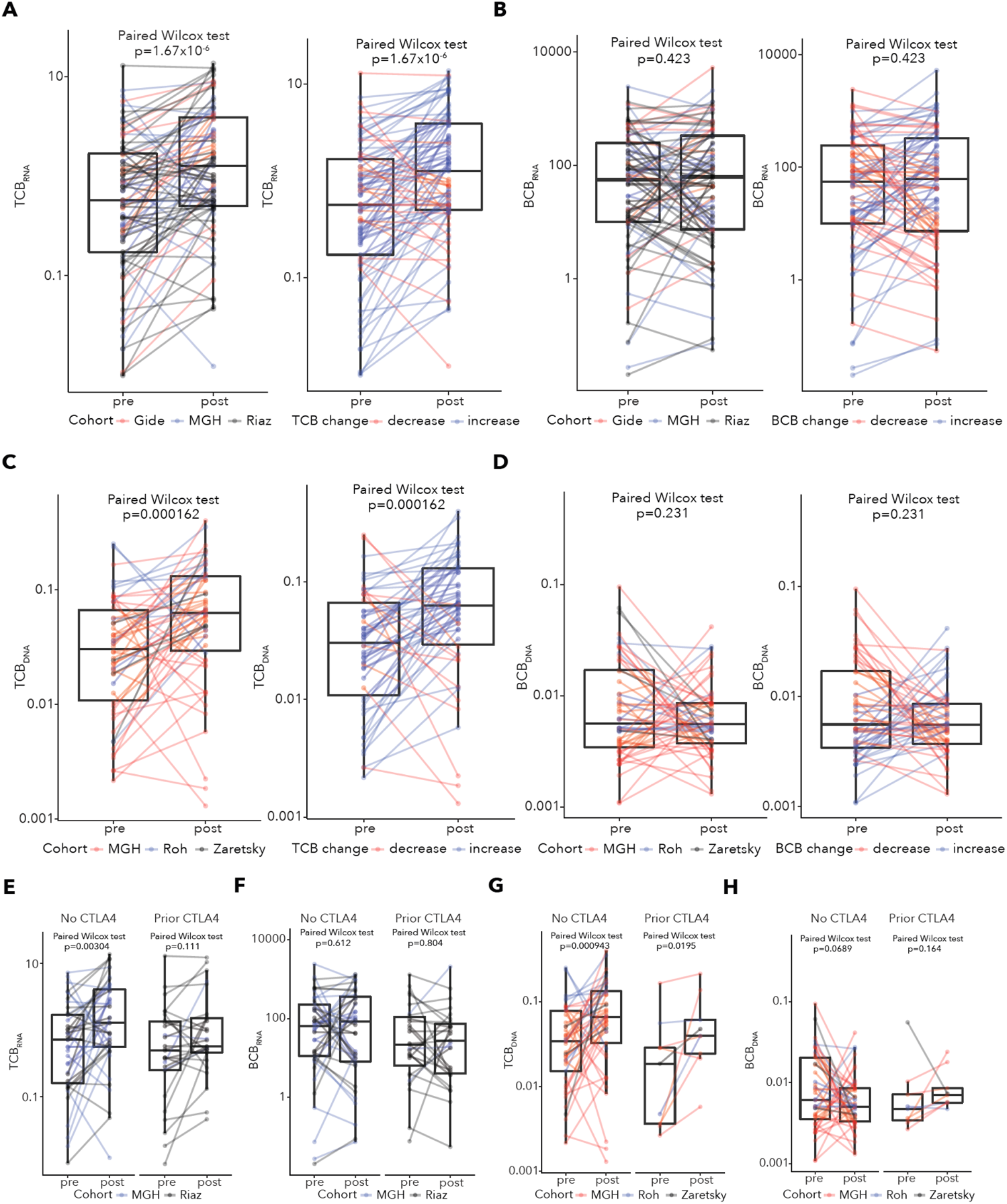
Dynamics of DNA and RNA-based TCB and BCB abundance between paired pre-treatment and post-treatment biopsies. **A.** Changes in TCBRNA between matched pre-treatment and post-treatment samples, colored by cohort (left) or decrease/increase (right). **B.** Changes in BCBRNA between matched pre-treatment and post-treatment samples, colored by cohort (left) or decrease/increase (right). **C.** Changes in TCBDNA between matched pre-treatment and post-treatment samples, colored by cohort (left) and decrease/increase (right). **D.** Changes in BCBDNA between matched pre-treatment and post-treatment samples, colored by cohort (left) and decrease/increase (right). **E.** Changes in TCBRNA between matched pre-treatment and post-treatment samples, with no prior CTLA4 therapy (left) or prior CTLA4 therapy (right). **F.** Changes in BCBRNA between matched pre-treatment and post-treatment samples, with no prior CTLA4 therapy (left) or prior CTLA4 therapy (right). **G.** Changes in TCBDNA between matched pre-treatment and post-treatment samples, with no prior CTLA4 therapy (left) or prior CTLA4 therapy (right). **H.** Changes in BCBDNA between matched pre-treatment and post-treatment samples, with no prior CTLA4 therapy (left) or prior CTLA4 therapy (right).

**Supplementary Figure 13.**
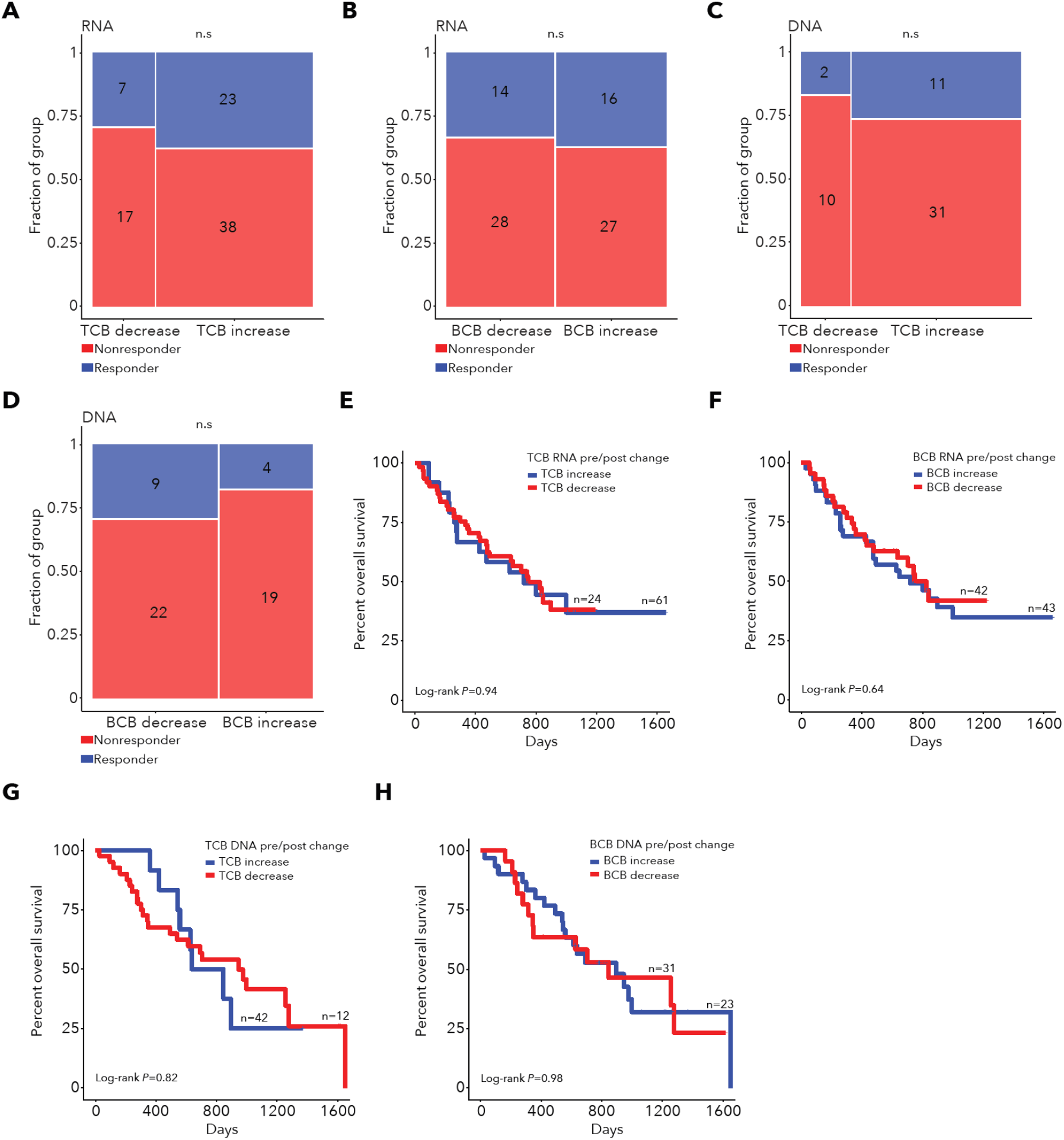
Associations between DNA and RNA-based TCB and BCB dynamics and outcomes. **A.** Response for TCBRNA decrease and TCBRNA increase groups. **B.** Response for BCBRNA decrease and BCBRNA increase groups. **C.** Response for TCBDNA decrease and TCBDNA increase groups. **D.** Response for BCBDNA decrease and BCBDNA increase groups. **E.** Kaplan-Meier survival curve for TCBRNA decrease and TCBRNA increase groups. **F.** Kaplan-Meier survival curve for BCBRNA decrease and BCBRNA increase groups. **G.** Kaplan-Meier survival curve for TCBDNA decrease and TCBDNA increase groups. **H.** Kaplan-Meier survival curve for BCBDNA decrease and BCBDNA increase groups.

**Supplementary Figure 14.**
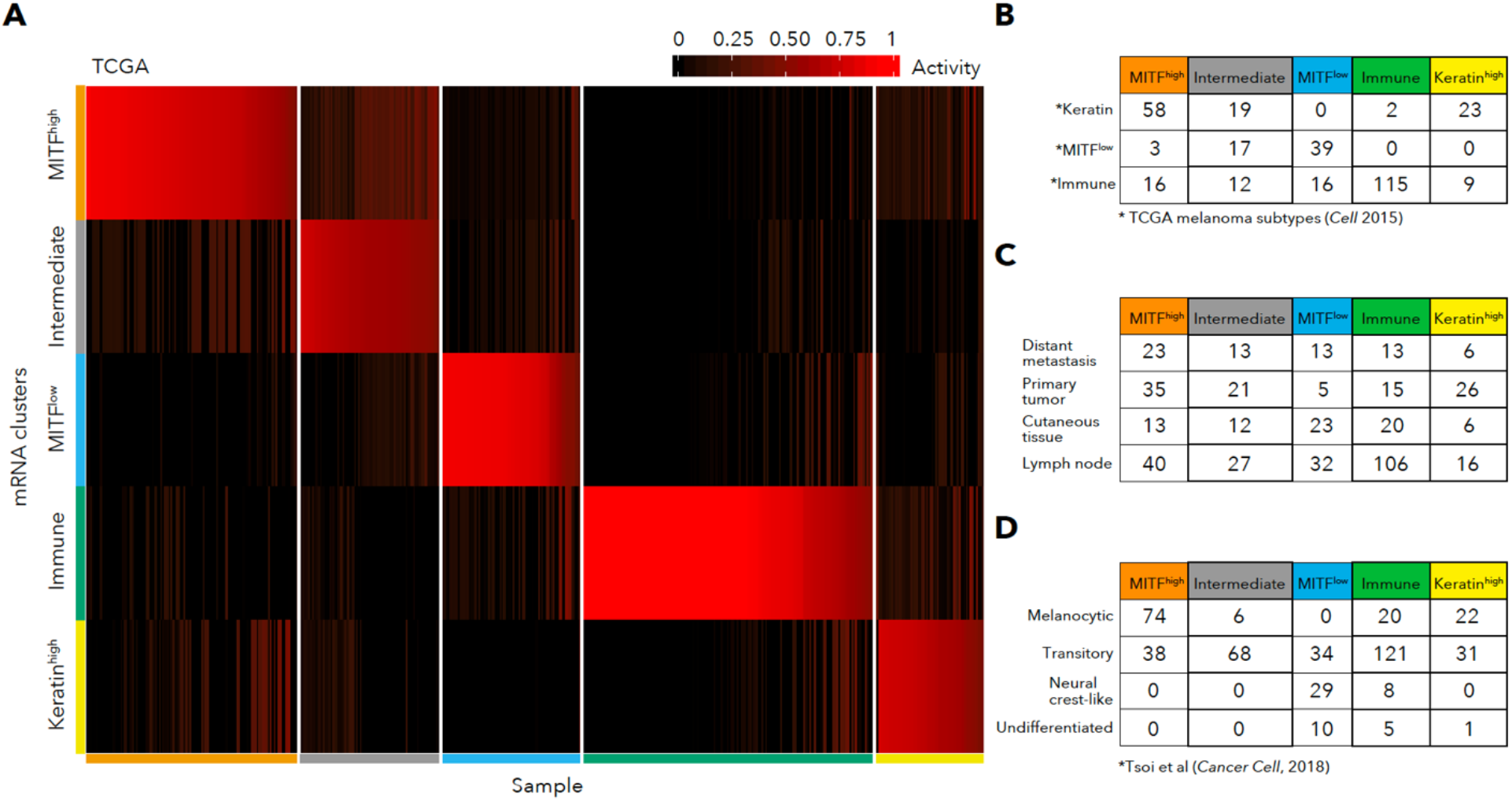
Subtypes identified using NMF clustering of TCGA melanoma RNA-seq. **A.** NMF H matrix from TCGA melanoma NMF clustering (n=469) identified 5 subtypes. Activity values indicate the probability that a sample is associated with a cluster. Samples are sorted by activity value within subtypes. **B.** Comparison of subtype membership to previously identified TCGA subtypes. **C.** TCGA biopsy locations and subtype membership for all samples. The Immune subtype is enriched for lymph node biopsy samples, but all other subtypes contain lymph node samples as well. **D.** Comparison of subtype membership to melanoma differentiation subtypes.

**Supplementary Figure 15.**
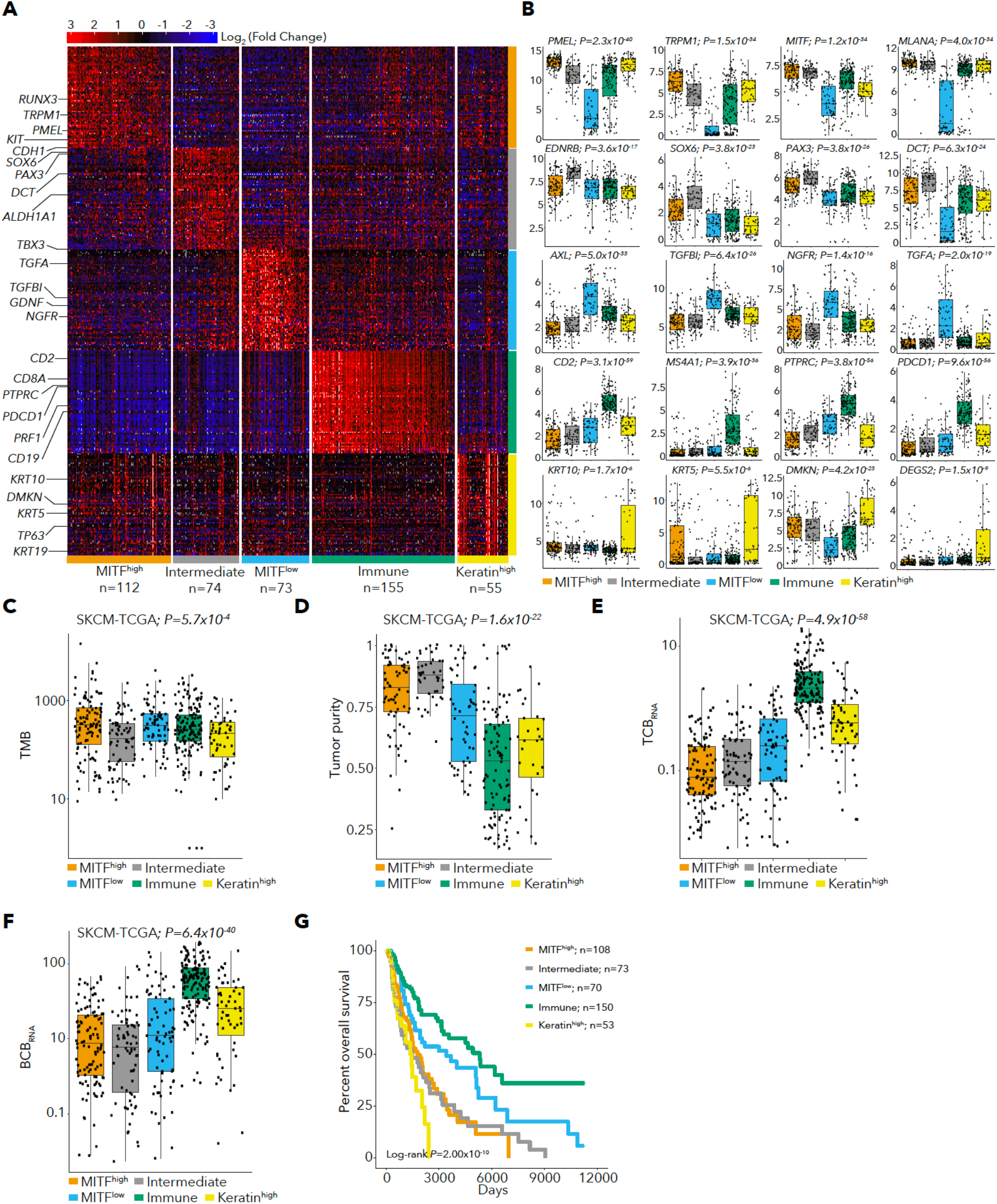
TCGA RNA-seq subtypes and their tumor related features. **A.** Heatmap of marker genes identified for each subtype in TCGA melanoma data. Initial log2(TPM+1) values were median centered to obtain log2(Fold change) values. We selected marker genes which were overexpressed in each cluster relative to all other samples. **B.** Boxplots of marker gene expression for selected genes. All genes were identified through automated marker selection and were included in the heatmap except *MITF*, *MLANA* and *AXL*. C-F. Kruskal-wallis p values for association of gene expression with subtype are displayed above plots, **(C)** TMB (log10 scale), (D) tumor purity, (E) TCBRNA and (F) BCBRNA, for TCGA samples by RNA-seq subtype. **G.** Kaplan-Meier survival curve for TCGA samples by RNA-seq subtype.

**Supplementary Figure 16.**
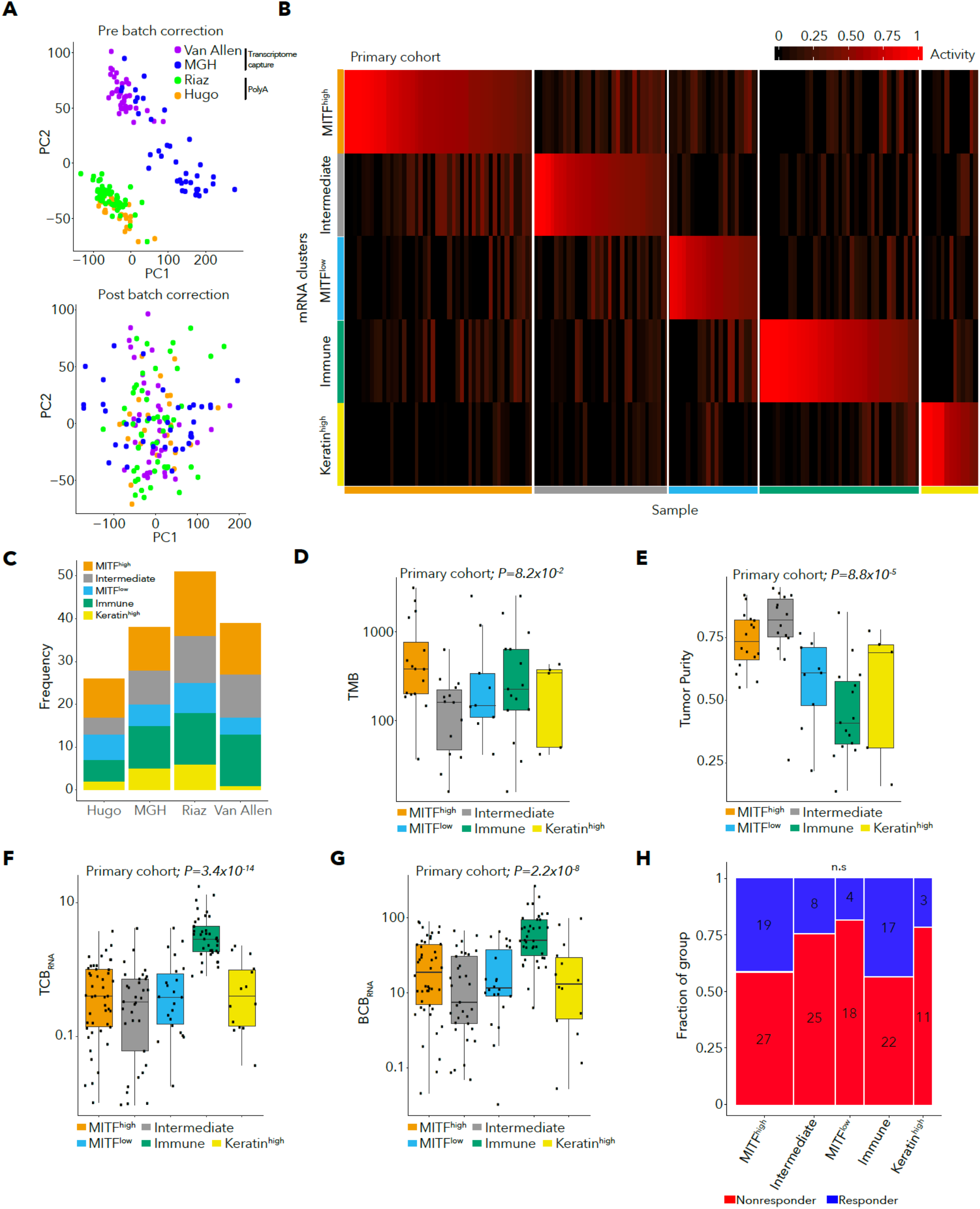
Subtype classification for primary cohort RNA-seq samples. **A.** PCA of protein coding gene log2(TPM+1) values before (upper plot) and after batch effects correction with ComBat^87^ (lower plot). Before batch effects correction, samples cluster by cohort and library preparation method (polyA selection vs. transcriptome capture). **B.** NMF H matrix for primary cohort sample subtyping using subtypes and marker genes identified in TCGA melanoma samples. **C.** Subtypes by RNA-seq sample for each sample in the primary cohort. D-G. Kruskal-wallis p values for association of gene expression with subtype are displayed above plots, (D) TMB (log10 scale), (E) tumor purity for RNA-seq samples with matched WES data, (F) TCBRNA and (G) BCBRNA. **H.** Number of responders and non-responders by subtype.

**Supplementary Figure 17.**
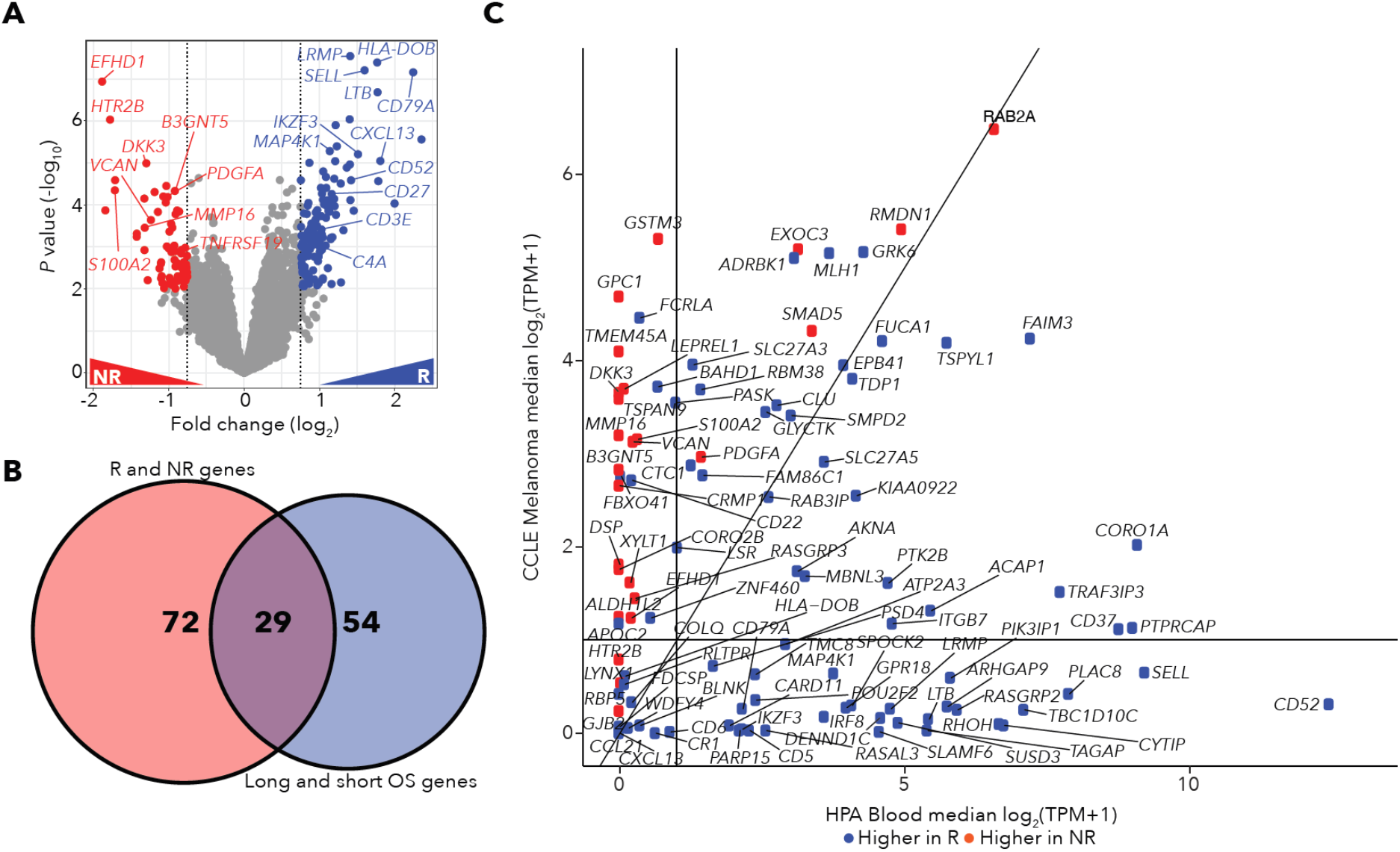
Genes associated with response in the primary cohort. **A.** Differential expression between responders (R) and non-responders (NR) in the primary cohort. **B.** Venn diagram for differentially expressed genes in the high vs. low OS and responder vs. non-responder comparisons. In total, 29 genes were differentially expressed in both analyses. **C.** Expression of responder and non-responder differentially expressed genes in melanoma CCLE cell lines and Human Protein Atlas blood cell types.

**Supplementary Figure 18.**
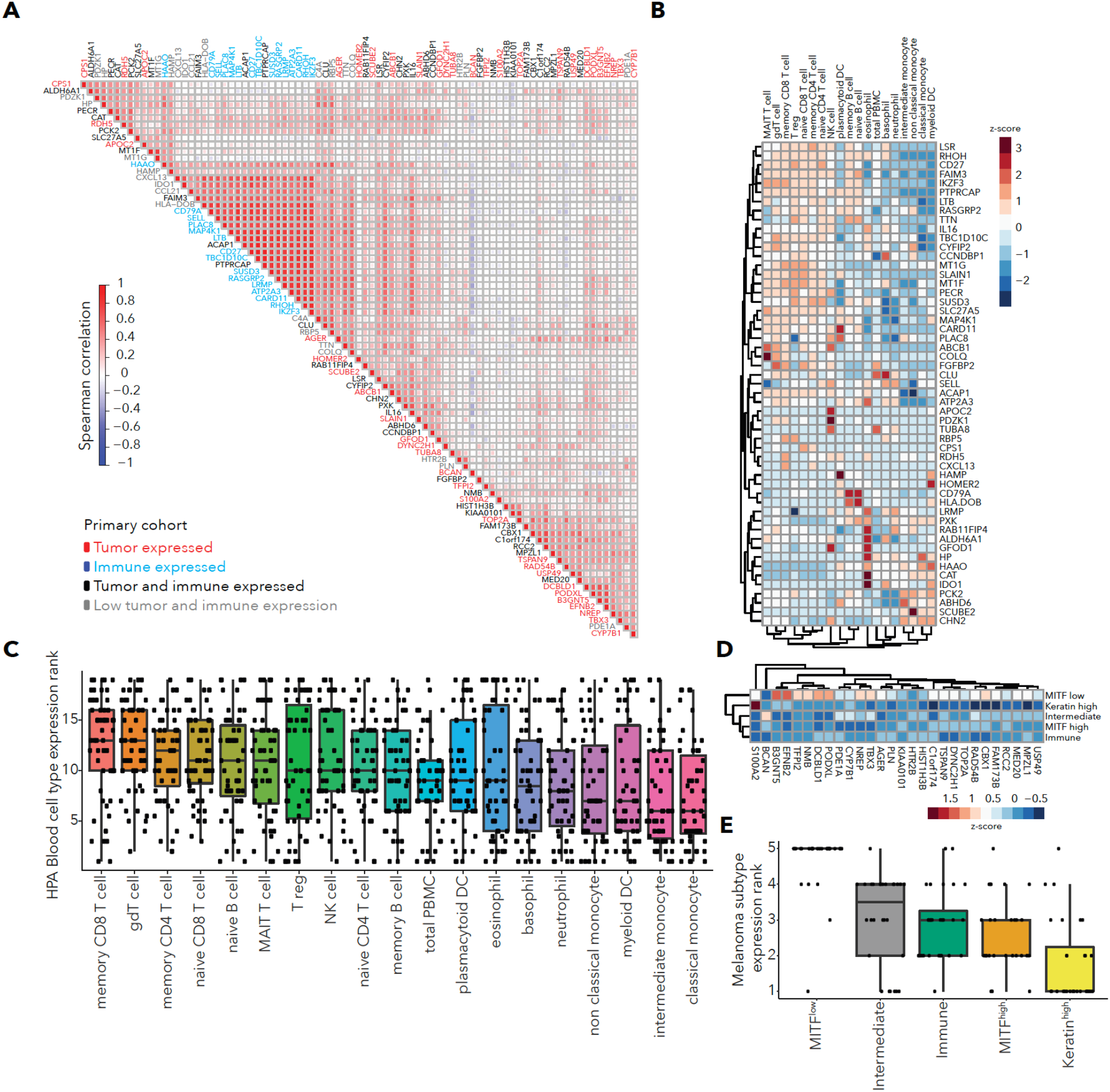
Expression patterns for long and short OS differentially expressed genes in the primary cohort. A. Co-expression for long and short OS differentially expressed genes in the primary cohort. **B.** Expression of genes overexpressed in long OS patients in Human Protein Atlas (HPA) blood cell types. **C.** Ranks of HPA cell type expression for genes overexpressed in long OS patients. **D.** Expression of genes overexpressed in short OS patients in primary cohort samples grouped by melanoma subtype. **E.** Ranks of melanoma subtype expression for genes overexpressed in short OS patients.

**Supplementary Figure 19.**
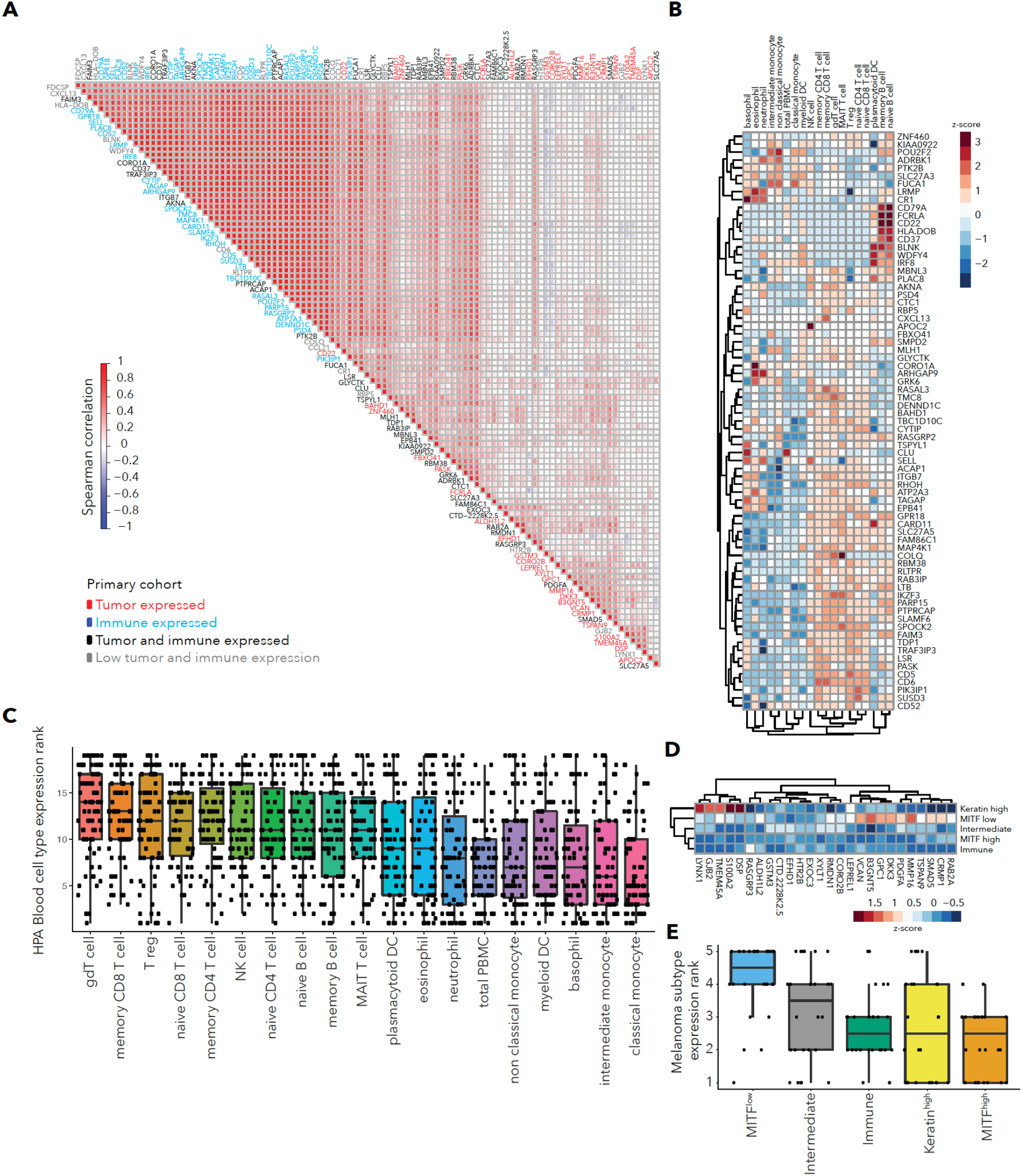
Expression patterns for responder and non-responder differentially expressed genes in the primary cohort. **A.** Co-expression for responder and non-responder differentially expressed genes in the primary cohort. **B.** Expression of genes overexpressed in responders in Human Protein Atlas (HPA) blood cell types. **C.** Ranks of HPA cell type expression for genes overexpressed in responders. **D.** Expression of genes overexpressed in non-responders in primary cohort samples grouped by melanoma subtype. **E.** Ranks of melanoma subtype expression for genes overexpressed in non-responders.

**Supplementary Figure 20.**
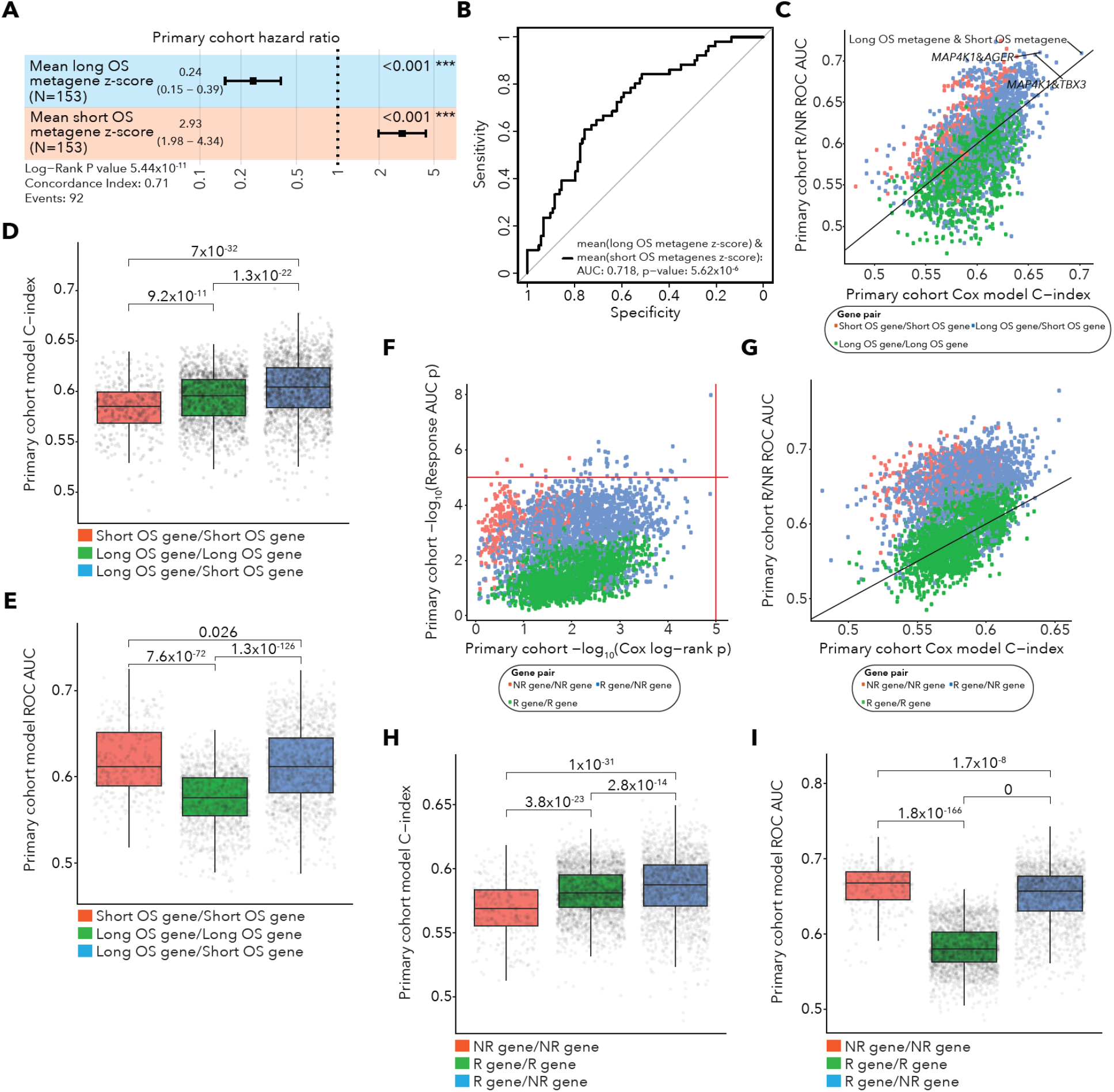
Performance of gene-pair models in the primary cohort. **A.** Forest plot for the long OS and short OS metagene pair model in the primary cohort. Error bars represent 95% confidence intervals for Cox model hazard ratio estimates. **B.** ROC curve for the long OS and short OS metagene pair model in the primary cohort. **C.** Performance for all gene pair models using genes derived from long OS vs. short OS differential expression in terms of survival and response predictions in the primary cohort based on survival C-index and response AUC. Each point represents one gene pair model, and points are colored by the gene pair type (Short OS/Short OS gene pair- red, Long OS/Long OS gene pair- green or Short OS/Long OS gene pair- blue). **D.** Survival C-index for gene pair models using genes derived from long OS vs. short OS differential expression by gene pair type. **E.** Response AUC for gene pair models using genes derived from long OS vs. short OS differential expression by gene pair type. **F.** Performance for all gene pair models using genes derived from responder (R) vs. non-responder (NR) differential expression in terms of significance of survival and response predictions in the primary cohort. Survival prediction significance is the Cox proportional hazards model Log-Rank P value and response prediction significance is the Logistic regression AUC p value. Each point represents one gene pair model, and points are colored by the gene pair type (NR/NR gene pair- red, R/R gene pair- blue or NR/R gene pair- green). **G.** Performance for all gene pair models using genes derived from R vs. NR differential expression in terms of effect size of survival and response predictions (C-index and AUC respectively). **H.** Survival C-index for gene pair models using genes derived from R vs. NR differential expression by gene pair type. **I.** Response AUC for gene pair models using genes derived from R vs. NR differential expression by gene pair type.

**Supplementary Figure 21.**
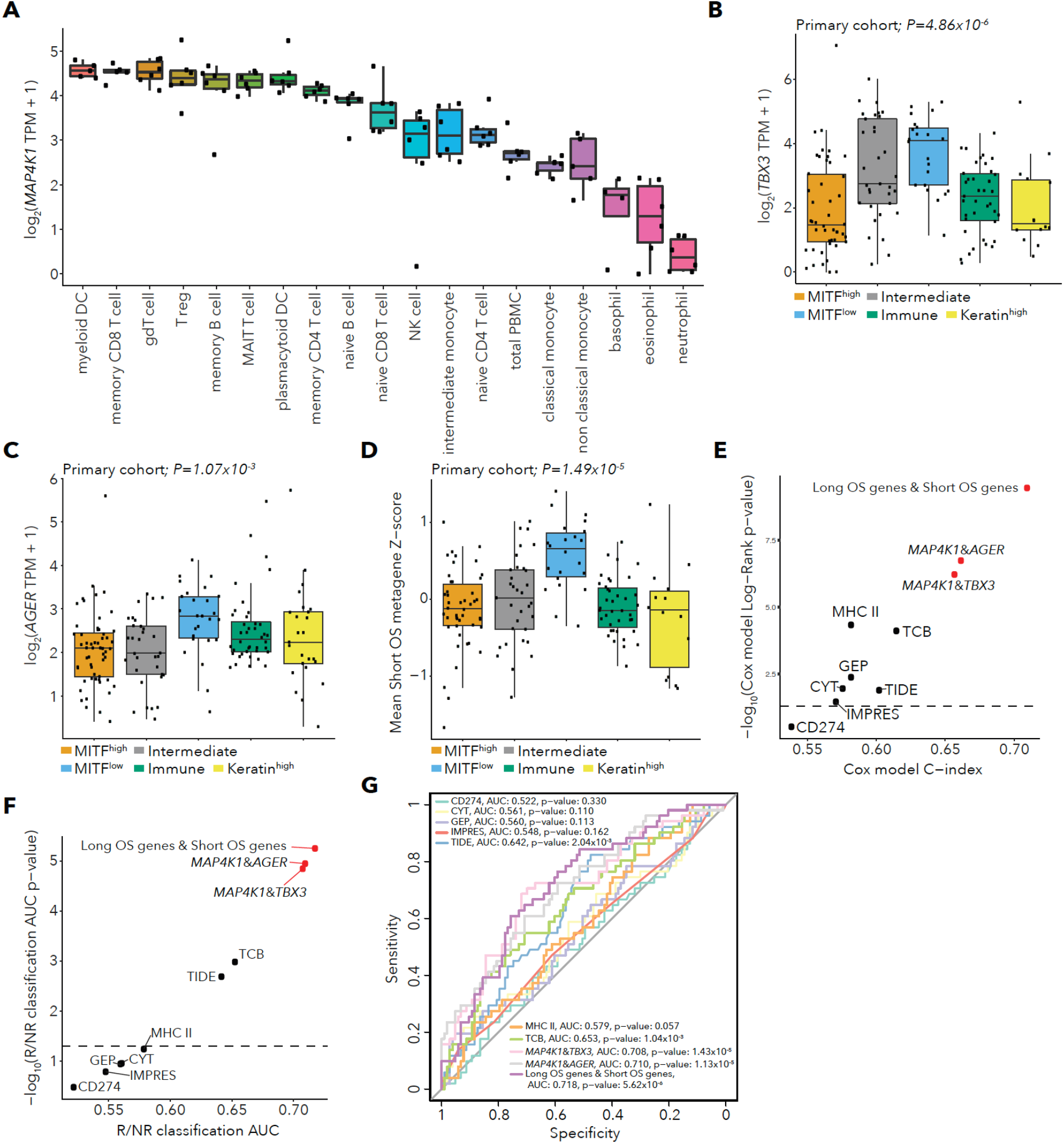
Expression of genes from top gene pair models and gene pair model performance. **A.** Expression of *MAP4K1* in Human Protein Atlas (HPA) cell types. **B.** Expression of *TBX3* in the primary cohort by melanoma subtype with Kruskal-Wallis test P-value. **C.** Expression of *AGER* in the primary cohort by melanoma subtype with Kruskal-Wallis test P-value. **D.** Expression of the short OS metagene in the primary cohort by melanoma subtype with Kruskal-Wallis test P-value. **E.** Performance of melanoma immunotherapy survival models in the primary cohort in terms of C-index and Cox model log-rank P-value. **F.** Performance of melanoma immunotherapy response models in the primary cohort in terms of AUC and AUC P-value. **G.** ROC curves for response classification for all models in the primary cohort.

**Supplementary Figure 22.**
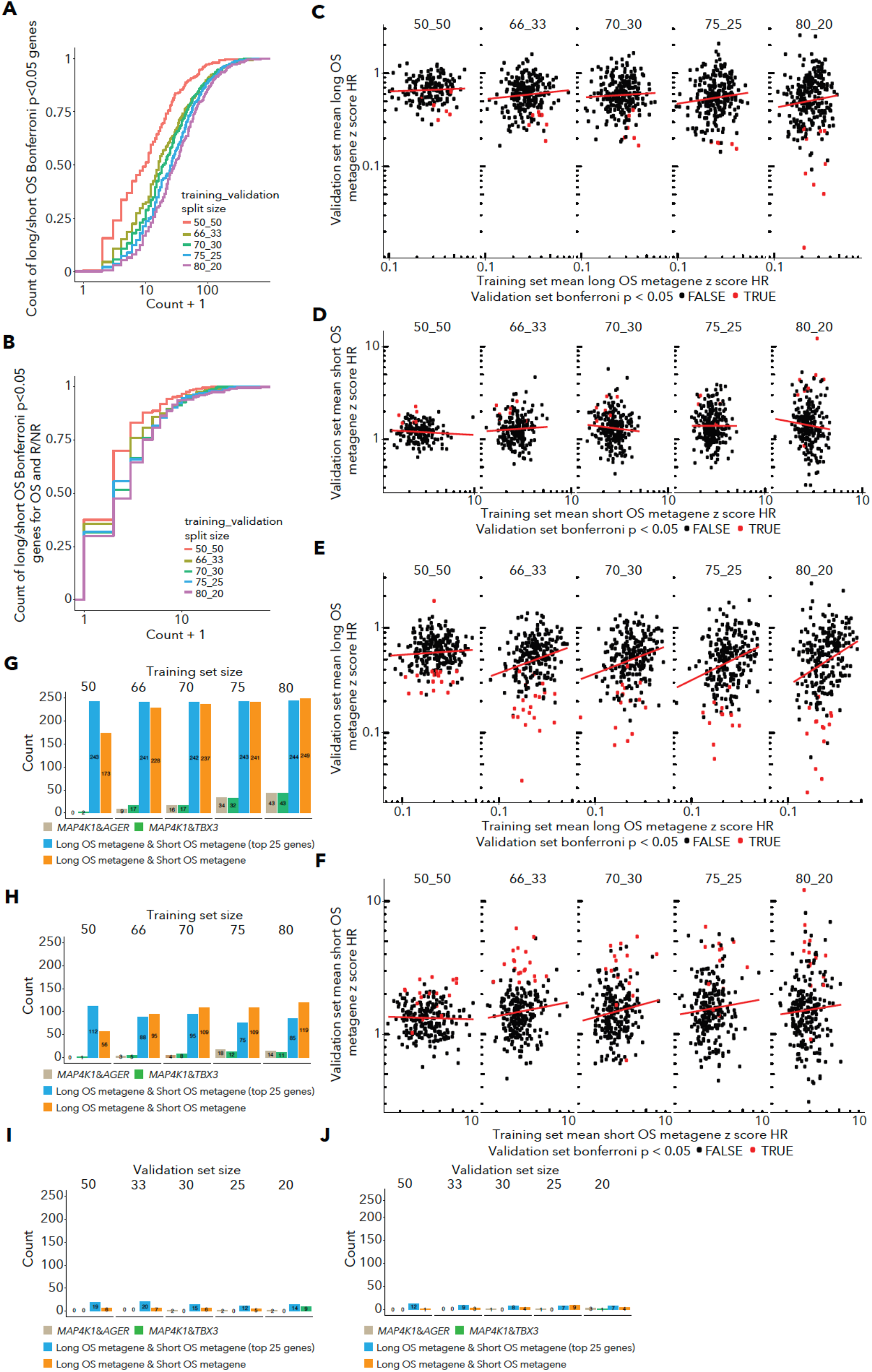
Cross-validation of gene pair model discovery and validation in the primary cohort. **A.** Empirical cumulative distribution of the number of gene pairs discovered with Bonferroni p<0.05 for association with survival in training sets in the cross validation using genes differentially expressed between patients with long and short OS (DESeq q<0.05) within the training set. Each line represents a different training and validation set split size, with 250 cross validation training/validation splits per split size. **B.** Empirical cumulative distribution of the number of gene pairs discovered with Bonferroni p<0.05 for association with survival and response in training sets in the cross validation using genes differentially expressed between patients with long and short OS (DESeq q<0.05) within the training set. Each line represents a different training and validation set split size, with 250 cross validation training/validation splits per split size. C-D. Performance of the long OS metagene & short OS metagene model in cross-validation training and validation sets. Each point represents the hazard ratio (HR) of the long OS metagene **(C)** and short OS metagene (D) discovered in that training/test set split. In training/validation splits, the metagenes are composed of the genes that were differentially expressed between long and short OS patients with DESeq q<0.05 in the training set samples. Panels represent different split sizes and red lines are linear regressions. Points are colored by whether the metagene had Bonferroni p<0.05 for survival association in the validation set. E-F. Performance of the the long OS metagene & short OS metagene model in cross-validation training and validation sets where metagenes were defined using top 25 long OS or short OS genes ranked by DESeq p value. G-H. Frequency of selected gene pairs with Bonferroni p<0.05 for association with survival (G) or Bonferroni p<0.05 for association with survival and response (H) in training sets. Training set sizes are listed above with 250 training/validation splits for each split size. Metagene pair models using the top 25 genes ranked by DESeq p value or the genes with DESeq q<0.05 in the training set are shown separately. I-J. Frequency of selected gene pairs with Bonferroni p<0.05 for association with survival (I) or Bonferroni p<0.05 for association with survival and response (J) in validation sets. Validation set sizes are listed above with 250 training/validation splits for each split size. Metagene pair models using the top 25 genes ranked by DESeq p value or the genes with DESeq q<0.05 in the training set are shown separately.

**Supplementary Figure 23.**
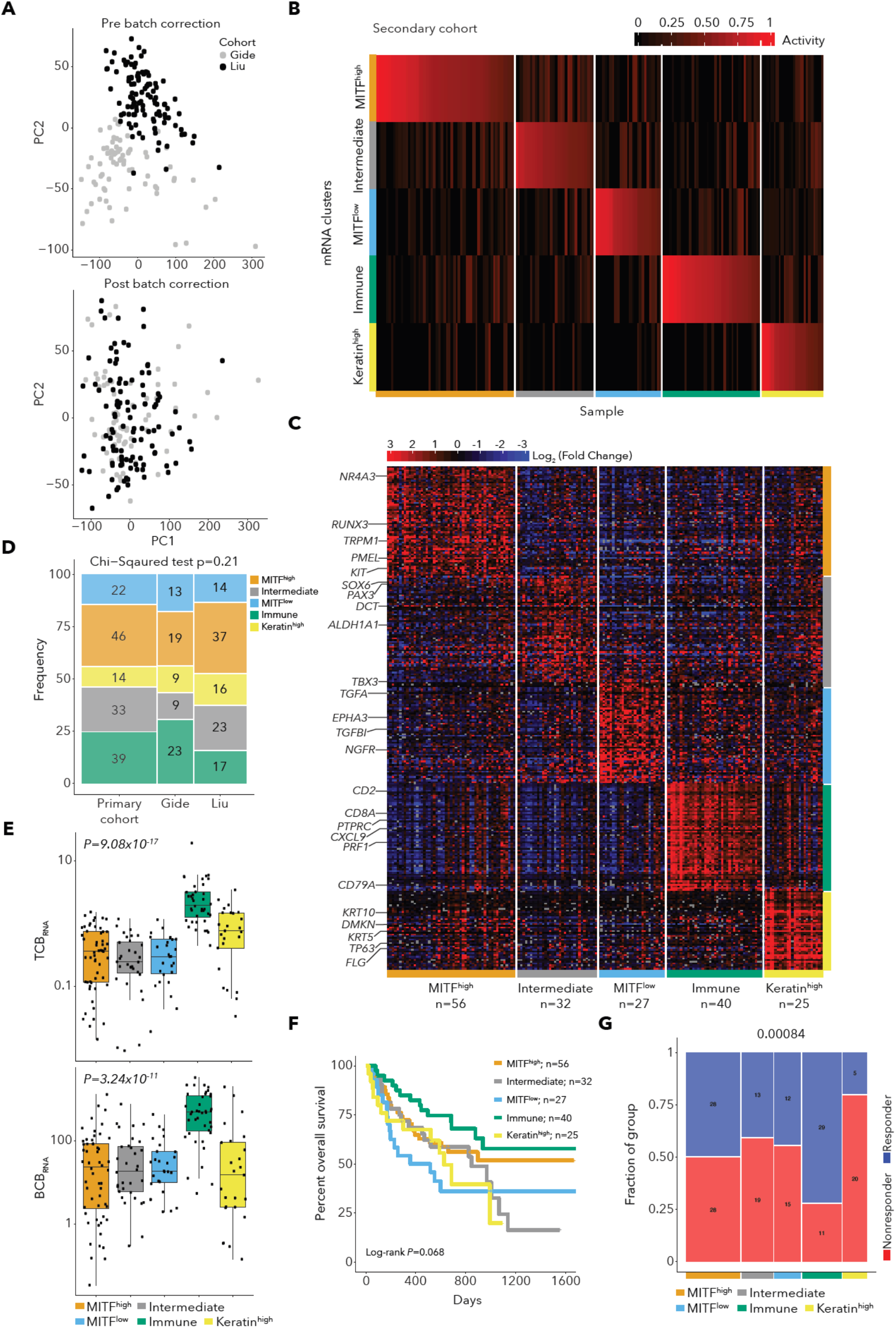
Batch effects correction and melanoma subtyping for the secondary cohort. **A.** PCA of secondary cohort (Gide and Liu cohorts) protein coding gene log2(TPM+1) values before (upper plot) and after (lower plot) batch-effects correction with ComBat^88^. Before batch effects correction, samples cluster by cohort. **B.** NMF H matrix for secondary cohort sample subtyping using subtypes and marker genes identified in TCGA melanoma samples. **C.** Heatmap of marker gene expression for samples in the secondary cohort with samples grouped by subtype. **D.** Comparison of frequency of subtypes in the primary cohort and the Gide and Liu cohorts. **E.** TCBRNA and BCBRNA by subtype for secondary cohort samples with Kruskal-Wallis P values for associations with subtype. **F.** Kaplan-Meier survival curve by subtype for patients in the secondary cohort. **G.** Number of responder and non-responders by subtype in the secondary cohort with Fisher’s exact test P value.

**Supplementary Figure 24.**
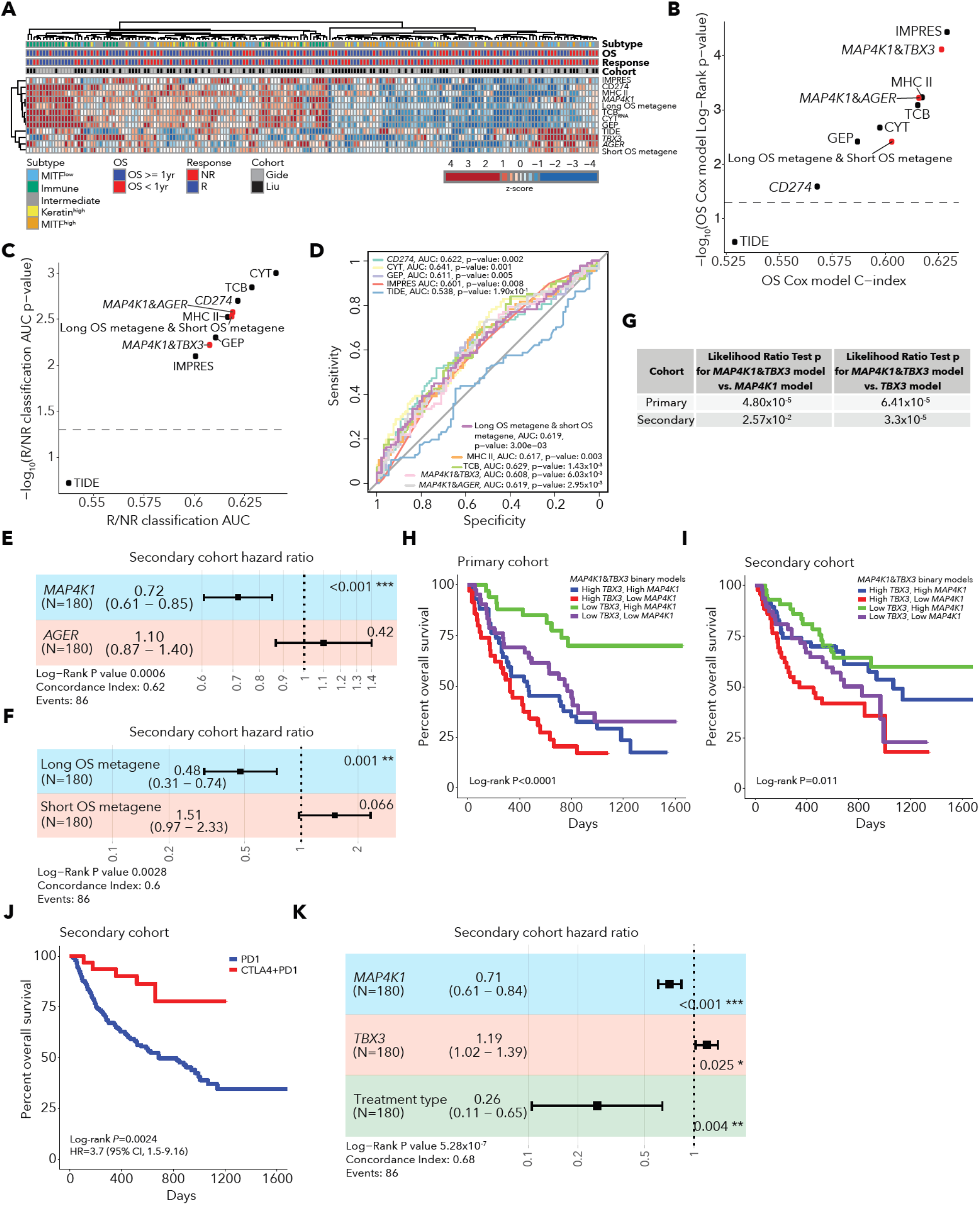
Performance of top gene pair models in the secondary cohort. **A.** Heatmap of z-scored values for immunotherapy predictive models and top gene pairs in the secondary cohort. **B.** Performance of pairwise gene models in comparison to previous immunotherapy predictive models in significance and effect size of predictions of survival in the secondary cohort. **C.** Performance of pairwise gene models in comparison to previous immunotherapy predictive models in significance and effect size of predictions of response in the secondary cohort. **D.** ROC curve of pairwise gene models and previous immunotherapy models in classification of response in the secondary cohort. **E.** Forest plot for *MAP4K1*&*AGER* gene pair Cox survival model in the secondary cohort. Error bars represent 95% confidence intervals for Cox model hazard ratio estimates. **F.** Forest plot for the metagene pair Cox survival model in the secondary cohort. Error bars represent 95% confidence intervals for Cox model hazard ratio estimates. **G.** Likelihood ratio test results comparing Cox survival models with *MAP4K1*&*TBX3* to Cox survival models with *MAP4K1* or *TBX3* alone in the primary and secondary cohorts. In both cohorts, the gene pair models outperformed both single gene models. **H.** Kaplan-Meier survival curve for groups based on binary *MAP4K1* and *TBX3* expression (above or below median) in the primary cohort. **I.** Kaplan-Meier survival curve for groups based on binary *MAP4K1* and *TBX3* expression (above or below median) in the secondary cohort. **J.** Kaplan-Meier survival curve for groups based on treatment (PD-1 alone or combination CTLA-4/PD-1 therapy) in the secondary cohort. **K.** Forest plot for Cox survival model incorporating *MAP4K1* expression, *TBX3* expression and treatment (PD-1 alone or combination CTLA-4/PD-1) in the secondary cohort. After including treatment in the model, *MAP4K1* and *TBX3* expression both remain significant. Error bars represent 95% confidence intervals for Cox model hazard ratio estimates.

**Supplementary Figure 25.**
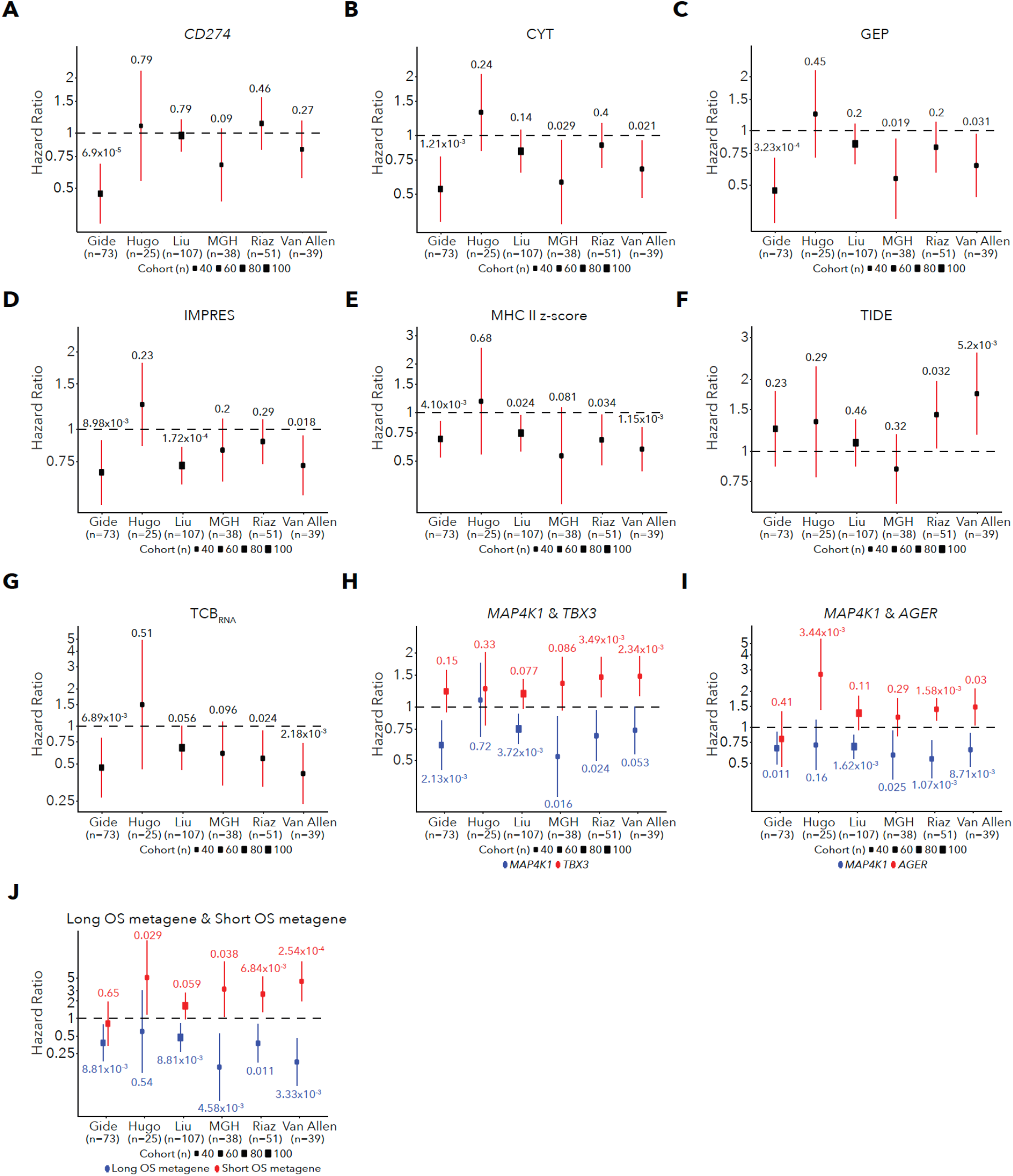
Performance of all models within each cohort separately. **A-G.** Performance of univariate Cox models within each cohort separately. Hazard ratios and error bars representing 95% confidence intervals of Hazard ratio estimates are plotted and Wald test P values are indicated. (A) *CD274* **(B)** CYT **(C)** GEP (D) IMPRES (E) MHC II (F) TIDE (G) TCBRNA. H-J. Performance of Cox models using gene pairs within each cohort. Hazard ratios and confidence intervals are colored by the gene or metagene and the Wald test P values are indicated. (H) *MAP4K1*&*TBX3* (I) *MAP4K1*&*AGER* (J) metagene pair model.

**Supplementary Figure 26.**
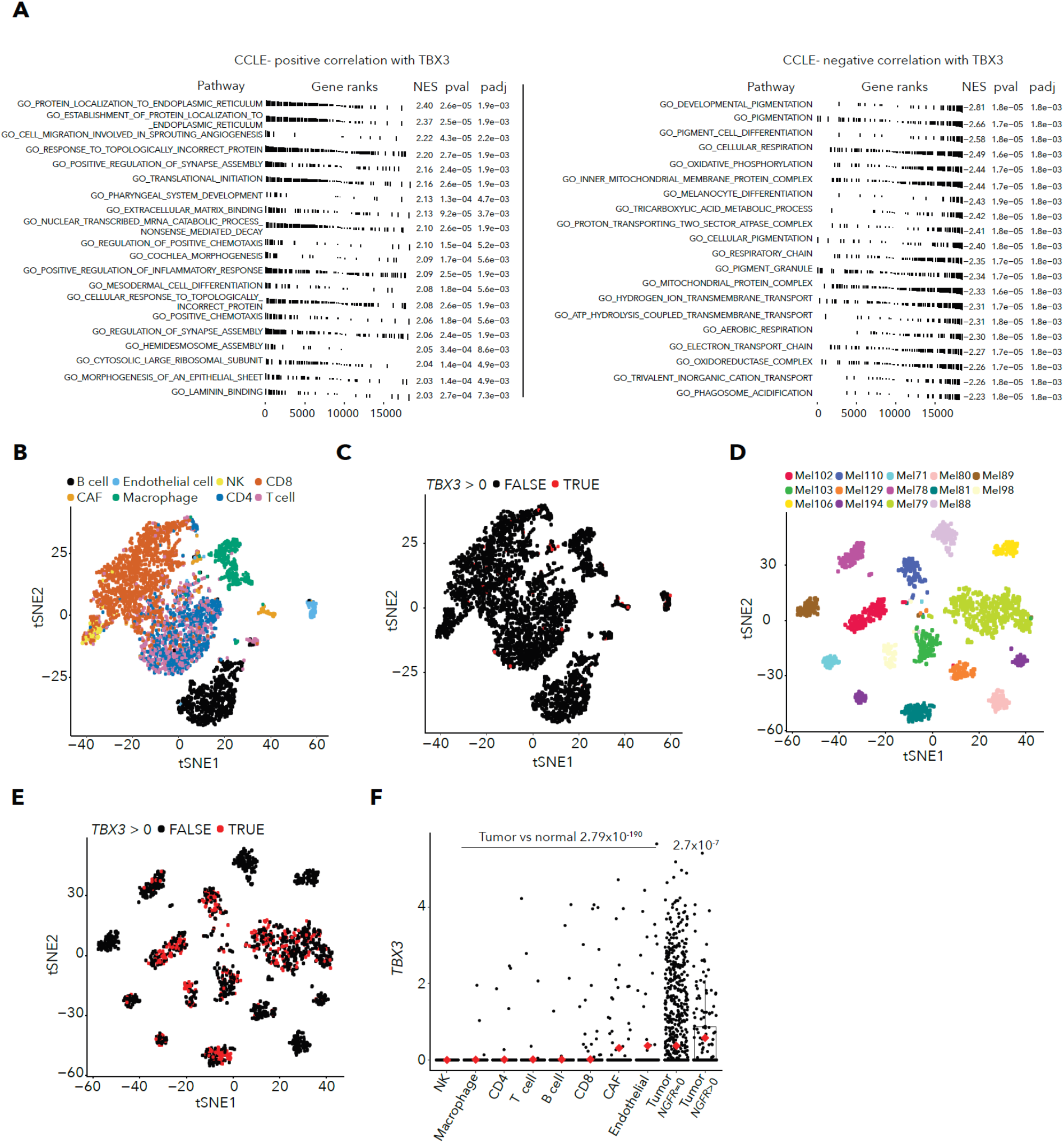
Analysis of melanoma *TBX3* expression. **A.** GSEA for genes ordered by spearman correlation of gene expression with *TBX3* gene expression in CCLE melanoma cell lines using GO terms. Top GSEA results for genes positively correlated with *TBX3* (left), and top GSEA results for genes negatively correlated with *TBX3* (right). GO terms associated with genes negatively correlated with *TBX3* include pigmentation and melanocyte gene sets. B-C. tSNE plot of scRNA data from immune cells with cells labelled by lineage **(B)** or *TBX3* expression status **(C)**. *TBX3* is rarely expressed in any immune cell type. D-E. tSNE plot of scRNA data from melanoma tumor cells with cells labelled by patient (D) or *TBX3* expression status (E). *TBX3* is expressed in tumor cells from some patients. **F.** *TBX3* expression in scRNA data by immune cell type or tumor cell type. Melanoma tumor cells are split between cells with no *NGFR* expression or cells with non-zero *NGFR* expression. Mean expression by group is indicated by red dots. P values for Wilcox tests of *TBX3* expression in tumor single cells vs. normal single cells and tumor *NGFR*>0 cells vs. tumor *NGFR*=0 cells are indicated above.

## References

1. Le, D. T. et al. PD-1 Blockade in Tumors with Mismatch-Repair Deficiency. N. Engl. J. Med. 372, 2509–2520 (2015).

2. Samstein, R. M. et al. Tumor mutational load predicts survival after immunotherapy across multiple cancer types. Nat. Genet. 51, 202–206 (2019).

3. Miao, D. et al. Genomic correlates of response to immune checkpoint blockade in microsatellite-stable solid tumors. Nat. Genet. 50, 1271–1281 (2018).

4. Liu, D. et al. Integrative molecular and clinical modeling of clinical outcomes to PD1 blockade in patients with metastatic melanoma. Nat. Med. 25, 1916–1927 (2019).

5. Tumeh, P. C. et al. PD-1 blockade induces responses by inhibiting adaptive immune resistance. Nature 515, 568–571 (2014).

6. Riaz, N. et al. Tumor and Microenvironment Evolution during Immunotherapy with Nivolumab. Cell 171, 934–949.e16 (2017).

7. Sade-Feldman, M. et al. Defining T Cell States Associated with Response to Checkpoint Immunotherapy in Melanoma. Cell 175, 998–1013.e20 (2018).

8. Keenan, T. E., Burke, K. P. & Van Allen, E. M. Genomic correlates of response to immune checkpoint blockade. Nat. Med. 25, 389–402 (2019).

9. Havel, J. J., Chowell, D. & Chan, T. A. The evolving landscape of biomarkers for checkpoint inhibitor immunotherapy. Nat. Rev. Cancer 19, 133–150 (2019).

10. Herbst, R. S. et al. Predictive correlates of response to the anti-PD-L1 antibody MPDL3280A in cancer patients. Nature 515, 563–567 (2014).

11. Sade-Feldman, M. et al. Resistance to checkpoint blockade therapy through inactivation of antigen presentation. Nat. Commun. 8, 1136 (2017).

12. Zaretsky, J. M. et al. Mutations Associated with Acquired Resistance to PD-1 Blockade in Melanoma. N. Engl. J. Med. 375, 819–829 (2016).

13. Miao, D. et al. Genomic correlates of response to immune checkpoint therapies in clear cell renal cell carcinoma. Science 359, 801–806 (2018).

14. George, S. et al. Loss of PTEN Is Associated with Resistance to Anti-PD-1 Checkpoint Blockade Therapy in Metastatic Uterine Leiomyosarcoma. Immunity 46, 197–204 (2017).

15. Riaz, N. et al. Recurrent SERPINB3 and SERPINB4 mutations in patients who respond to anti-CTLA4 immunotherapy. Nat. Genet. 48, 1327–1329 (2016).

16. Snyder, A. et al. Genetic basis for clinical response to CTLA-4 blockade in melanoma. N. Engl. J. Med. 371, 2189–2199 (2014).

17. Van Allen, E. M. et al. Genomic correlates of response to CTLA-4 blockade in metastatic melanoma. Science 350, 207–211 (2015).

18. Valero, C. et al. The association between tumor mutational burden and prognosis is dependent on treatment context. Nat. Genet. 53, 11–15 (2021).

19. Gurjao, C., Tsukrov, D., Imakaev, M., Luquette, L. J. & Mirny, L. A. Limited evidence of tumour mutational burden as a biomarker of response to immunotherapy. bioRxiv 2020.09.03.260265 (2020) doi:10.1101/2020.09.03.260265.

20. Litchfield, K. et al. Meta-analysis of tumor- and T cell-intrinsic mechanisms of sensitization to checkpoint inhibition. Cell 184, 596–614.e14 (2021).

21. Ayers, M. et al. IFN-γ-related mRNA profile predicts clinical response to PD-1 blockade. J. Clin. Invest. 127, 2930–2940 (2017).

22. Cristescu, R. et al. Pan-tumor genomic biomarkers for PD-1 checkpoint blockade-based immunotherapy. Science 362, (2018).

23. Chowell, D. et al. Patient HLA class I genotype influences cancer response to checkpoint blockade immunotherapy. Science 359, 582–587 (2018).

24. Łuksza, M. et al. A neoantigen fitness model predicts tumour response to checkpoint blockade immunotherapy. Nature 551, 517–520 (2017).

25. Shin, D. S. et al. Primary Resistance to PD-1 Blockade Mediated by JAK1/2 Mutations. Cancer Discov. 7, 188–201 (2017).

26. Mehta, A. et al. Immunotherapy Resistance by Inflammation-Induced Dedifferentiation. Cancer Discov. 8, 935–943 (2018).

27. Rodig, S. J. et al. MHC proteins confer differential sensitivity to CTLA-4 and PD-1 blockade in untreated metastatic melanoma. Sci. Transl. Med. 10, (2018).

28. Gao, J. et al. Loss of IFN-γ Pathway Genes in Tumor Cells as a Mechanism of Resistance to Anti-CTLA-4 Therapy. Cell 167, 397–404.e9 (2016).

29. Thommen, D. S. et al. A transcriptionally and functionally distinct PD-1+ CD8+ T cell pool with predictive potential in non-small-cell lung cancer treated with PD-1 blockade. Nat. Med. 24, 994–1004 (2018).

30. Huang, A. C. et al. T-cell invigoration to tumour burden ratio associated with anti-PD-1 response. Nature 545, 60–65 (2017).

31. Petitprez, F. et al. B cells are associated with survival and immunotherapy response in sarcoma. Nature 577, 556–560 (2020).

32. Cabrita, R. et al. Tertiary lymphoid structures improve immunotherapy and survival in melanoma. Nature 577, 561–565 (2020).

33. Wu, T. D. et al. Peripheral T cell expansion predicts tumour infiltration and clinical response. Nature 579, 274–278 (2020).

34. Gandara, D. R. et al. Blood-based tumor mutational burden as a predictor of clinical benefit in non-small-cell lung cancer patients treated with atezolizumab. Nat. Med. 24, 1441–1448 (2018).

35. Hellmann, M. D. et al. Genomic Features of Response to Combination Immunotherapy in Patients with Advanced Non-Small-Cell Lung Cancer. Cancer Cell 33, 843–852.e4 (2018).

36. Roh, W. et al. Integrated molecular analysis of tumor biopsies on sequential CTLA-4 and PD-1 blockade reveals markers of response and resistance. Sci. Transl. Med. 9, (2017).

37. Hugo, W. et al. Genomic and Transcriptomic Features of Response to Anti-PD-1 Therapy in Metastatic Melanoma. Cell vol. 165 35–44 (2016).

38. . Cancer Genome Atlas Network. Genomic Classification of Cutaneous Melanoma. Cell 161, 1681–1696 (2015).

39. Davoli, T., Uno, H., Wooten, E. C. & Elledge, S. J. Tumor aneuploidy correlates with markers of immune evasion and with reduced response to immunotherapy. Science 355, (2017).

40. Helmink, B. A. et al. B cells and tertiary lymphoid structures promote immunotherapy response. Nature 577, 549–555 (2020).

41. Chen, P.-L. et al. Analysis of Immune Signatures in Longitudinal Tumor Samples Yields Insight into Biomarkers of Response and Mechanisms of Resistance to Immune Checkpoint Blockade. Cancer Discov. 6, 827–837 (2016).

42. Mariathasan, S. et al. TGFβ attenuates tumour response to PD-L1 blockade by contributing to exclusion of T cells. Nature 554, 544–548 (2018).

43. Kim, S. T. et al. Comprehensive molecular characterization of clinical responses to PD-1 inhibition in metastatic gastric cancer. Nat. Med. 24, 1449–1458 (2018).

44. 44. Kim, J. et al. The Cancer Genome Atlas Expression Subtypes Stratify Response to Checkpoint Inhibition in Advanced Urothelial Cancer and Identify a Subset of Patients with High Survival Probability. Eur. Urol. (2019).

45. Robertson, A. G. et al. Comprehensive Molecular Characterization of Muscle-Invasive Bladder Cancer. Cell 174, 1033 (2018).

46. Tsoi, J. et al. Multi-stage Differentiation Defines Melanoma Subtypes with Differential Vulnerability to Drug-Induced Iron-Dependent Oxidative Stress. Cancer Cell 33, 890– 904.e5 (2018).

47. Konieczkowski, D. J. et al. A melanoma cell state distinction influences sensitivity to MAPK pathway inhibitors. Cancer Discov. 4, 816–827 (2014).

48. Tirosh, I. et al. Dissecting the multicellular ecosystem of metastatic melanoma by single-cell RNA-seq. Science 352, 189–196 (2016).

49. Hernandez, S. et al. The Kinase Activity of Hematopoietic Progenitor Kinase 1 Is Essential for the Regulation of T Cell Function. Cell Rep. 25, 80–94 (2018).

50. Liu, J. et al. Critical role of kinase activity of hematopoietic progenitor kinase 1 in anti-tumor immune surveillance. PLoS One 14, e0212670 (2019).

51. Auslander, N. et al. Robust prediction of response to immune checkpoint blockade therapy in metastatic melanoma. Nat. Med. 24, 1545–1549 (2018).

52. Jiang, P. et al. Signatures of T cell dysfunction and exclusion predict cancer immunotherapy response. Nat. Med. 24, 1550–1558 (2018).

53. Rooney, M. S., Shukla, S. A., Wu, C. J., Getz, G. & Hacohen, N. Molecular and genetic properties of tumors associated with local immune cytolytic activity. Cell 160, 48–61 (2015).

54. Gide, T. N. et al. Distinct Immune Cell Populations Define Response to Anti-PD-1 Monotherapy and Anti-PD-1/Anti-CTLA-4 Combined Therapy. Cancer Cell 35, 238–255.e6 (2019).

55. Barretina, J. et al. The Cancer Cell Line Encyclopedia enables predictive modelling of anticancer drug sensitivity. Nature vol. 483 603–307 (2012).

56. Jerby-Arnon, L. et al. A Cancer Cell Program Promotes T Cell Exclusion and Resistance to Checkpoint Blockade. Cell 175, 984–997.e24 (2018).

57. Boshuizen, J. et al. Reversal of pre-existing NGFR-driven tumor and immune therapy resistance. Nat. Commun. 11, 3946 (2020).

58. Peres, J. & Prince, S. The T-box transcription factor, TBX3, is sufficient to promote melanoma formation and invasion. Mol. Cancer 12, 117 (2013).

59. Li, H. & Durbin, R. Fast and accurate short read alignment with Burrows–Wheeler transform. Bioinformatics 25, 1754–1760 (2009).

60. Birger, C. et al. FireCloud, a scalable cloud-based platform for collaborative genome analysis: Strategies for reducing and controlling costs. doi:10.1101/209494.

61. Cibulskis, K. et al. ContEst: estimating cross-contamination of human samples in next- generation sequencing data. Bioinformatics 27, 2601–2602 (2011).

62. Cibulskis, K. et al. Sensitive detection of somatic point mutations in impure and heterogeneous cancer samples. Nat. Biotechnol. 31, 213–219 (2013).

63. Saunders, C. T. et al. Strelka: accurate somatic small-variant calling from sequenced tumor–normal sample pairs. Bioinformatics 28, 1811–1817 (2012).

64. Taylor-Weiner, A. et al. DeTiN: overcoming tumor-in-normal contamination. Nat. Methods 15, 531–534 (2018).

65. Ramos, A. H. et al. Oncotator: cancer variant annotation tool. Hum. Mutat. 36, E2423–9 (2015).

66. Costello, M. et al. Discovery and characterization of artifactual mutations in deep coverage targeted capture sequencing data due to oxidative DNA damage during sample preparation. Nucleic Acids Res. 41, e67 (2013).

67. Ellrott, K. et al. Scalable Open Science Approach for Mutation Calling of Tumor Exomes Using Multiple Genomic Pipelines. Cell Syst 6, 271–281.e7 (2018).

68. Kent, W. J. BLAT—The BLAST-Like Alignment Tool. Genome Res. 12, 656–664 (2002).

69. Carter, S. L. et al. Absolute quantification of somatic DNA alterations in human cancer. Nature Biotechnology vol. 30 413–421 (2012).

70. Lawrence, M. S. et al. Mutational heterogeneity in cancer and the search for new cancer- associated genes. Nature 499, 214–218 (2013).

71. Lawrence, M. S. et al. Discovery and saturation analysis of cancer genes across 21 tumour types. Nature 505, 495–501 (2014).

72. Kim, J. et al. Somatic ERCC2 mutations are associated with a distinct genomic signature in urothelial tumors. Nat. Genet. 48, 600–606 (2016).

73. Kasar, S. et al. Whole-genome sequencing reveals activation-induced cytidine deaminase signatures during indolent chronic lymphocytic leukaemia evolution. Nature Communications vol. 6 (2015).

74. Tan, V. Y. F. & Fevotte, C. Automatic Relevance Determination in Nonnegative Matrix Factorization with the /spl beta/-Divergence. IEEE Transactions on Pattern Analysis and Machine Intelligence vol. 35 1592–1605 (2013).

75. Alexandrov, L. B. et al. Signatures of mutational processes in human cancer. Nature 500, 415–421 (2013).

76. Alexandrov, L. B., Nik-Zainal, S., Wedge, D. C., Campbell, P. J. & Stratton, M. R. Deciphering signatures of mutational processes operative in human cancer. Cell Rep. 3, 246–259 (2013).

77. Shukla, S. A. et al. Comprehensive analysis of cancer-associated somatic mutations in class I HLA genes. Nat. Biotechnol. 33, 1152–1158 (2015).

78. Hoof, I. et al. NetMHCpan, a method for MHC class i binding prediction beyond humans. Immunogenetics 61, 1–13 (2009).

79. Nielsen, M. & Andreatta, M. NetMHCpan-3.0; improved prediction of binding to MHC class I molecules integrating information from multiple receptor and peptide length datasets. Genome Medicine vol. 8 (2016).

80. Jurtz, V. et al. NetMHCpan-4.0: Improved Peptide-MHC Class I Interaction Predictions Integrating Eluted Ligand and Peptide Binding Affinity Data. J. Immunol. 199, 3360–3368 (2017).

81. McKenna, A. et al. The Genome Analysis Toolkit: a MapReduce framework for analyzing next-generation DNA sequencing data. Genome Res. 20, 1297–1303 (2010).

82. Mermel, C. H. et al. GISTIC2. 0 facilitates sensitive and confident localization of the targets of focal somatic copy-number alteration in human cancers. Genome Biol. 12, R41 (2011).

83. Landau, D. A. et al. Evolution and impact of subclonal mutations in chronic lymphocytic leukemia. Cell 152, 714–726 (2013).

84. 84. Leshchiner, I., Livitz, D., Gainor, J. F., Rosebrock, D. & Spiro, O. Comprehensive analysis of tumour initiation, spatial and temporal progression under multiple lines of treatment. bioRxiv (2019).

85. GTEx Consortium. The GTEx Consortium atlas of genetic regulatory effects across human tissues. Science 369, 1318–1330 (2020).

86. Dobin, A. et al. STAR: ultrafast universal RNA-seq aligner. Bioinformatics 29, 15–21 (2013).

87. Li, B. & Dewey, C. N. RSEM: accurate transcript quantification from RNA-Seq data with or without a reference genome. BMC Bioinformatics 12, 323 (2011).

88. Johnson, W. E., Li, C. & Rabinovic, A. Adjusting batch effects in microarray expression data using empirical Bayes methods. Biostatistics 8, 118–127 (2007).

89. Bolotin, D. A. et al. MiXCR: software for comprehensive adaptive immunity profiling. Nat. Methods 12, 380–381 (2015).

90. Bolotin, D. A. et al. Antigen receptor repertoire profiling from RNA-seq data. Nat. Biotechnol. 35, 908–911 (2017).

91. Love, M. I., Huber, W. & Anders, S. Moderated estimation of fold change and dispersion for RNA-seq data with DESeq2. Genome Biol. 15, 550 (2014).

92. Uhlen, M. et al. A genome-wide transcriptomic analysis of protein-coding genes in human blood cells. Science 366, (2019).

93. Mason, S. J. & Graham, N. E. Areas beneath the relative operating characteristics (ROC) and relative operating levels (ROL) curves: Statistical significance and interpretation. of the Royal Meteorological Society: A … (2002).

94. Korotkevich, G. et al. Fast gene set enrichment analysis. Cold Spring Harbor Laboratory 060012 (2021) doi:10.1101/060012.

